# Deep embedded clustering by relevant scales and genome-wide association study in autism

**DOI:** 10.1101/2022.07.25.500917

**Authors:** Fumihiko Ueno, Tomomi Onuma, Ippei Takahashi, Hisashi Ohseto, Akira Narita, Taku Obara, Mami Ishikuro, Keiko Murakami, Aoi Noda, Fumiko Matsuzaki, Hirohito Metoki, Gen Tamiya, Shigeo Kure, Shinichi Kuriyama

## Abstract

The etiology of autism spectrum disorders (ASD) remains unclear. Stratifying patients with ASD may help to identify genetically homogeneous subgroups. Using a deep embedded clustering algorithm, we conducted cluster analyses of Simons Foundation Powering Autism Research for Knowledge (SPARK) datasets and performed genome-wide association studies (GWAS) of the clusters. We observed no significant associations in the conventional GWAS comparing all patients to all controls. However, in the GWAS, comparing patients divided into clusters with similar phenotypes to controls (cluster-based GWAS), we identified 90 chromosomal loci that satisfied the *P* < 5.0 × 10^−8^, several of which were located within or near previously reported candidate genes for ASD. Our findings suggest that clustering may successfully identify subgroups with relatively homogeneous disease etiologies.

---

Autism spectrum disorder (ASD) is a neurodevelopmental disorder primarily characterized by difficulties in communication and repetitive behaviors (*1*). Despite efforts to understand the molecular mechanisms underlying ASD, its etiology remains unclear (*2*). Evidence suggests that genetic factors strongly contribute to the risk of ASD development (*3*). For instance, identical twins have a higher concordance rate of ASD (92%) than dizygotic twins (10%) (*4*); the risk ratio for ASD recurrence between siblings is 22 (*5*).

Results from previous genome-wide association studies (GWAS) have identified numerous genetic variants associated with ASD (*6, 7*). The association of these genetic variants with a single disease may be explained by a polygenic model in which the effects of each genetic variant are weak, yet contribute to the disease onset (*8, 9*). When performing GWAS for a disease, the larger the sample size, the more signals are generally identified, while as the sample size decreases, so does the identified signal (Supplementary Figure S1). When conducting GWAS with a certain sample size where no significant signal is produced, the signal will become increasingly difficult to identify if the sample size is reduced by dividing the patients. However, if ASD includes a subtype in which a small number of genes have a relatively strong influence, a subtype described by the oligogenic model, then it may be possible to identify some signals by dividing patients into phenotypically similar clusters and exploring genetic factors. A simulation study has suggested that dividing patients into more homogeneous populations increases the power of GWAS, regardless of the sample size (*10*).

The purpose of this study is to clarify the genetic architecture of ASD by dividing patients into clusters using phenotypic variables and performing GWAS (cluster-based GWAS: cGWAS). We have previously suggested that ASD is an aggregation of etiologically heterogeneous subtypes, and within that aggregation are subtypes whose genetic architecture is described by the oligogenic model (*11*), by performing cGWAS with ASD using different datasets and k-means algorithm (*12, 13*). Herein, we increased the sample size (*14*) and introduced deep embedded clustering (DEC) algorithm (*15, 16*). If it can be demonstrated that ASD is an aggregation of etiologically heterogeneous subtypes through this study, it can be expected to lead to precision medicine that is tailored to the characteristics of each subtype.

## Results

### Cluster-based genome-wide association study

In a preliminary study, we conducted a conventional GWAS comparing all patients vs. all controls with the Simons Foundation Powering Autism Research for Knowledge (SPARK) WES1(27K) genotypic dataset and observed no significant associations (Figure 1).

**Figure 1.**
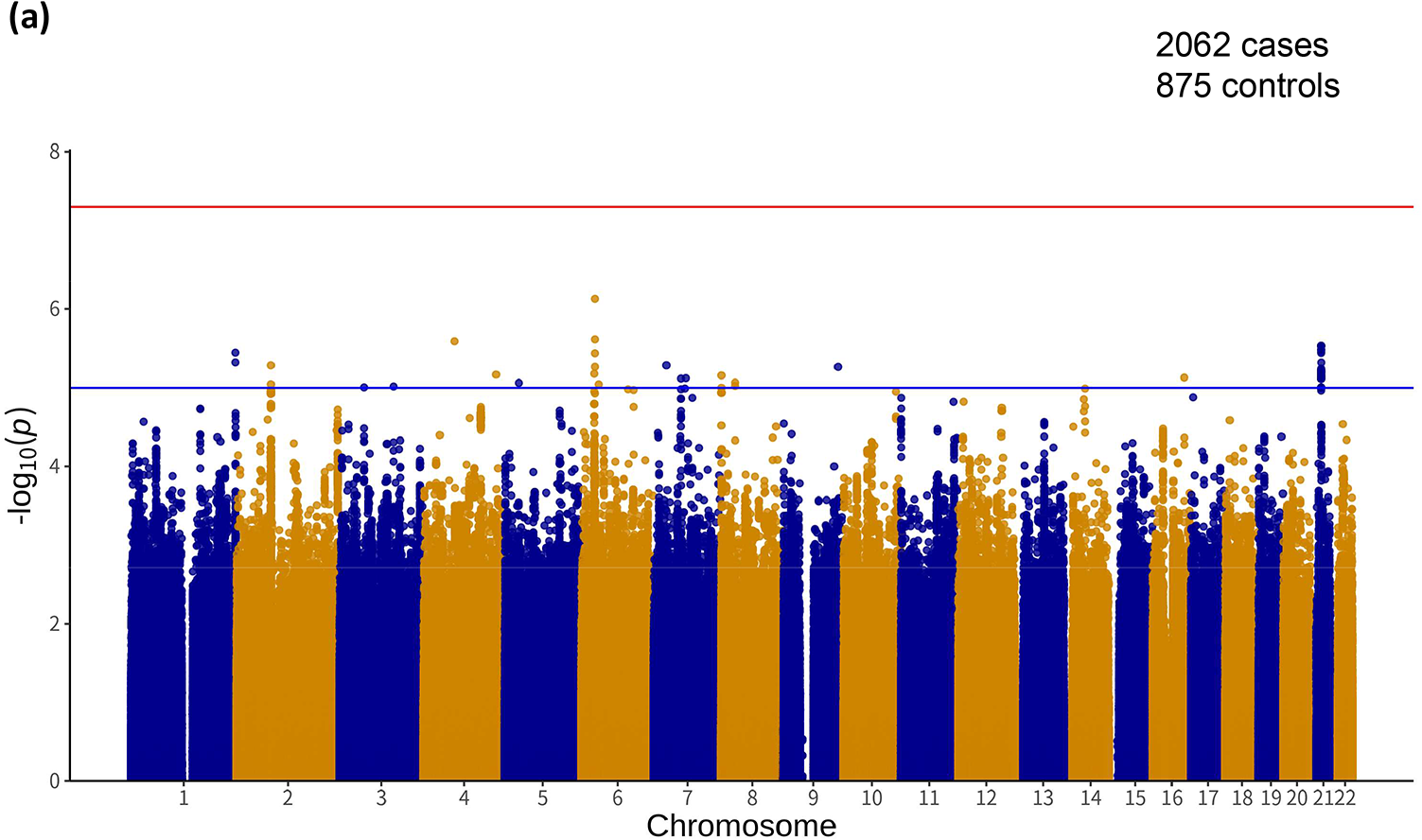
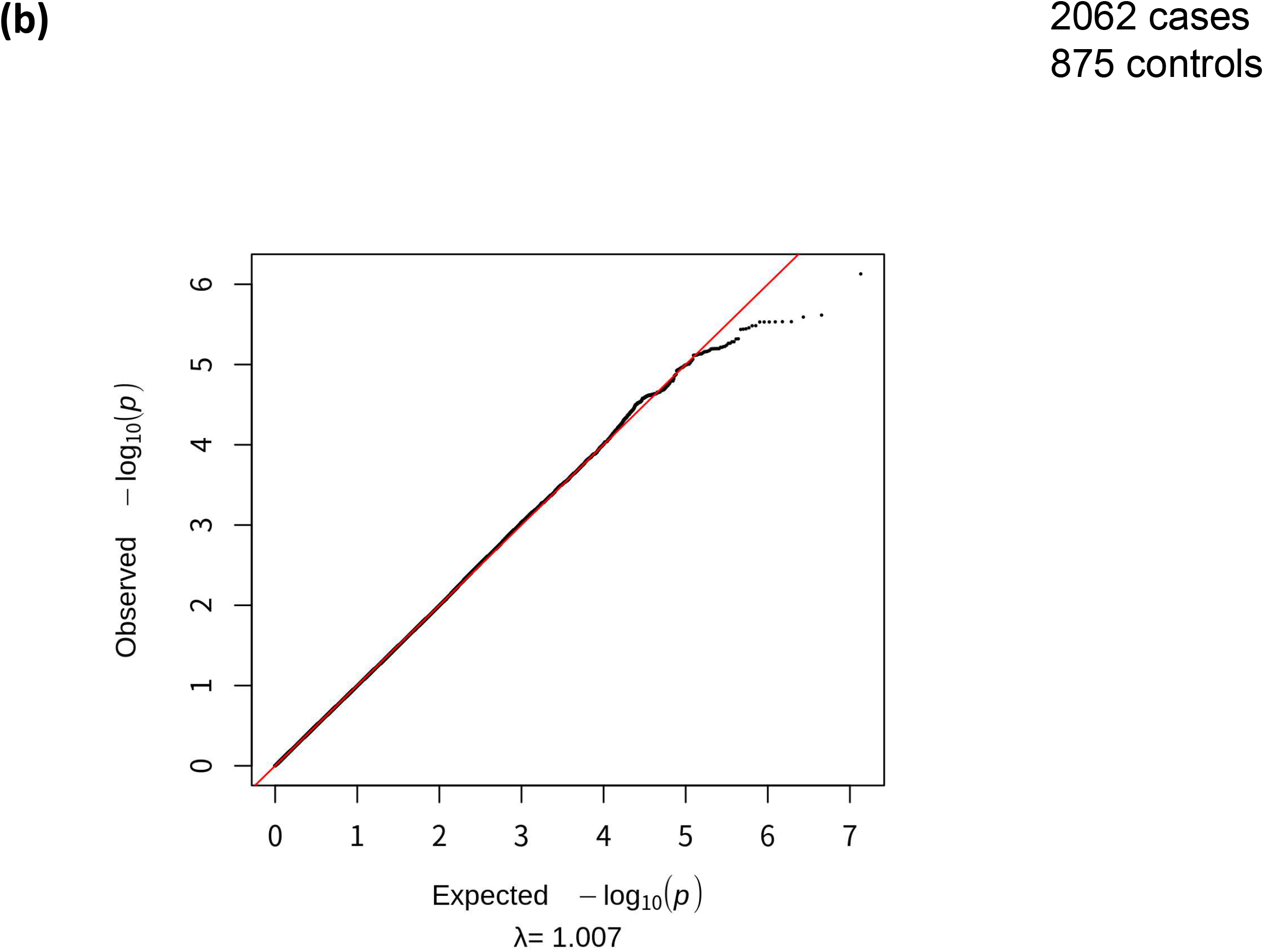
Manhattan plots (a) and corresponding quantile-quantile plots (b) in GWAS for all male probands vs unrelated controls. We conducted a GWAS in the SPARK dataset of 2,062 male probands and 875 unrelated controls genotyped by the Illumina GSA v1.0 array using the logistic regression model. We observed no significant associations in this GWAS with the genome-wide threshold of *P* < 5.0 × 10^−8^.

The DEC algorithm requires a cluster number (k) determined by researchers. The average inflation factor λ for a cGWAS with k = 40 was 0.9. Regarding an appropriate threshold of λ, empirically, a value <1.05 is deemed safe for avoiding false positives (*17*). Therefore, we regarded the condition of λ < 1.05 as one of the indicators of successful clustering. We, therefore, considered a cGWAS using cluster analysis with k = 40 as the most appropriate approach to the present dataset. Fixing hyperparameter k of DEC at 40 eventually led to the division of the dataset into 39 clusters.

The characteristics of each cluster are presented in Table 1. The heat map shows the characteristics of the clusters. For example, cluster 5 has a relatively high Developmental Coordination Disorder Questionnaire (DCDQ) (*18*) and Repetitive Behaviors Scale-Revised (RBS-R) (*19*), average Social Communication Questionnaire (SCQ) (*20*), no underlying disease, and a high rate of interventional treatment.

**Table 1.** Characteristics of clusters.

| Cluster no. | 1 | 2 | 3 | 4 | 5 | 6 | 7 | 8 | 9 | 10 | 11 | 12 | 13 | 14 | 15 | 16 | 17 | 18 | 19 | 20 | 21 | 22 | 23 | 24 | 25 | 26 | 27 | 28 | 29 | 30 | 31 | 32 | 33 | 34 | 35 | 36 | 37 | 38 | 39 |
| --- | --- | --- | --- | --- | --- | --- | --- | --- | --- | --- | --- | --- | --- | --- | --- | --- | --- | --- | --- | --- | --- | --- | --- | --- | --- | --- | --- | --- | --- | --- | --- | --- | --- | --- | --- | --- | --- | --- | --- |
| n | 195 | 6 | 16 | 2 | 80 | 262 | 21 | 38 | 11 | 6 | 195 | 10 | 168 | 32 | 193 | 52 | 63 | 11 | 11 | 126 | 14 | 17 | 14 | 11 | 36 | 2 | 1 | 19 | 4 | 39 | 12 | 70 | 5 | 91 | 62 | 83 | 25 | 32 | 27 |
| DCDQ, mean |  |  |  |  |  |  |  |  |  |  |  |  |  |  |  |  |  |  |  |  |  |  |  |  |  |  |  |  |  |  |  |  |  |  |  |  |  |  |  |
| Control during movement subscale score | 16.3 | 17.7 | 15.8 | 16.5 | 19.2 | 24.1 | 14.5 | 15.6 | 17.8 | 20.5 | 12.6 | 12.7 | 18.3 | 18.1 | 10.8 | 13.2 | 13.3 | 20.6 | 18.5 | 12.3 | 16.4 | 15.7 | 16.9 | 20 | 20.3 | 10.5 | 16 | 12.1 | 20.2 | 17.4 | 14 | 15.6 | 21.8 | 18.3 | 17.6 | 15.8 | 14.6 | 18.2 | 17.4 |
| Fine motor/handwriting subscale score | 8.9 | 10.3 | 8.6 | 5 | 11.7 | 15.1 | 10.1 | 6.7 | 8.5 | 9 | 5.5 | 8.6 | 11.8 | 11.4 | 5.8 | 7.6 | 7.5 | 11.5 | 10.3 | 8.4 | 10.1 | 7.6 | 8.7 | 10.5 | 12.9 | 8 | 12 | 9.4 | 9.5 | 8.5 | 7.1 | 8.2 | 12.4 | 11.2 | 11.4 | 8.2 | 7.4 | 7.4 | 8.8 |
| General Coordination subscale score | 11.2 | 11 | 11.1 | 12 | 11.4 | 16.8 | 9.1 | 8.4 | 10.6 | 11.2 | 9.7 | 8.2 | 12.2 | 13.4 | 7.8 | 8 | 8.2 | 10.1 | 13.3 | 7.6 | 9.4 | 11.6 | 11 | 14.5 | 12.2 | 7 | 11 | 7.9 | 12.8 | 10 | 10.8 | 10 | 14.8 | 11.9 | 10.5 | 8 | 8.4 | 11.5 | 9.7 |
| Total score | 36.5 | 39 | 35.5 | 33.5 | 42.5 | 56 | 33.5 | 30.6 | 36.9 | 40.8 | 27.8 | 29.5 | 42.4 | 42.9 | 24.3 | 28.8 | 28.7 | 42.3 | 42.4 | 28.2 | 35.9 | 34.9 | 36.6 | 45 | 45.5 | 25.5 | 39 | 29.4 | 42.5 | 35.9 | 32.5 | 33.8 | 49 | 41.3 | 39.6 | 32 | 30.5 | 37.5 | 36.6 |
| RBS-R, mean |  |  |  |  |  |  |  |  |  |  |  |  |  |  |  |  |  |  |  |  |  |  |  |  |  |  |  |  |  |  |  |  |  |  |  |  |  |  |  |
| Stereotyped behavior subscale score | 10.1 | 5 | 6.6 | 3.5 | 7.7 | 3 | 4.9 | 6.5 | 6.3 | 4.7 | 6.1 | 5.5 | 2.9 | 4 | 8.8 | 2.3 | 2.9 | 5.9 | 4 | 6.3 | 5.6 | 7.4 | 5.3 | 4.9 | 3.8 | 6.5 | 10 | 5.1 | 6.2 | 4.9 | 7.2 | 5.1 | 4.2 | 5.4 | 6.3 | 4.8 | 3.7 | 7.4 | 5.3 |
| Self-injurious behavior subscale score | 8.6 | 6.2 | 0.6 | 0 | 6.4 | 1.3 | 1.7 | 6.7 | 0.5 | 1.7 | 2.7 | 5.1 | 1.5 | 1 | 5.1 | 2.3 | 1.5 | 4.5 | 1.5 | 4.4 | 1.5 | 2.4 | 1.4 | 0.8 | 1.9 | 1 | 4 | 1.4 | 3.5 | 1.8 | 5.6 | 2.8 | 3.2 | 1.9 | 3 | 4.7 | 2.7 | 2.4 | 3.9 |
| Compulsive behavior subscale score | 11.3 | 4.2 | 4.6 | 2 | 7.6 | 2.4 | 4.9 | 6.1 | 8.3 | 4.2 | 3.4 | 2.8 | 2.3 | 5.4 | 9.2 | 2 | 4.1 | 4.4 | 5.6 | 6.4 | 3.9 | 6.5 | 5.6 | 5.4 | 3.2 | 8.5 | 0 | 3.1 | 3.5 | 4.6 | 6.4 | 3 | 5.2 | 4.5 | 6.1 | 3.3 | 3.5 | 8.6 | 4.3 |
| Ritualistic behavior subscale score | 10.8 | 6 | 6.9 | 1.5 | 9 | 2.9 | 5 | 8.8 | 7.5 | 8 | 2.6 | 6.1 | 3 | 6.3 | 9.5 | 3.3 | 6.3 | 6.5 | 6 | 8 | 6.1 | 9.8 | 9.1 | 6.5 | 4.4 | 8 | 6 | 6 | 5.5 | 5.8 | 7 | 3.6 | 8.8 | 5 | 7.2 | 4.7 | 6.4 | 9.1 | 6 |
| Sameness behavior subscale score | 18.2 | 7.7 | 7.5 | 3.5 | 15.8 | 4 | 7.2 | 14.2 | 10.2 | 10.2 | 4.7 | 7.8 | 4.1 | 9.1 | 16.7 | 5.8 | 9.3 | 10.2 | 9.4 | 12.3 | 7.9 | 12.5 | 9.9 | 8.2 | 7.1 | 11.5 | 13 | 5.7 | 7.2 | 7.7 | 11.9 | 5.1 | 11.8 | 6.8 | 11.4 | 7.7 | 7.5 | 13.9 | 8.9 |
| Restricted behavior subscale score | 7.5 | 3.8 | 3.9 | 2.5 | 6.2 | 2 | 3.7 | 6.7 | 6.3 | 3.2 | 3.8 | 3.8 | 2 | 4.3 | 6.7 | 2.8 | 3.1 | 4.7 | 3.5 | 4.8 | 2.5 | 3.4 | 3.6 | 4.7 | 3.1 | 5.5 | 6 | 2.9 | 5 | 4.7 | 5.7 | 4 | 4.4 | 3.9 | 4.6 | 3.4 | 3.4 | 7.2 | 5.6 |
| Total score | 66.6 | 32.8 | 30.1 | 13 | 52.7 | 15.7 | 27.5 | 49 | 39 | 31.8 | 23.4 | 31.1 | 15.7 | 30.2 | 56 | 18.4 | 27.2 | 36.1 | 30 | 42.2 | 27.6 | 41.9 | 34.9 | 30.5 | 23.7 | 41 | 39 | 24.1 | 31 | 29.6 | 43.8 | 23.6 | 37.6 | 27.5 | 38.7 | 28.6 | 27.2 | 48.6 | 33.8 |
| SCQ, mean |  |  |  |  |  |  |  |  |  |  |  |  |  |  |  |  |  |  |  |  |  |  |  |  |  |  |  |  |  |  |  |  |  |  |  |  |  |  |  |
| Total score | 27 | 22.8 | 20 | 24 | 23.9 | 17.8 | 22.5 | 27.6 | 24.4 | 27.7 | 23.6 | 27 | 14.1 | 18.6 | 29.6 | 21.7 | 19.7 | 22.5 | 24.3 | 27.1 | 26.1 | 22.6 | 21.8 | 26.5 | 26.6 | 25 | 23 | 16.5 | 32 | 24.2 | 23.3 | 23.3 | 26.4 | 19.7 | 18.7 | 25 | 15.8 | 24.8 | 20.8 |
| Basic screening, % |  |  |  |  |  |  |  |  |  |  |  |  |  |  |  |  |  |  |  |  |  |  |  |  |  |  |  |  |  |  |  |  |  |  |  |  |  |  |  |
| Premature birth | 12.8 | 33.3 | 0 | 0 | 10 | 8 | 4.8 | 13.5 | 0 | 0 | 9.7 | 50 | 6.5 | 25 | 18.2 | 3.9 | 17.7 | 0 | 9.1 | 22.2 | 7.1 | 17.6 | 21.4 | 9.1 | 27.8 | 0 | 0 | 15.8 | 50 | 15.4 | 0 | 5.7 | 0 | 7.7 | 6.5 | 10.8 | 12 | 21.9 | 33.3 |
| Macrocephaly | 6.7 | 0 | 0 | 0 | 0 | 2.3 | 0 | 5.4 | 9.1 | 0 | 7.2 | 0 | 1.2 | 0 | 7.3 | 5.9 | 8.1 | 0 | 0 | 3.2 | 0 | 0 | 0 | 0 | 2.8 | 0 | 0 | 5.3 | 0 | 5.1 | 8.3 | 4.3 | 0 | 1.1 | 0 | 2.4 | 0 | 3.1 | 3.7 |
| Seizure disorder or epilepsy | 6.2 | 16.7 | 0 | 0 | 1.2 | 3.4 | 0 | 10.8 | 0 | 0 | 7.2 | 20 | 2.4 | 0 | 12 | 2 | 3.2 | 9.1 | 9.1 | 7.1 | 7.1 | 0 | 0 | 0 | 5.6 | 50 | 0 | 0 | 0 | 2.6 | 0 | 4.3 | 0 | 1.1 | 1.6 | 4.8 | 4 | 9.4 | 11.1 |
| Dominant hand, % |  |  |  |  |  |  |  |  |  |  |  |  |  |  |  |  |  |  |  |  |  |  |  |  |  |  |  |  |  |  |  |  |  |  |  |  |  |  |  |
| Either or both | 24.7 | 16.7 | 31.2 | 0 | 12.5 | 9.2 | 4.8 | 28.9 | 18.2 | 0 | 28.1 | 10 | 13.1 | 9.4 | 21.8 | 3.8 | 4.8 | 9.1 | 0 | 15.9 | 0 | 5.9 | 7.1 | 18.2 | 0 | 0 | 0 | 26.3 | 0 | 2.6 | 45.5 | 21.4 | 0 | 17.6 | 17.7 | 7.2 | 16 | 21.9 | 7.4 |
| Left | 11.3 | 0 | 18.8 | 0 | 5 | 9.2 | 0 | 15.8 | 18.2 | 16.7 | 18.2 | 30 | 14.3 | 28.1 | 18.7 | 9.6 | 9.5 | 18.2 | 27.3 | 11.1 | 0 | 11.8 | 7.1 | 9.1 | 5.6 | 50 | 0 | 5.3 | 0 | 20.5 | 36.4 | 15.7 | 0 | 12.1 | 14.5 | 3.6 | 8 | 18.8 | 14.8 |
| Right | 63.9 | 83.3 | 50 | 100 | 82.5 | 81.7 | 95.2 | 55.3 | 63.6 | 83.3 | 53.6 | 60 | 72.6 | 62.5 | 59.6 | 86.5 | 85.7 | 72.7 | 72.7 | 73 | 100 | 82.4 | 85.7 | 72.7 | 94.4 | 50 | 100 | 68.4 | 100 | 76.9 | 18.2 | 62.9 | 100 | 70.3 | 67.7 | 89.2 | 76 | 59.4 | 77.8 |
| Intervention, % |  |  |  |  |  |  |  |  |  |  |  |  |  |  |  |  |  |  |  |  |  |  |  |  |  |  |  |  |  |  |  |  |  |  |  |  |  |  |  |
| Medication | 49.7 | 50 | 6.2 | 100 | 63.7 | 45.4 | 33.3 | 78.9 | 18.2 | 100 | 32.8 | 100 | 44 | 56.2 | 65.8 | 78.8 | 74.6 | 72.7 | 54.5 | 69 | 50 | 47.1 | 35.7 | 45.5 | 83.3 | 0 | 100 | 57.9 | 50 | 66.7 | 33.3 | 22.9 | 100 | 25.3 | 30.6 | 81.9 | 72 | 68.8 | 81.5 |
| Biomedical | 26.7 | 33.3 | 12.5 | 0 | 32.5 | 20.6 | 4.8 | 26.3 | 0 | 83.3 | 27.2 | 0 | 13.1 | 31.2 | 31.6 | 11.5 | 27 | 36.4 | 9.1 | 23 | 35.7 | 47.1 | 42.9 | 18.2 | 30.6 | 0 | 0 | 5.3 | 25 | 15.4 | 8.3 | 20 | 60 | 11 | 19.4 | 30.1 | 12 | 40.6 | 18.5 |
| Intensive behavioral | 52.8 | 33.3 | 37.5 | 50 | 41.2 | 39.7 | 14.3 | 44.7 | 27.3 | 50 | 51.3 | 0 | 34.5 | 21.9 | 52.8 | 30.8 | 25.4 | 81.8 | 54.5 | 31 | 42.9 | 64.7 | 57.1 | 54.5 | 50 | 50 | 0 | 47.4 | 50 | 23.1 | 58.3 | 55.7 | 100 | 37.4 | 30.6 | 36.1 | 48 | 40.6 | 33.3 |

DCDQ: the developmental coordination disorder questionnaire, RBS-R: repetitive behavior scale-revised, SCQ: social communication questionnaire, SD: standerd deviation

### Gene interpretation

We observed 90 chromosomal loci that satisfied the threshold of *P* < 5.0 × 10^−8^. The results of each GWAS analysis with more than the sample size of 9 cases after clustering are shown in Figure 2; 17 out of 38 loci were located within 16 genes, while the remaining 52 were intergenic (Table 2). Twenty-nine loci were located within or near the genes associated with the Human Gene module of the SFARI Gene scoring system (*6*), such as *RFX3* (score 1, Rare Single Gene Mutation, Syndromic) in Cluster 15; *HCN1* (score S, Rare Single Gene Mutation, Genetic Association) in Cluster 18; *CSMD1* (score 3, Rare Single Gene Mutation, Genetic Association) in Cluster 23; *HIVEP3* (score 2, Rare Single Gene Mutation, Genetic Association) in Cluster 24; and *CNTNAP2* (score 2S, Rare Single Gene Mutation, Syndromic, Genetic Association) in Cluster 30.

**Figure 2.**
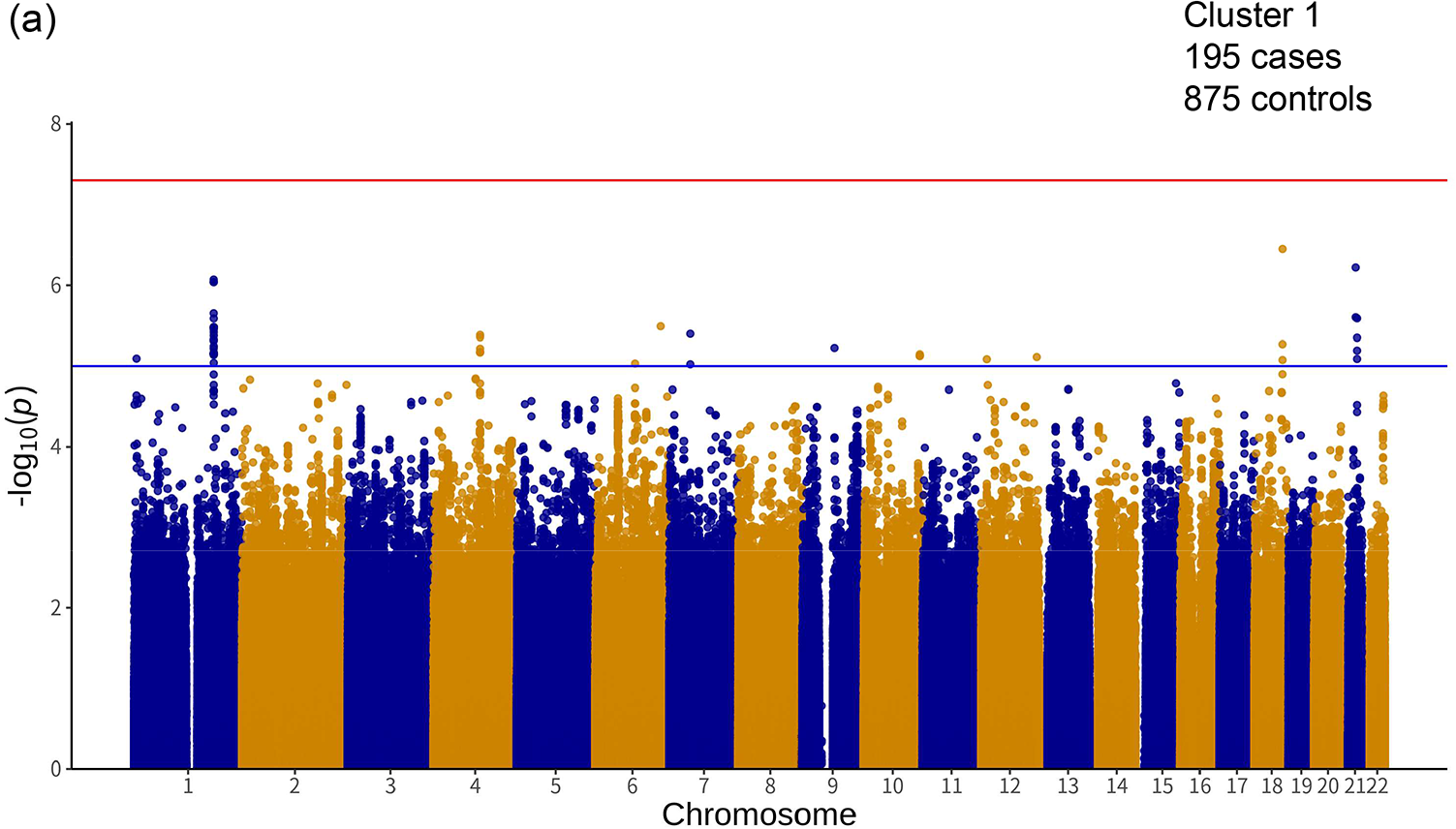
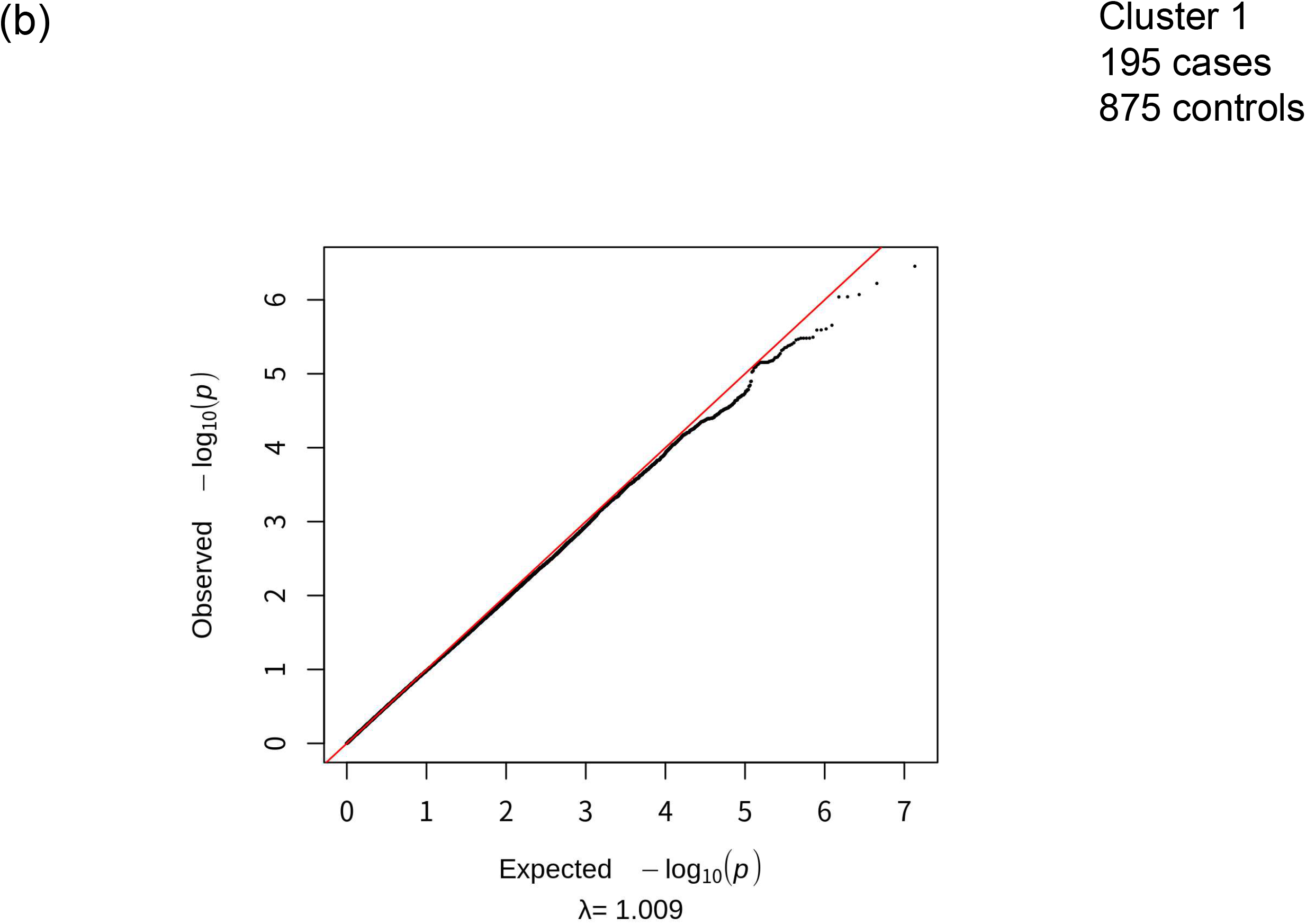
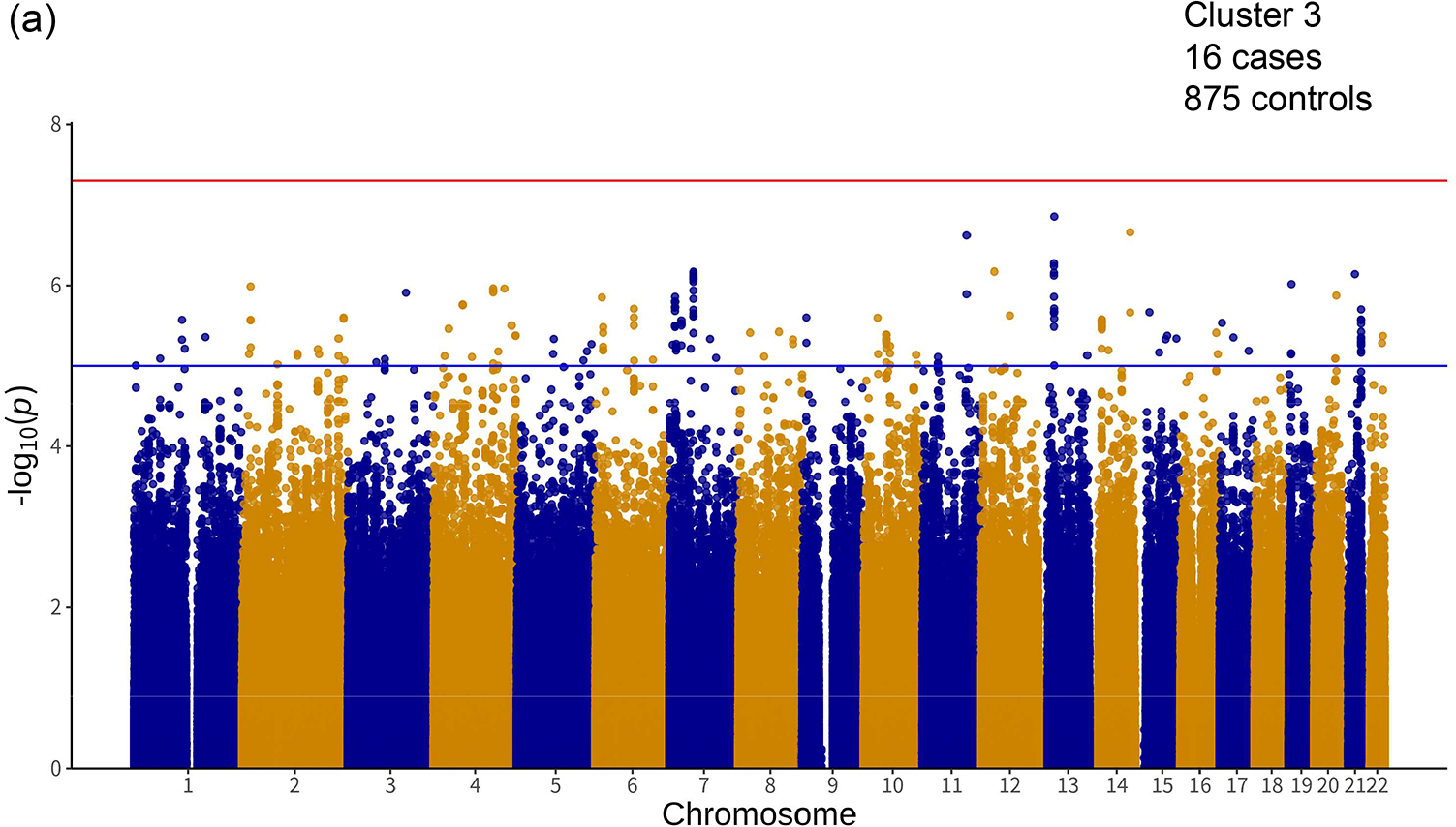
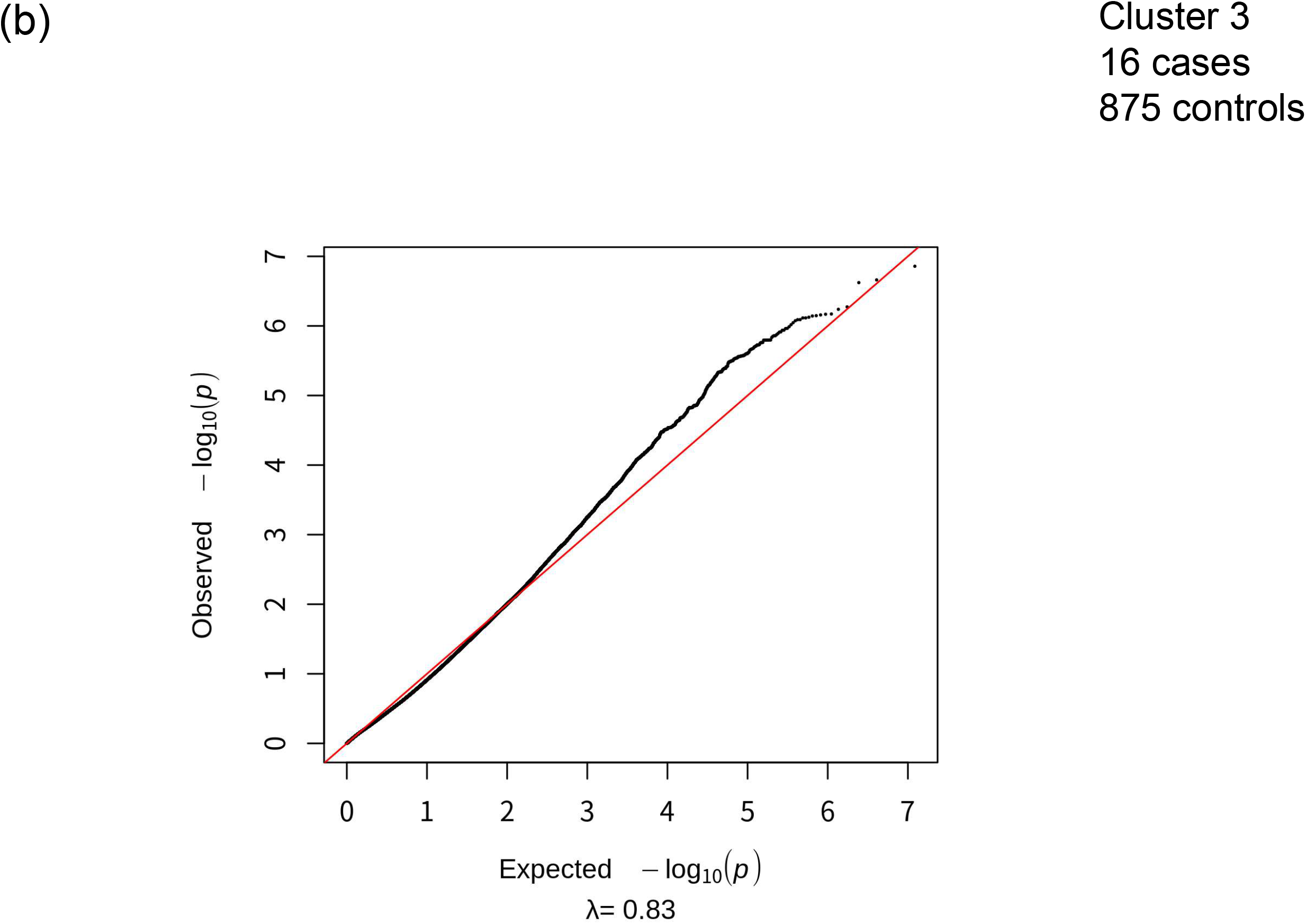
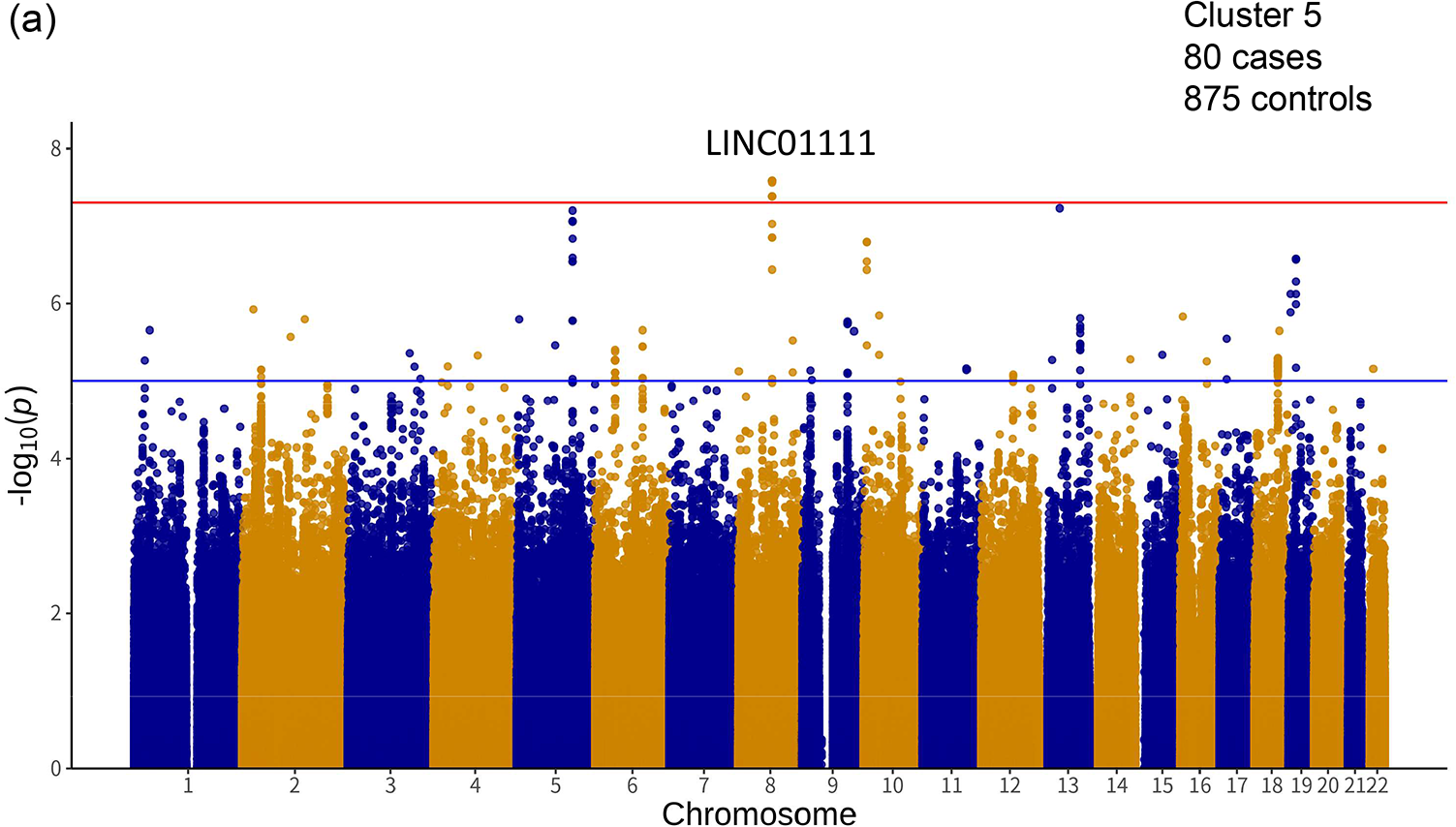
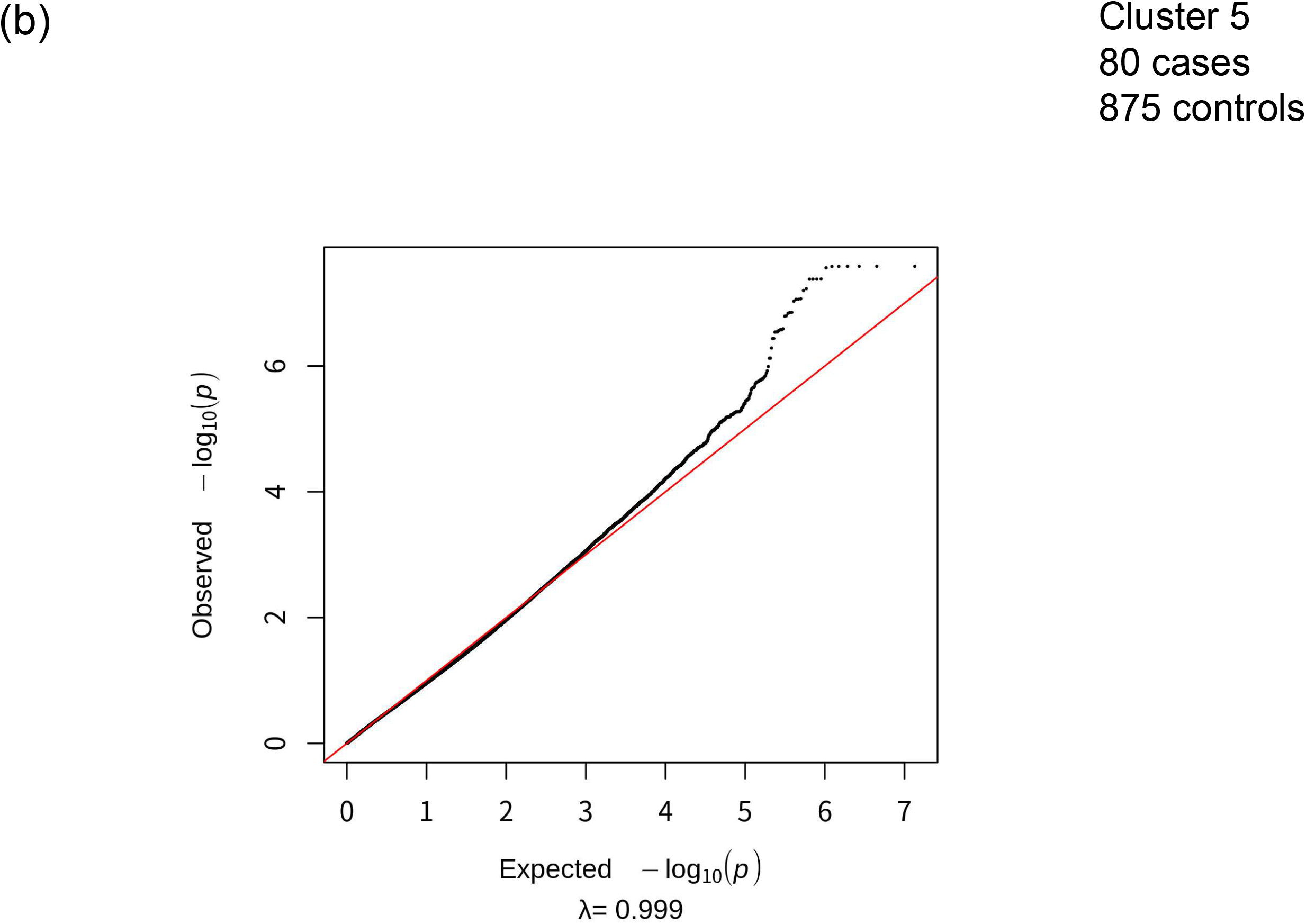
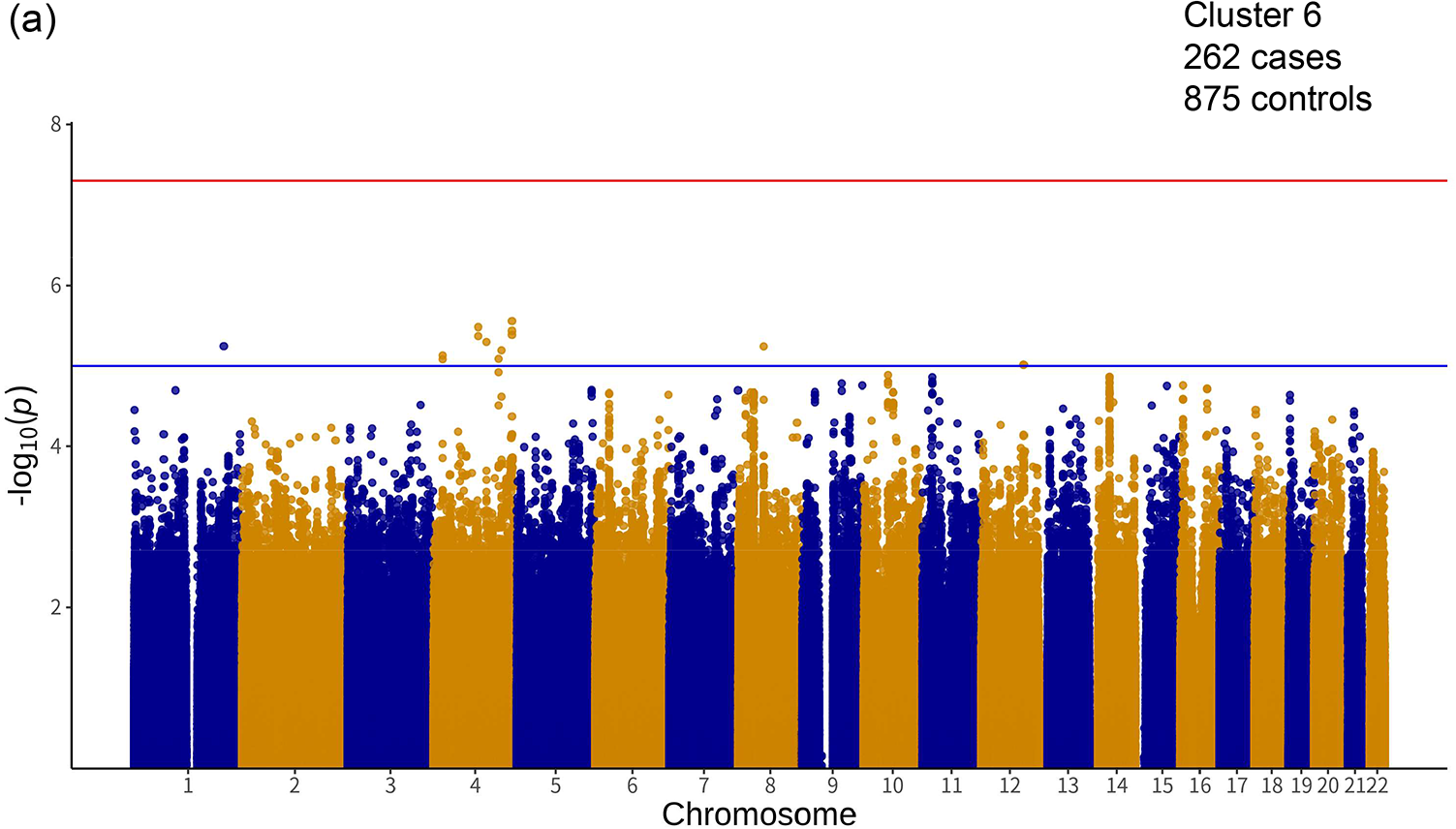
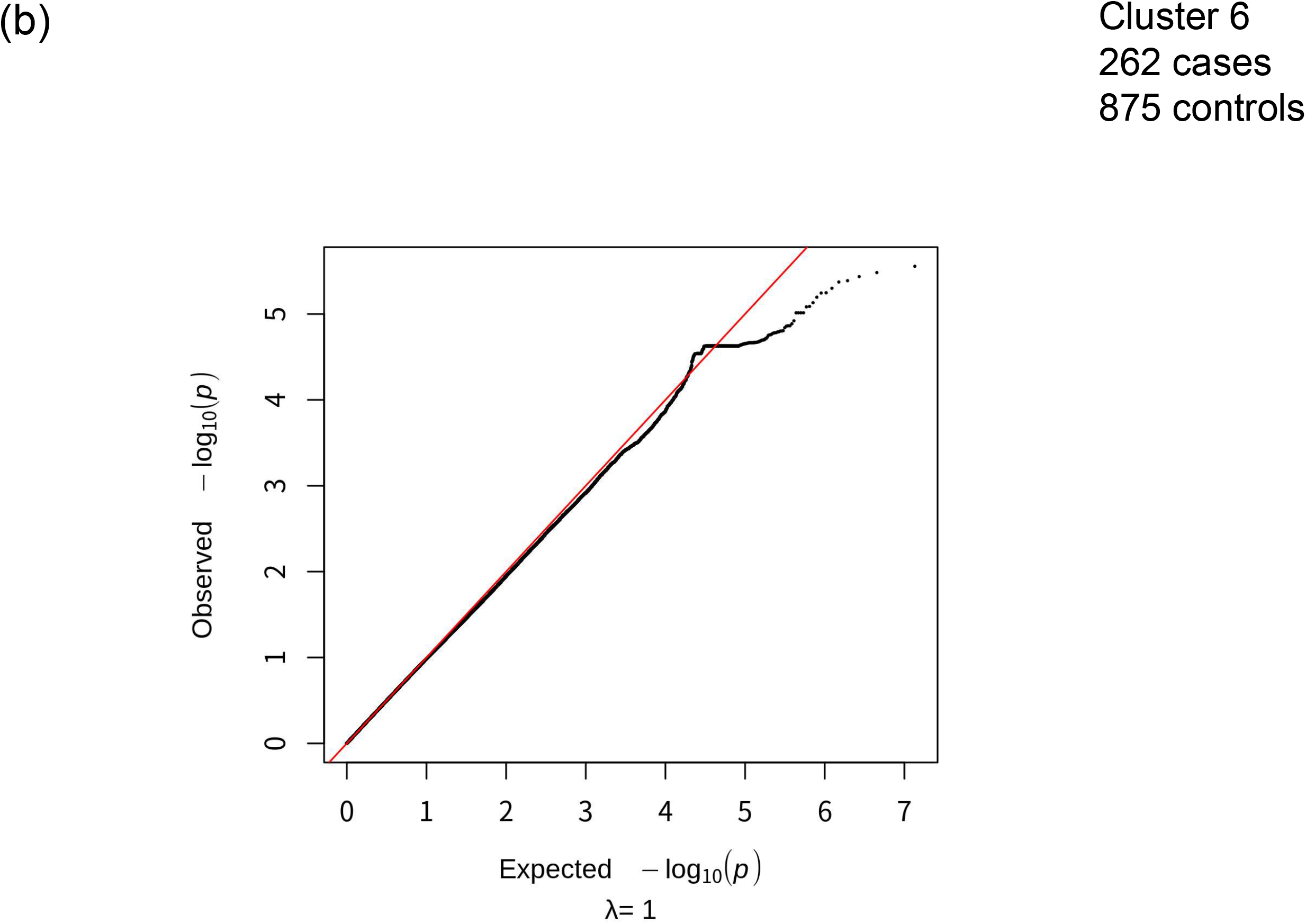
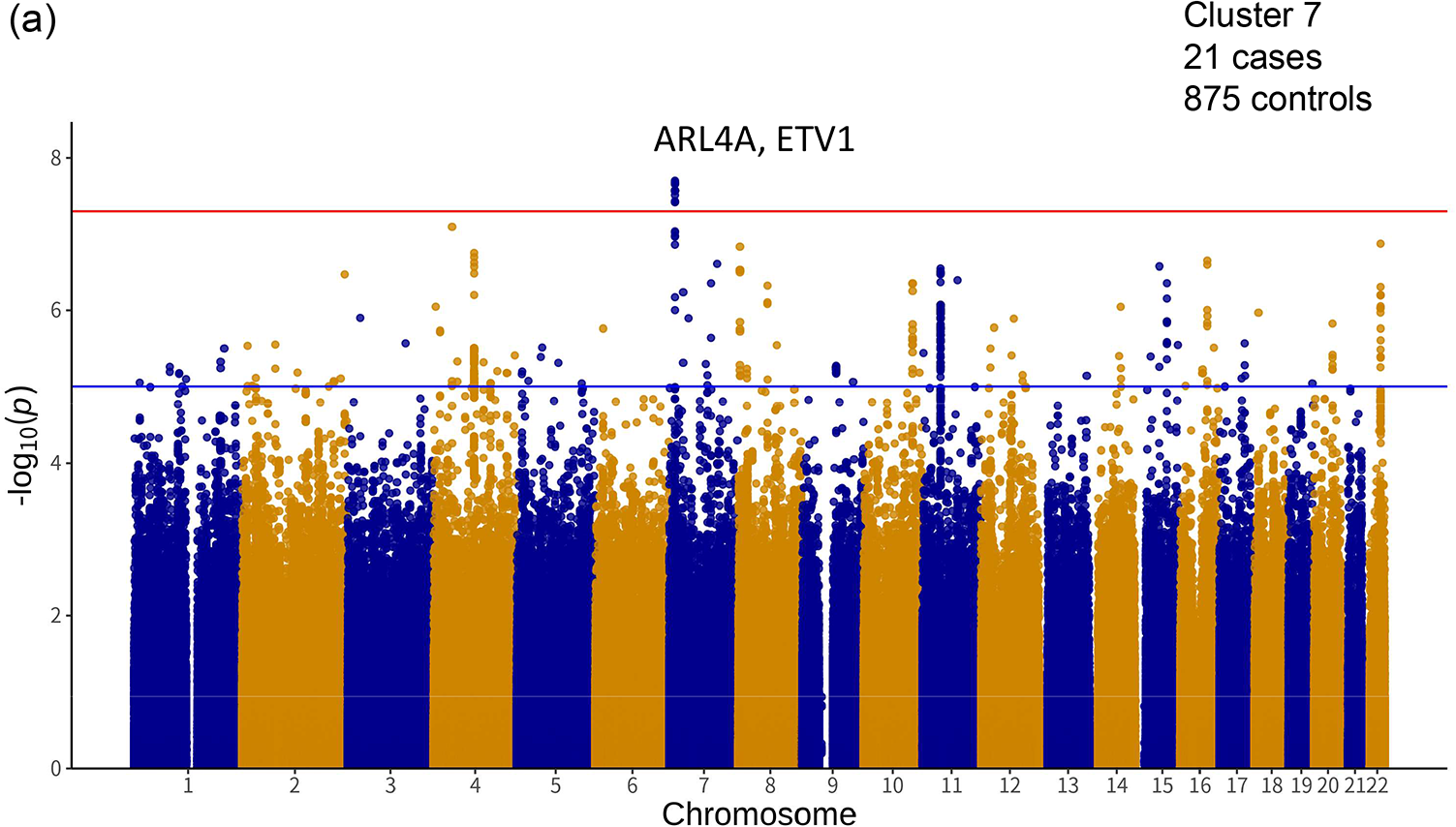
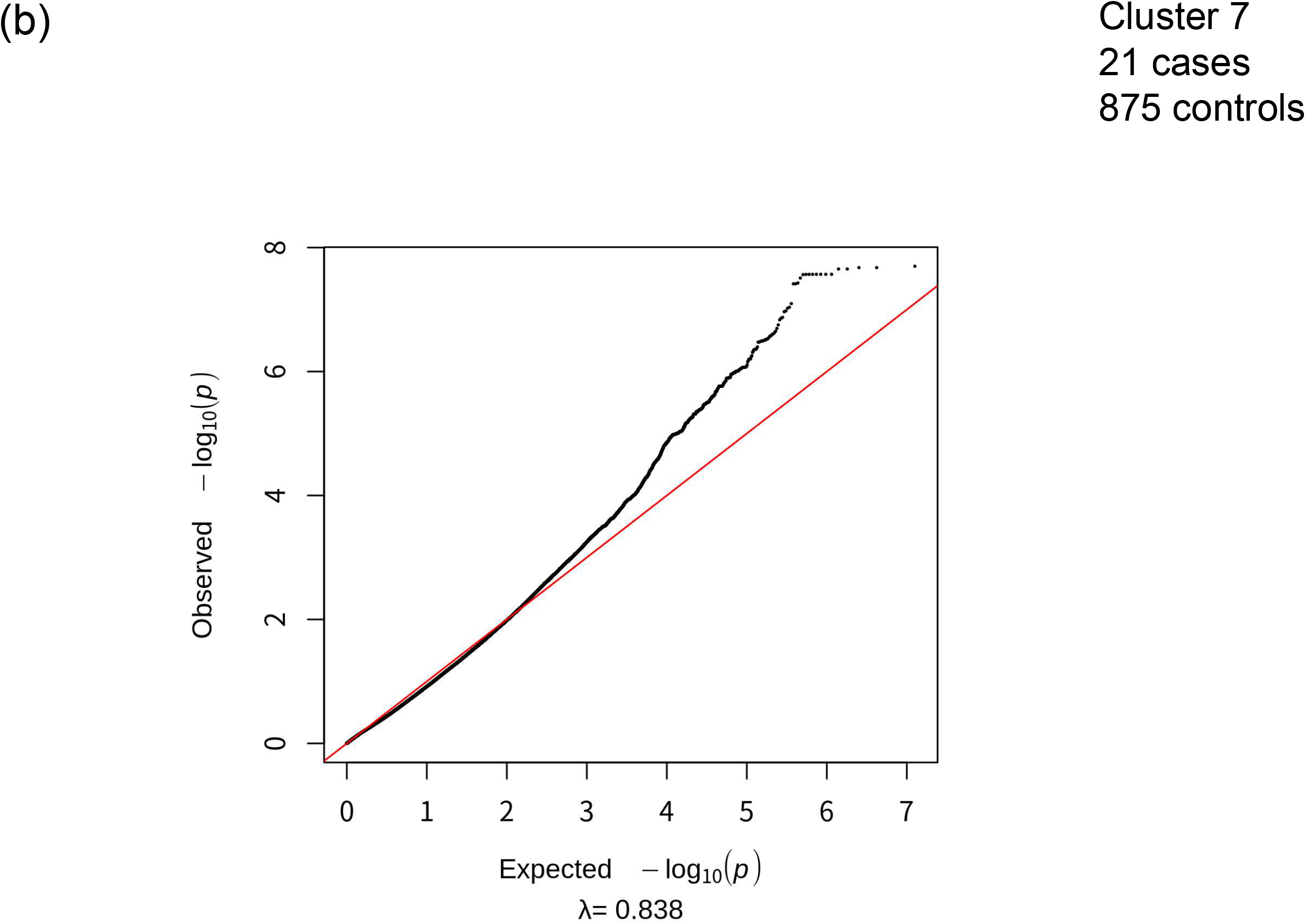
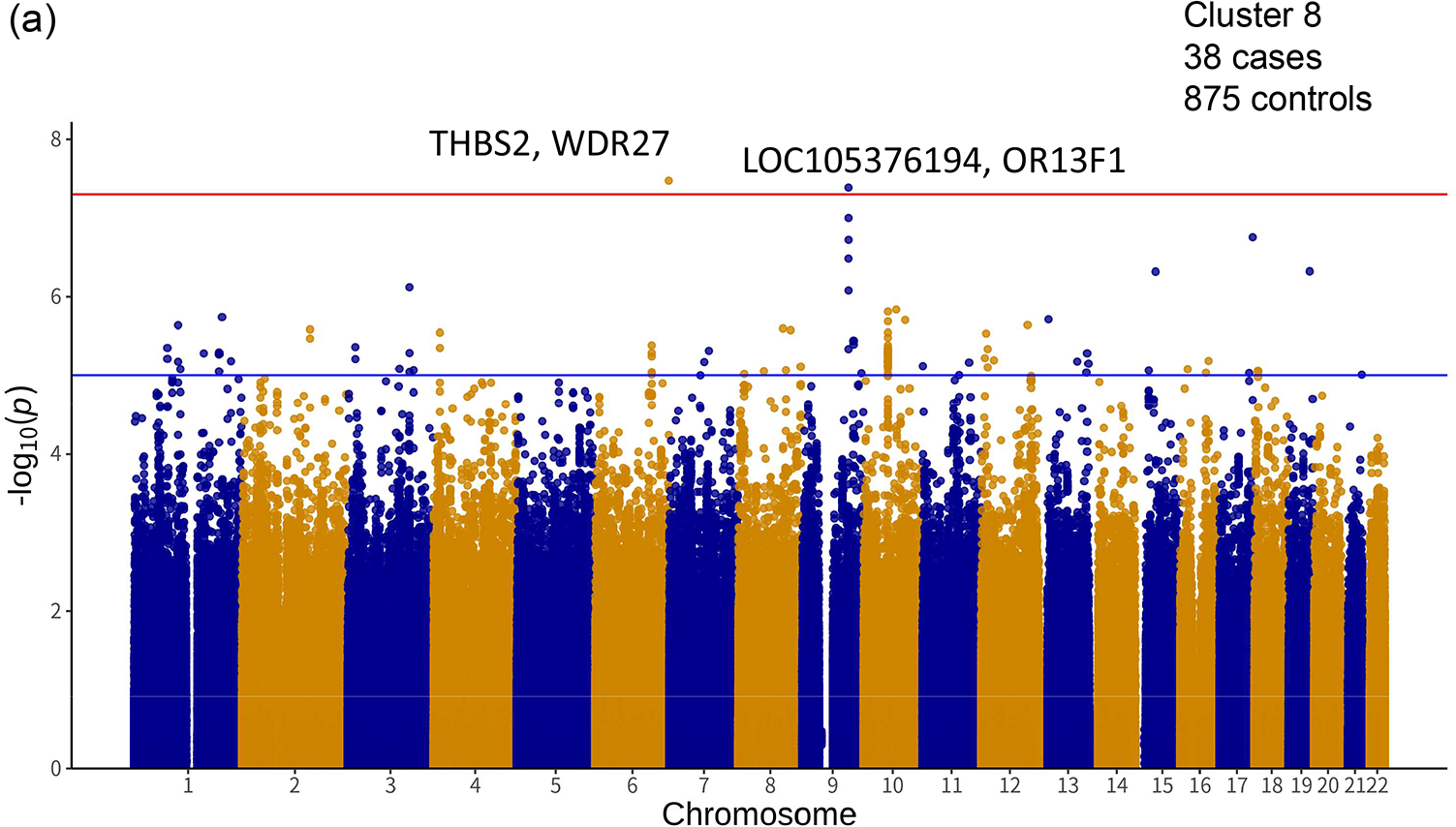
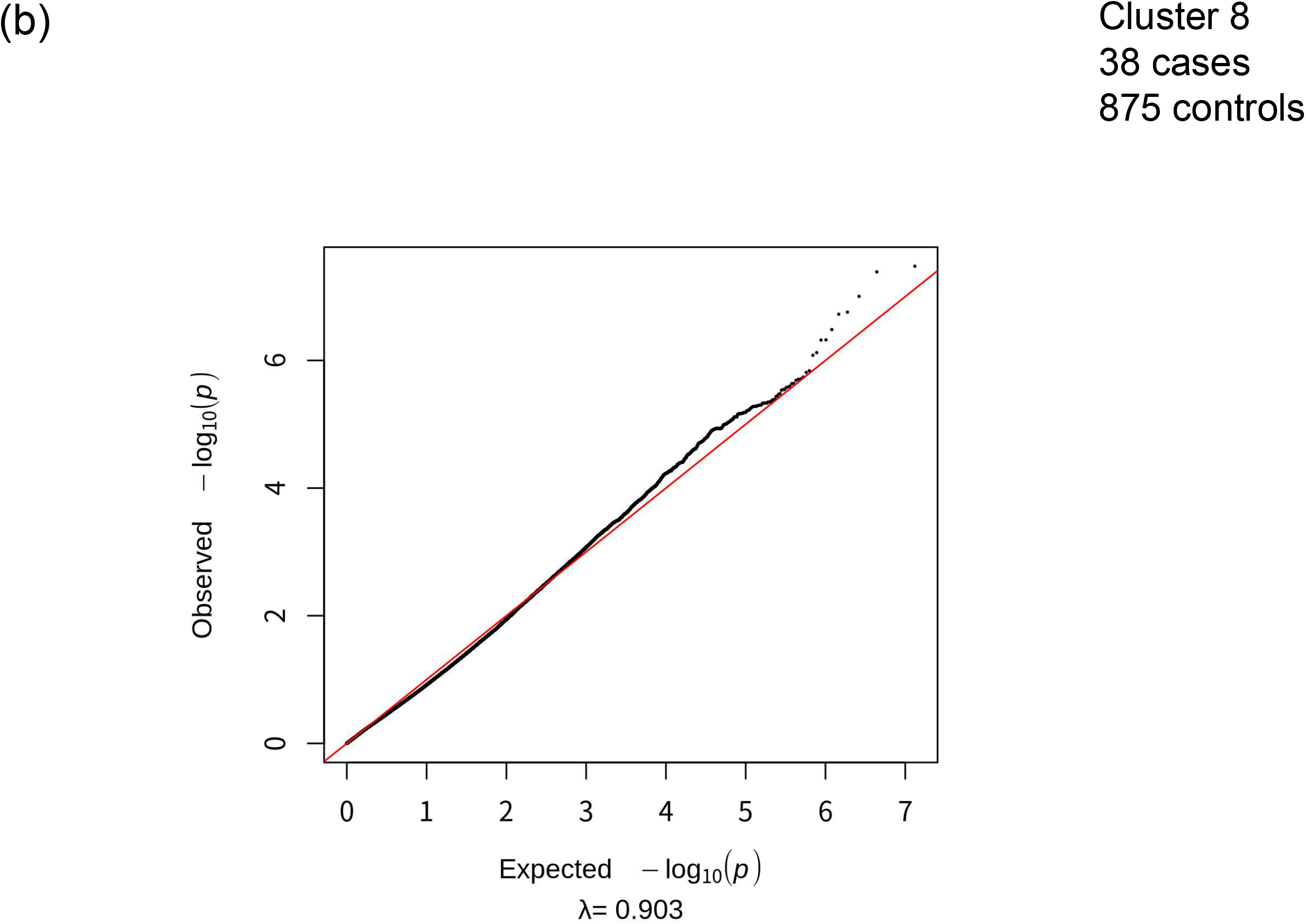
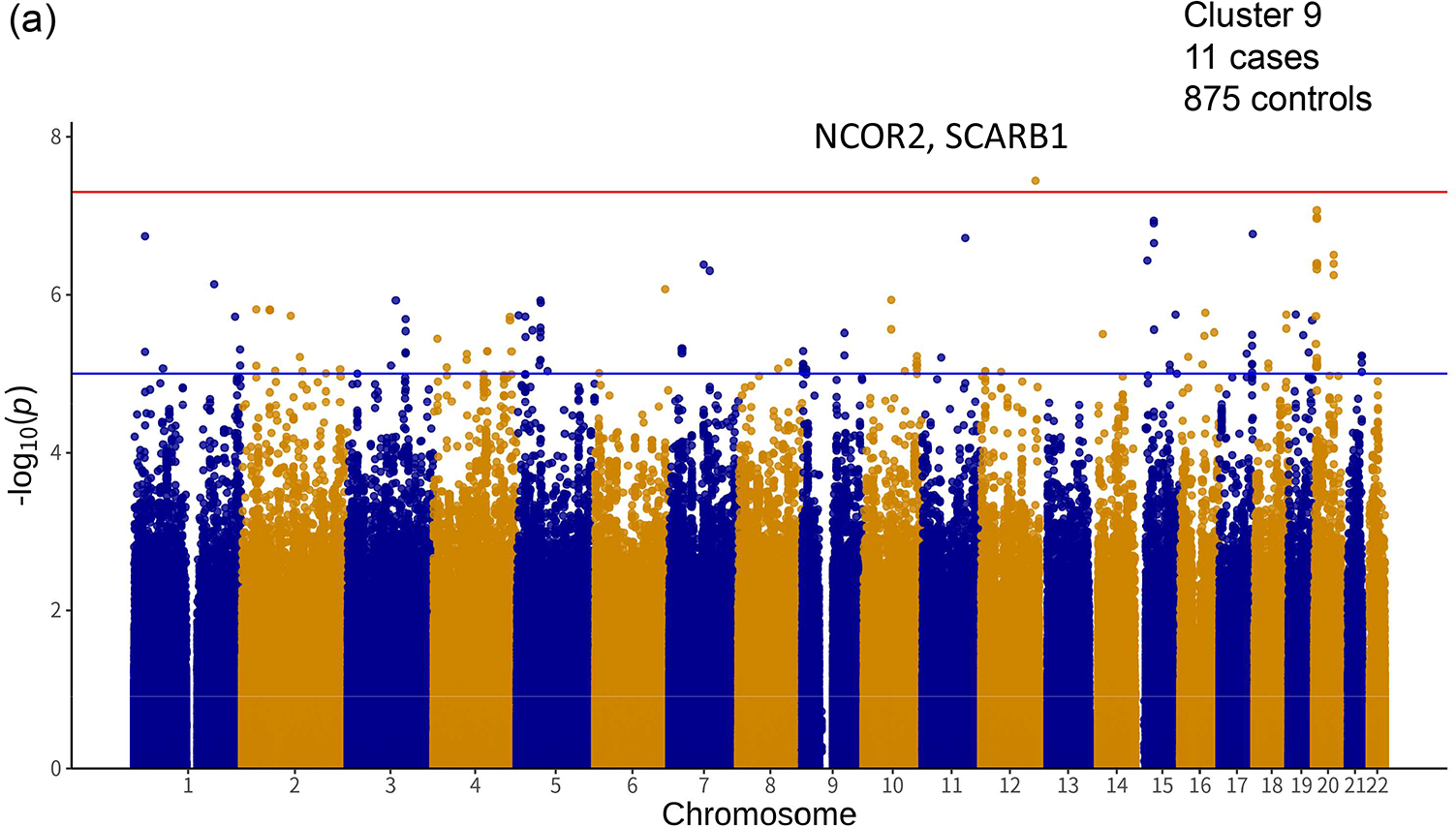
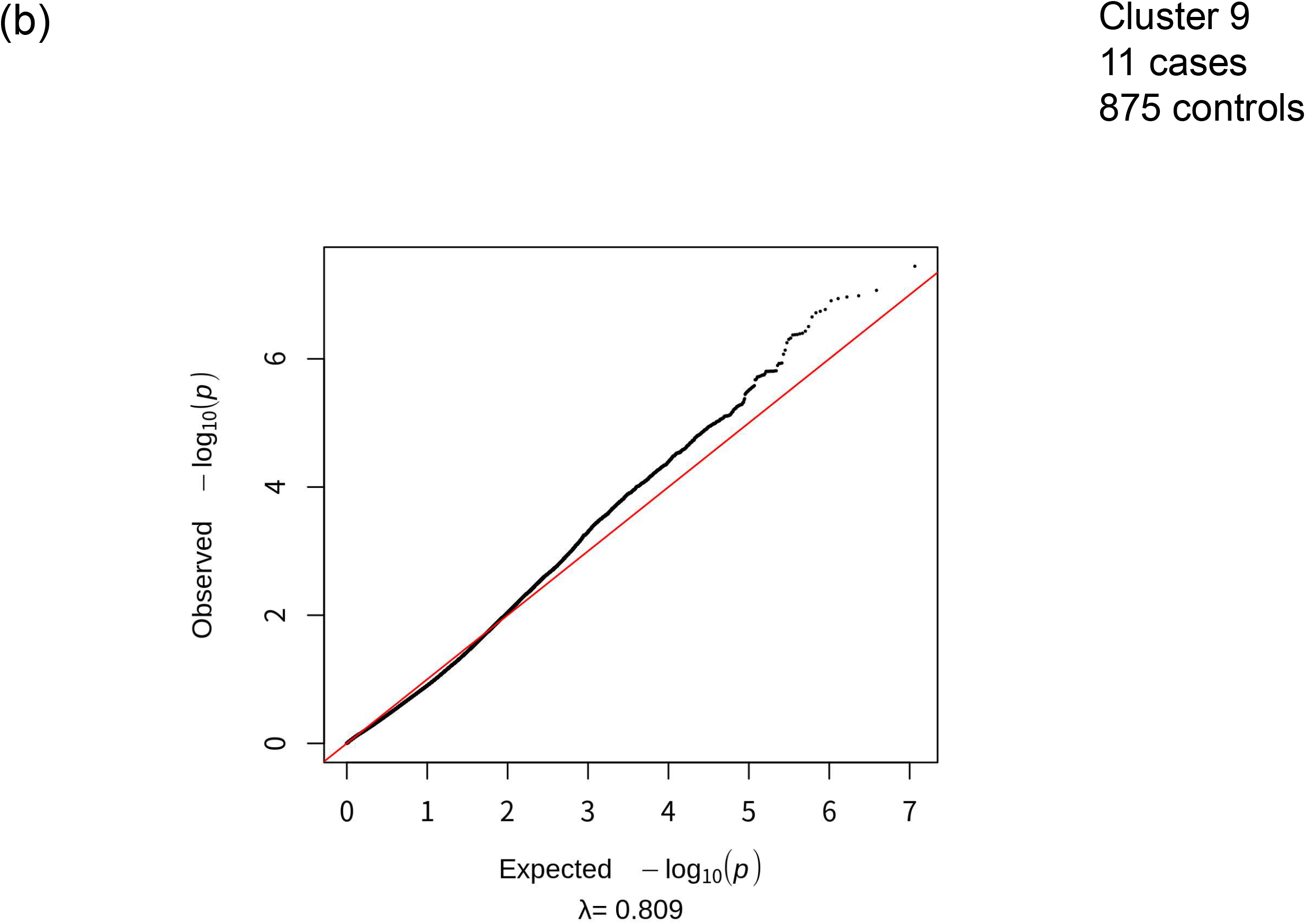
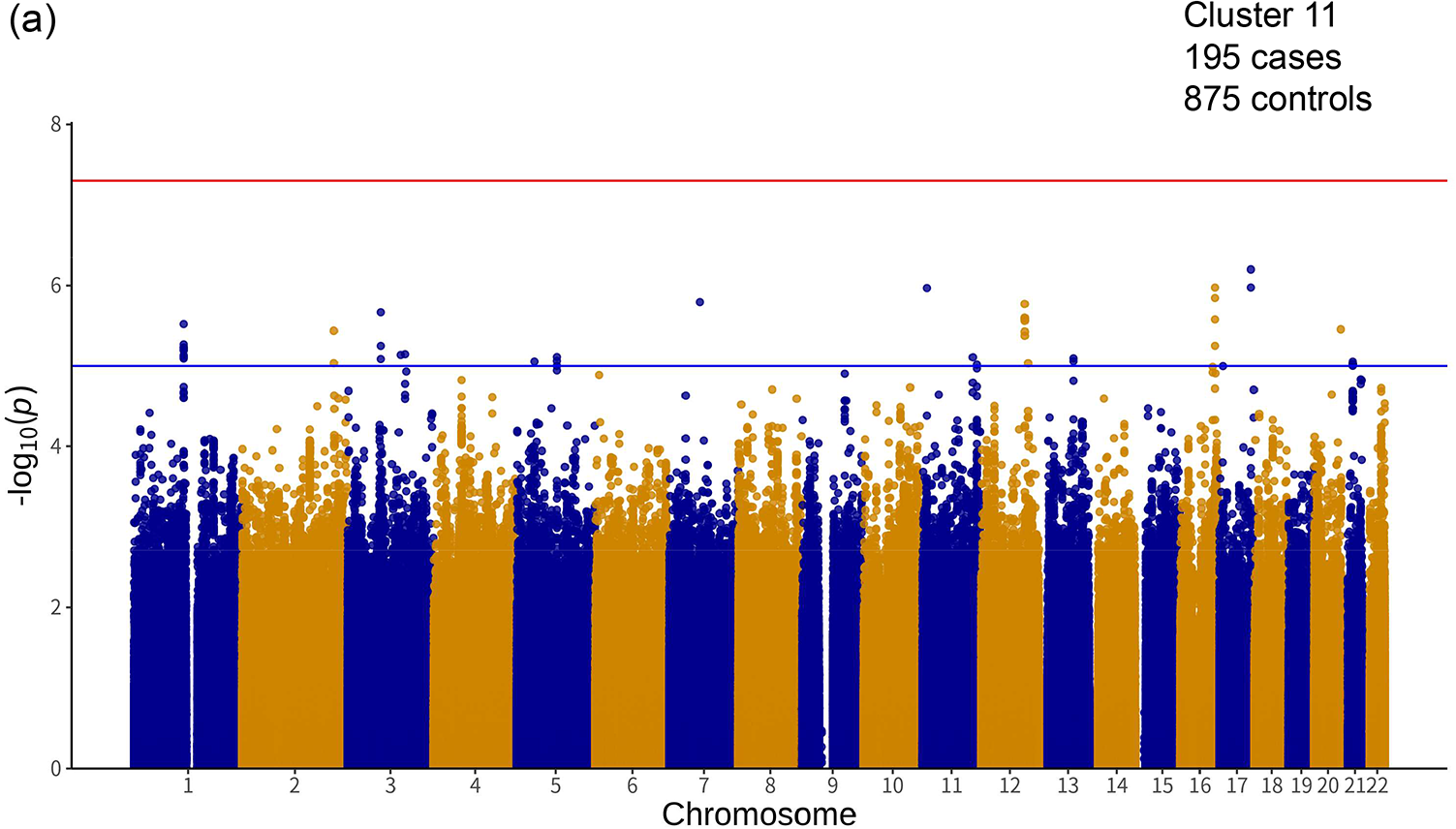
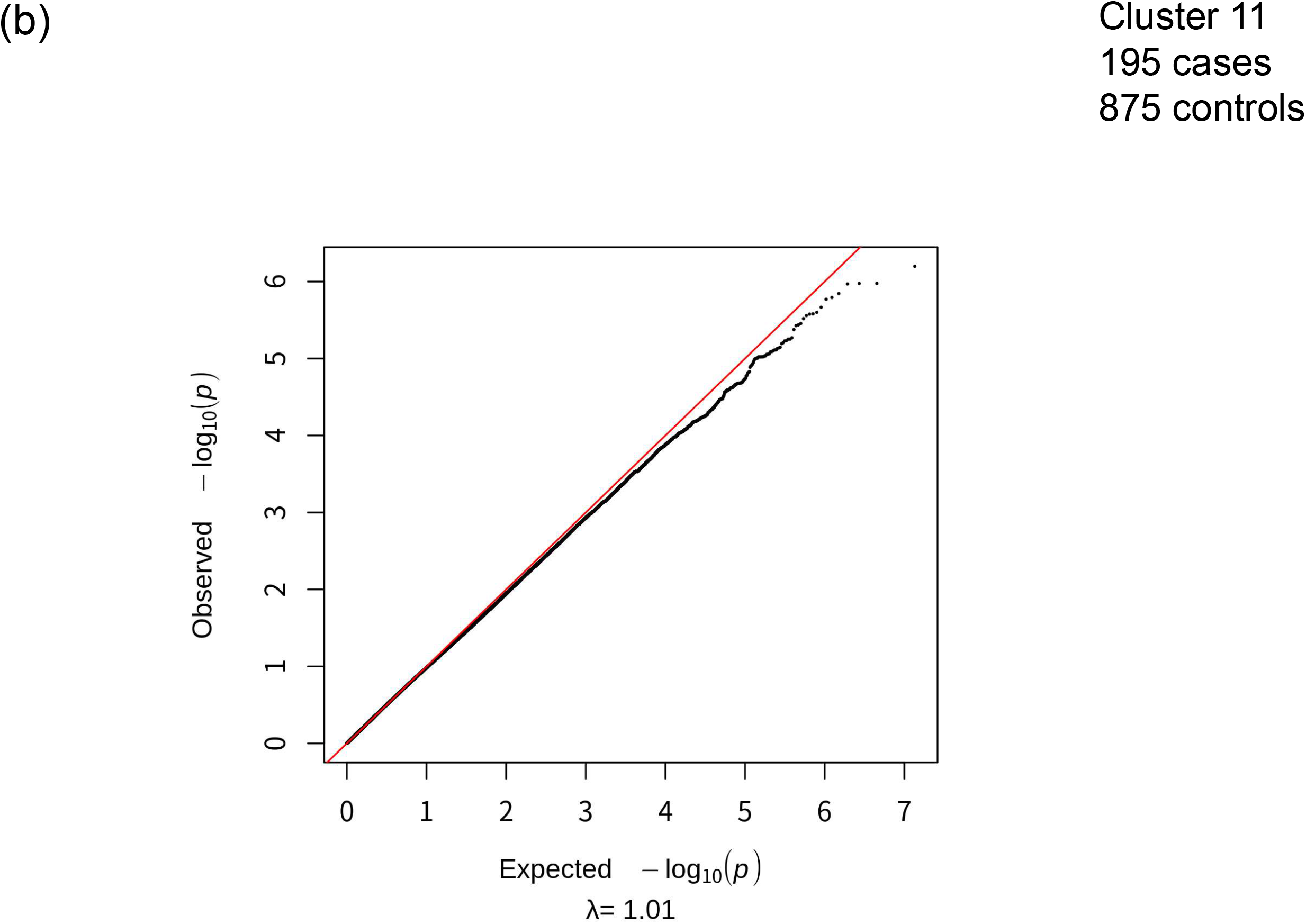
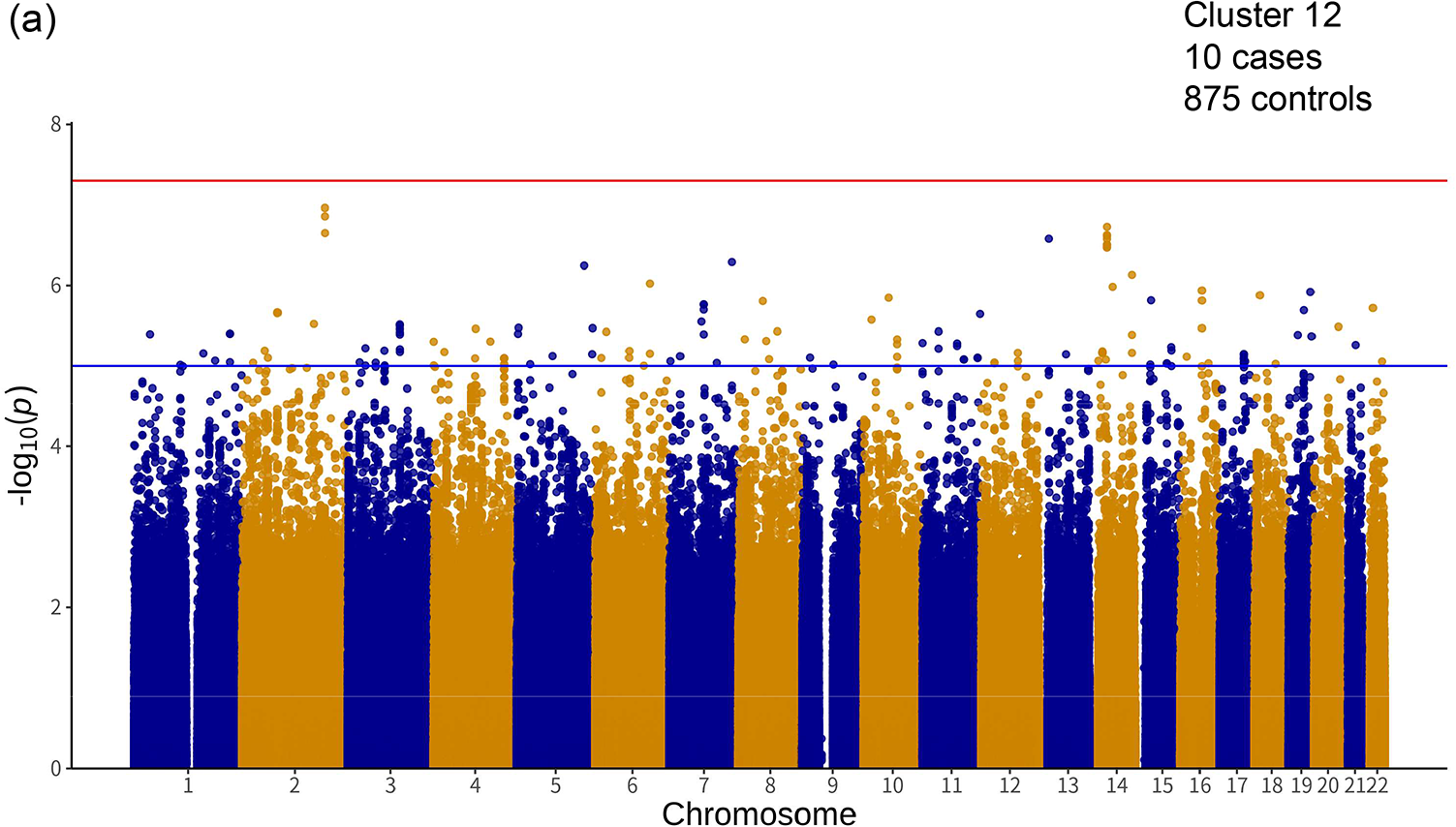
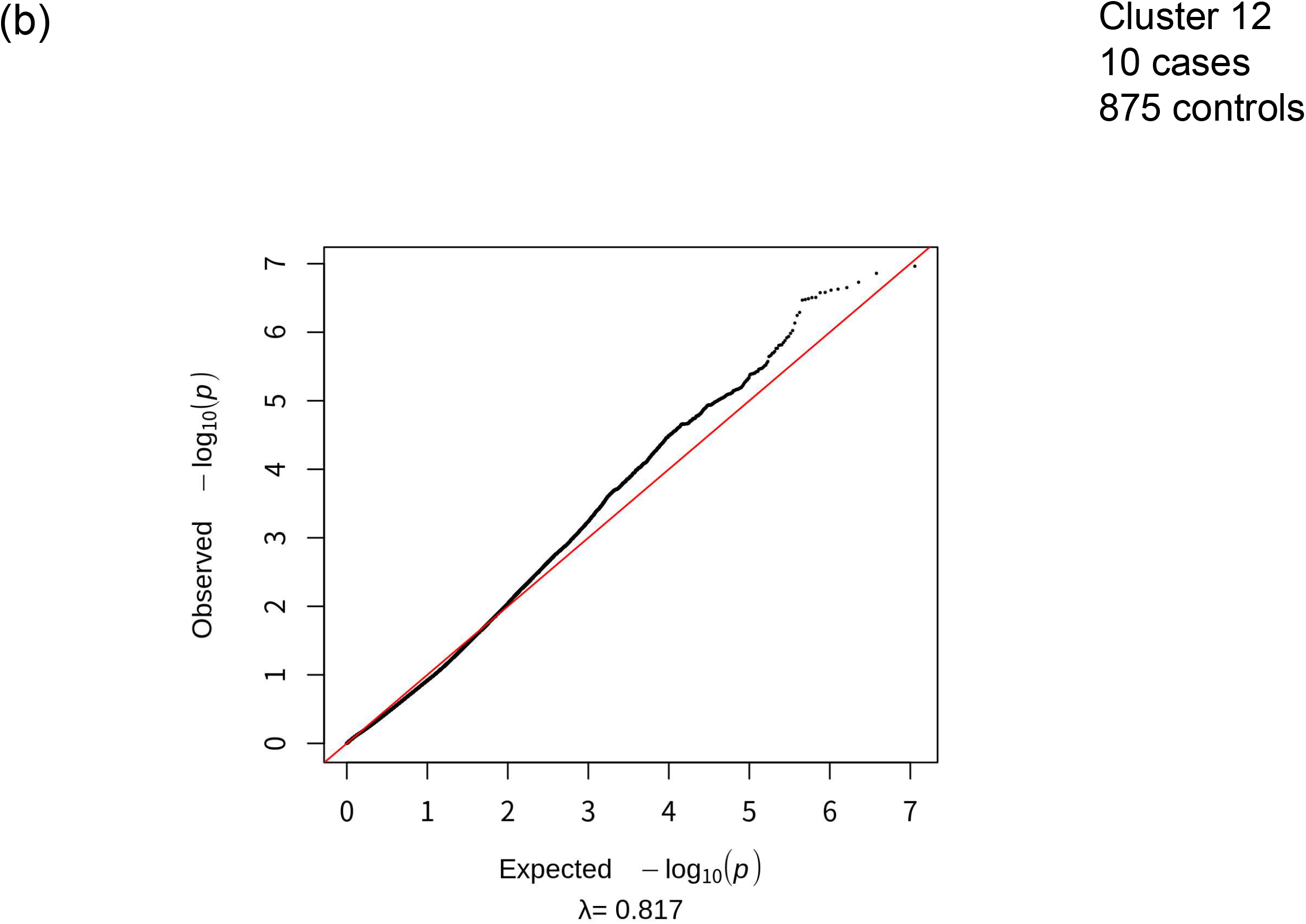
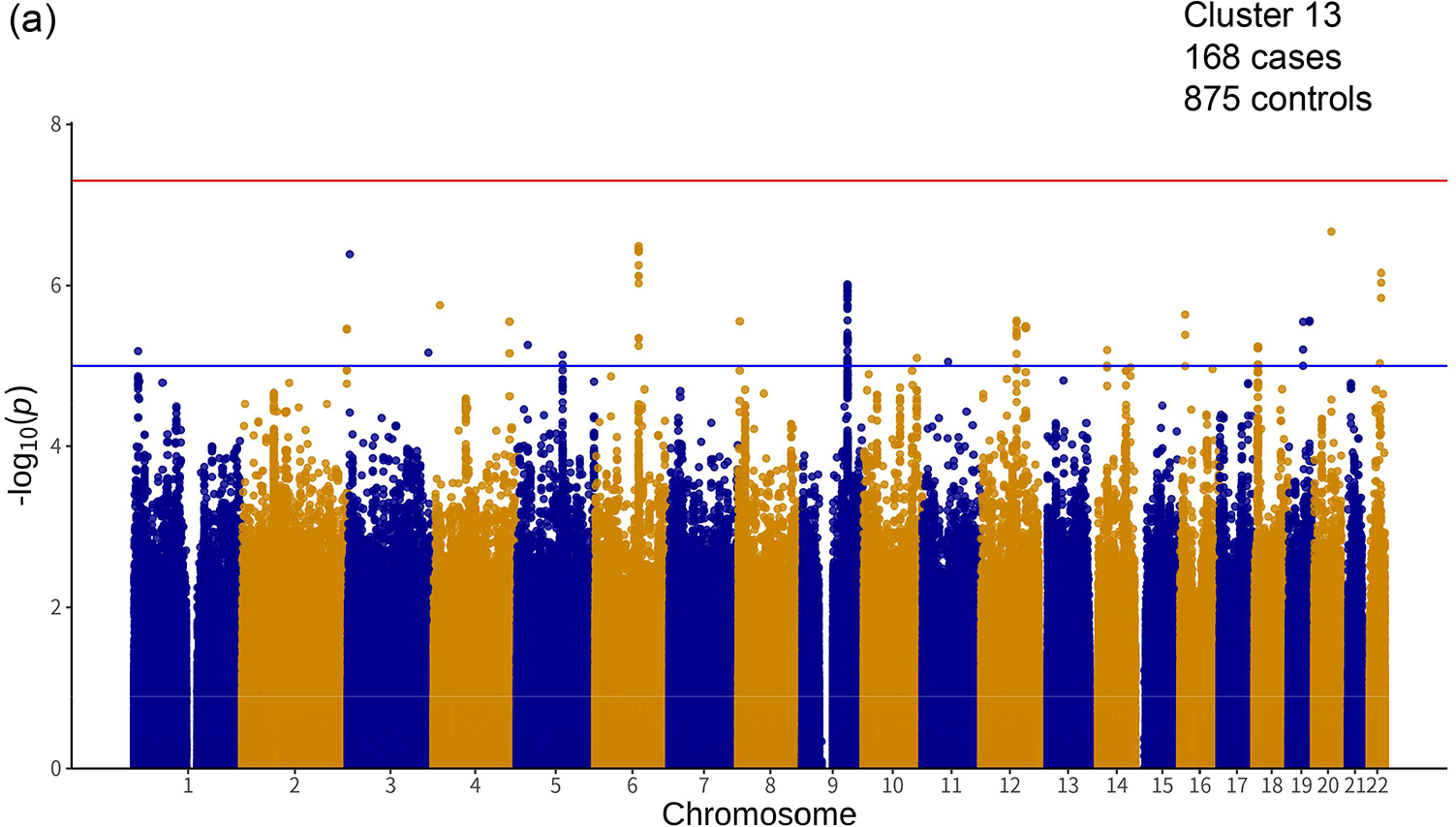
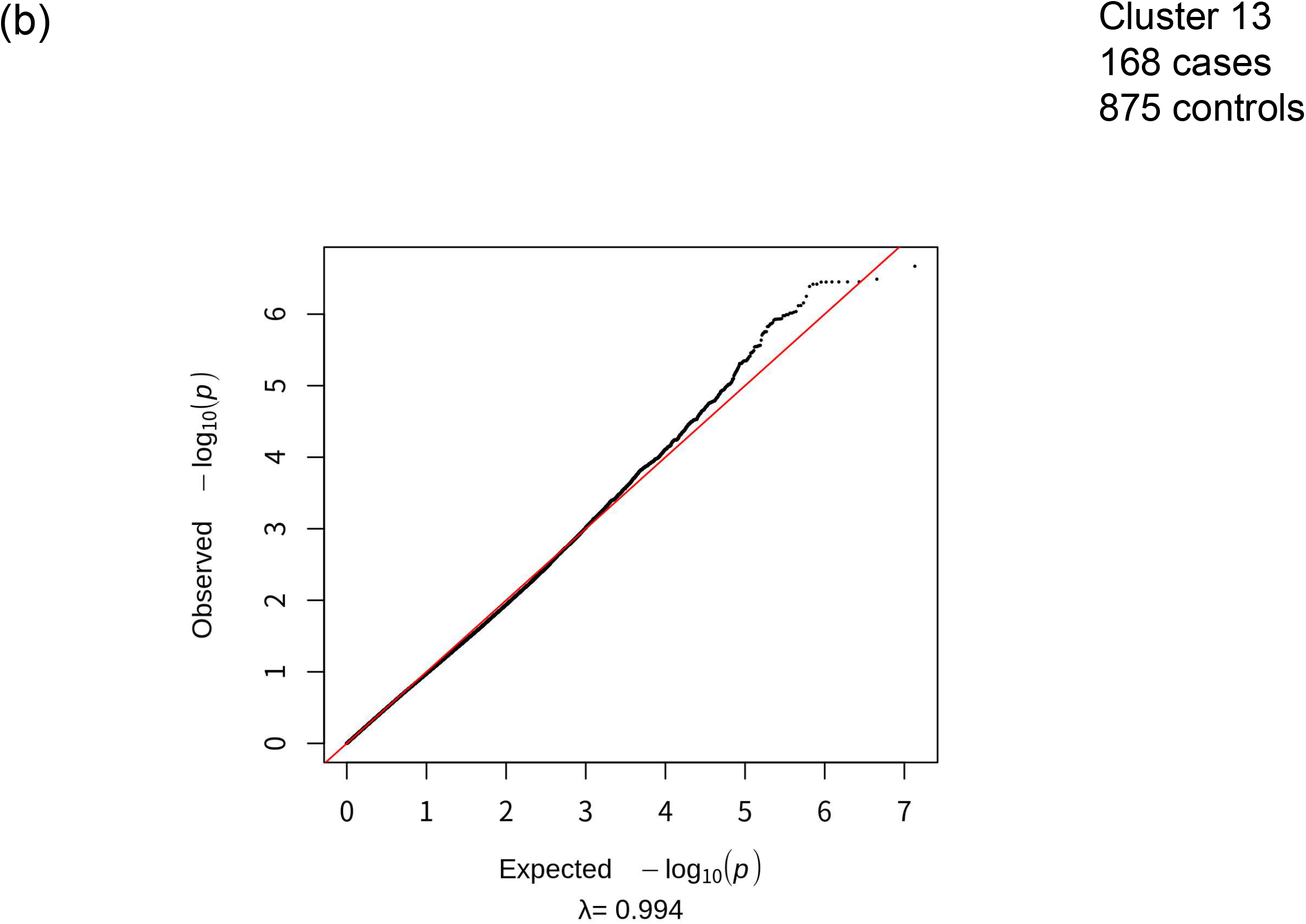
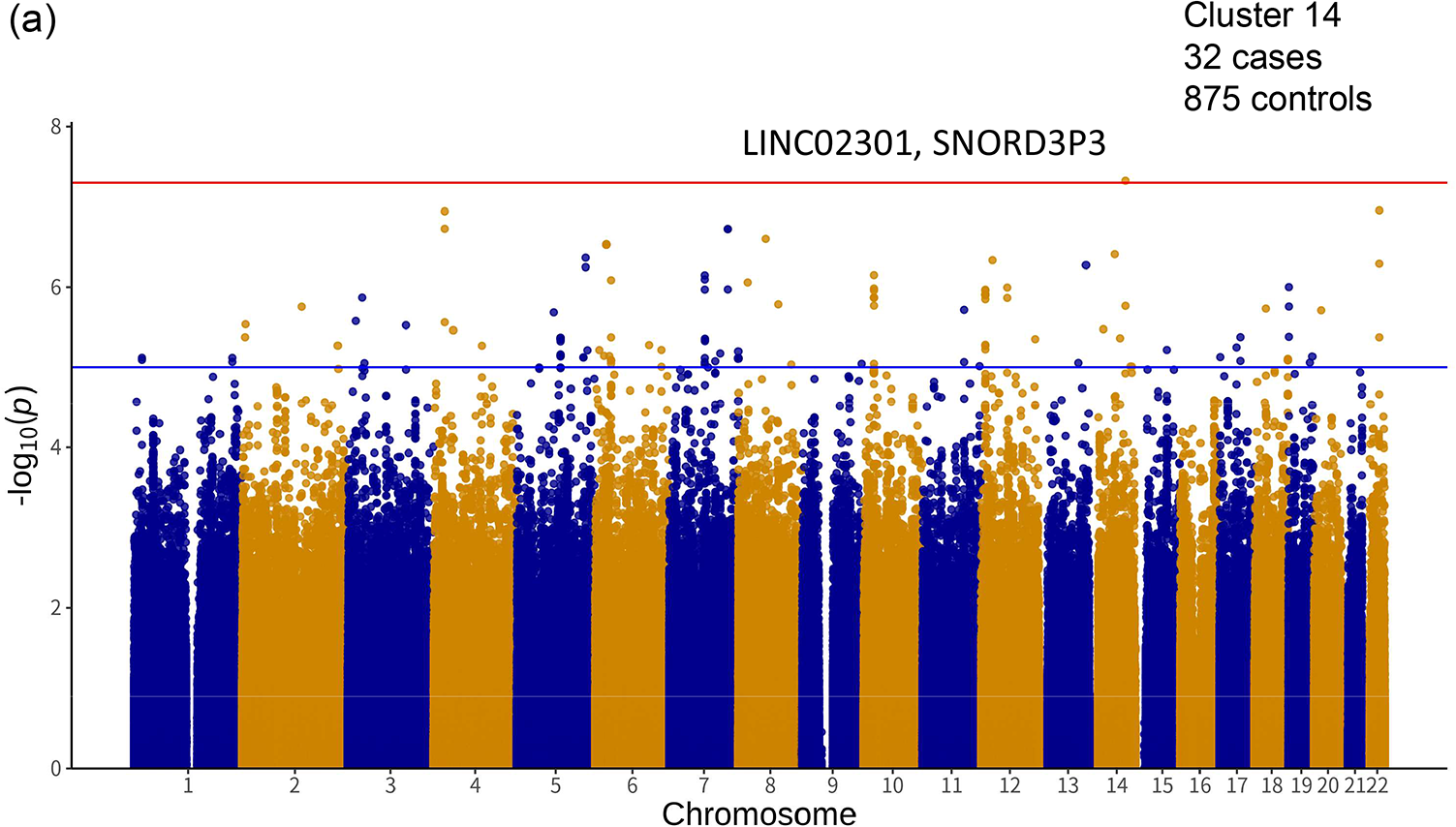
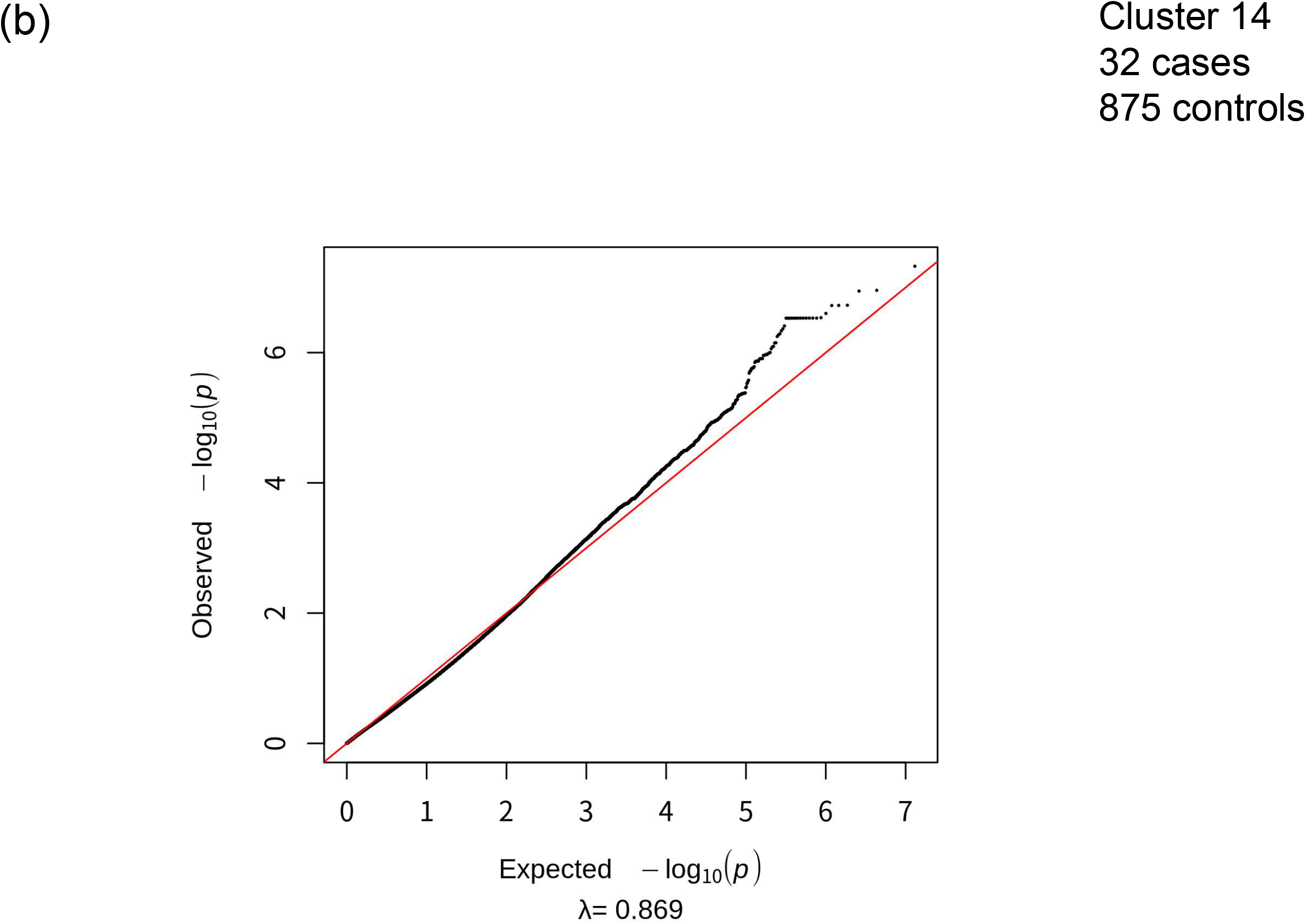
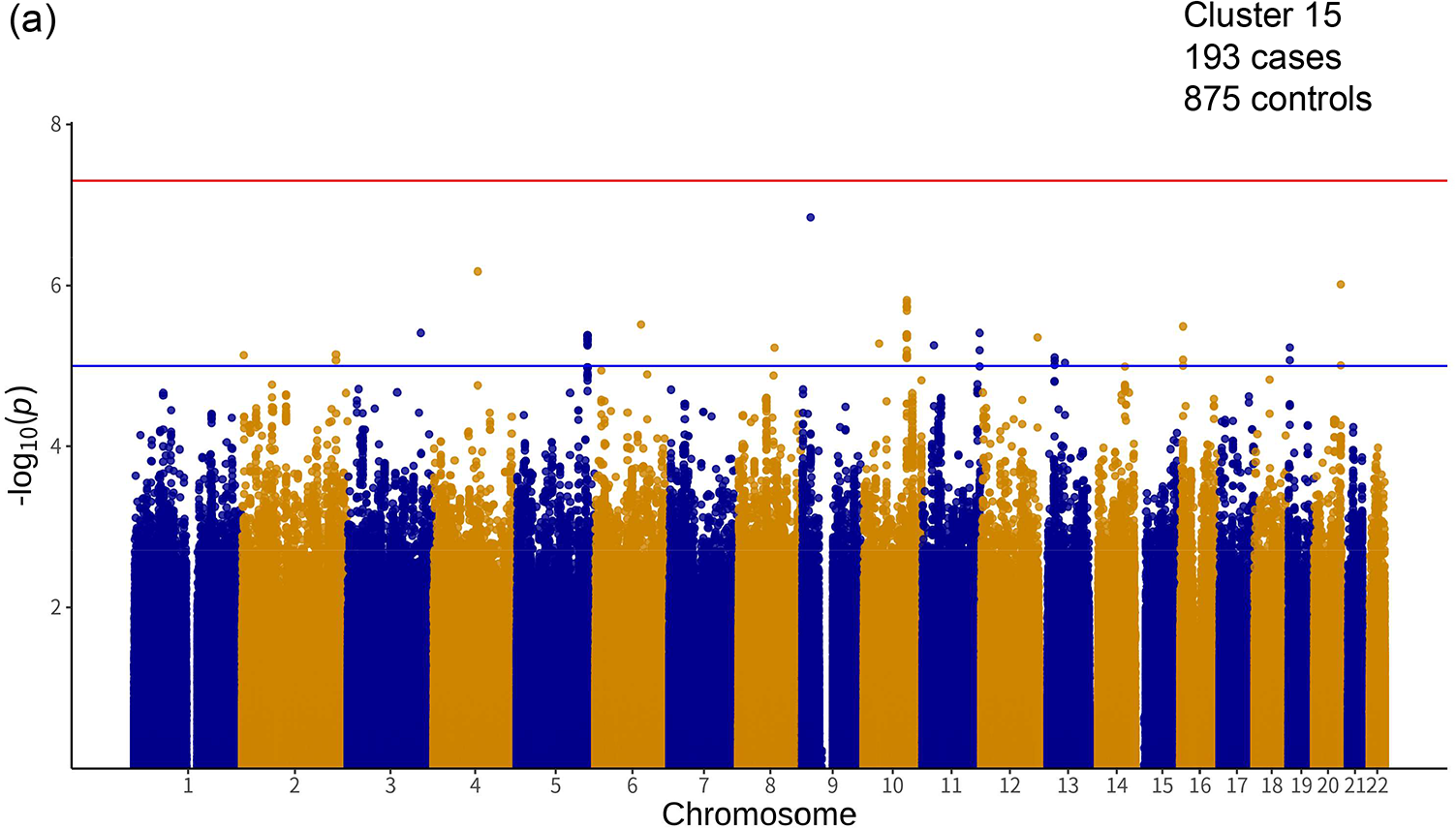
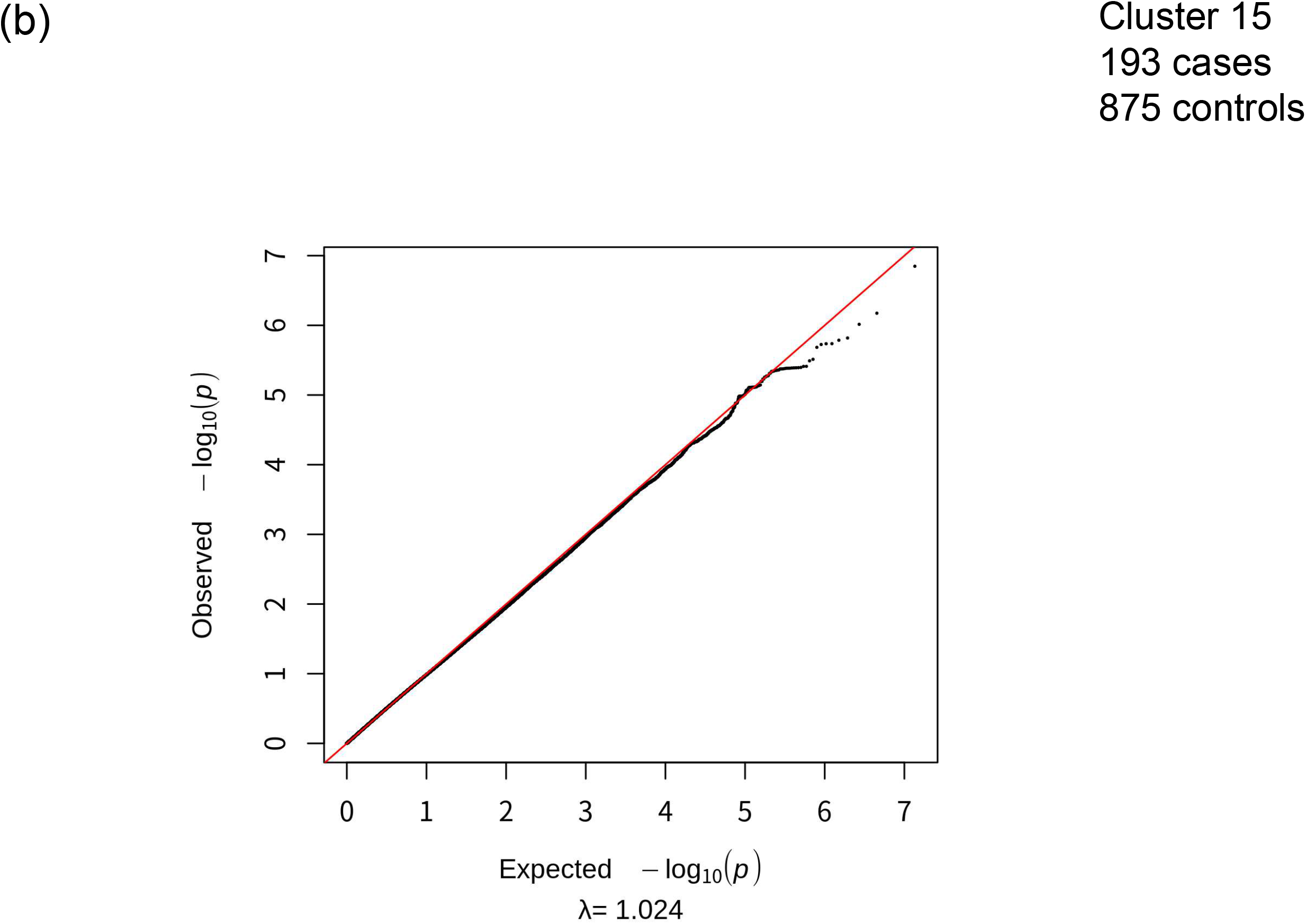
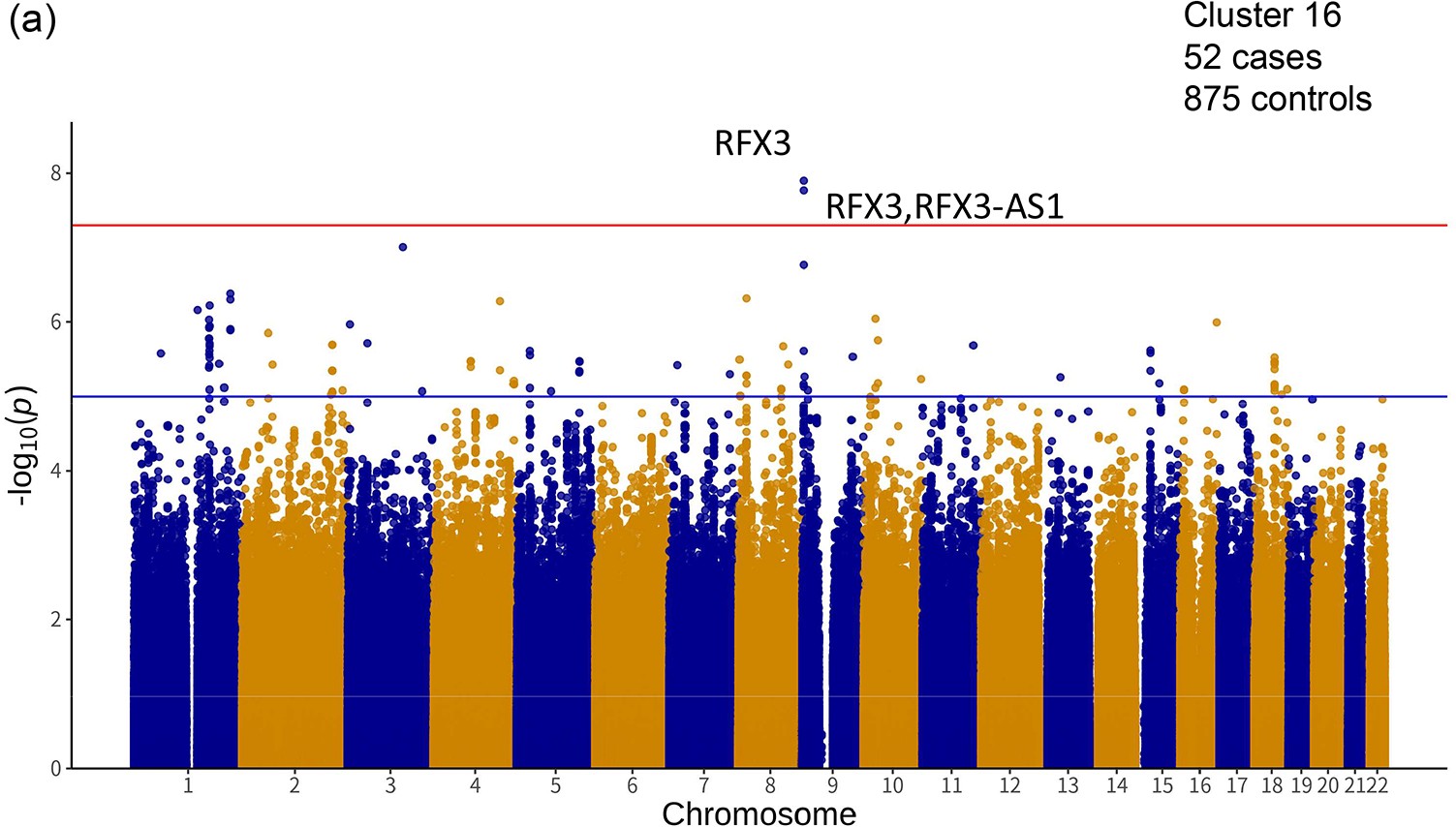
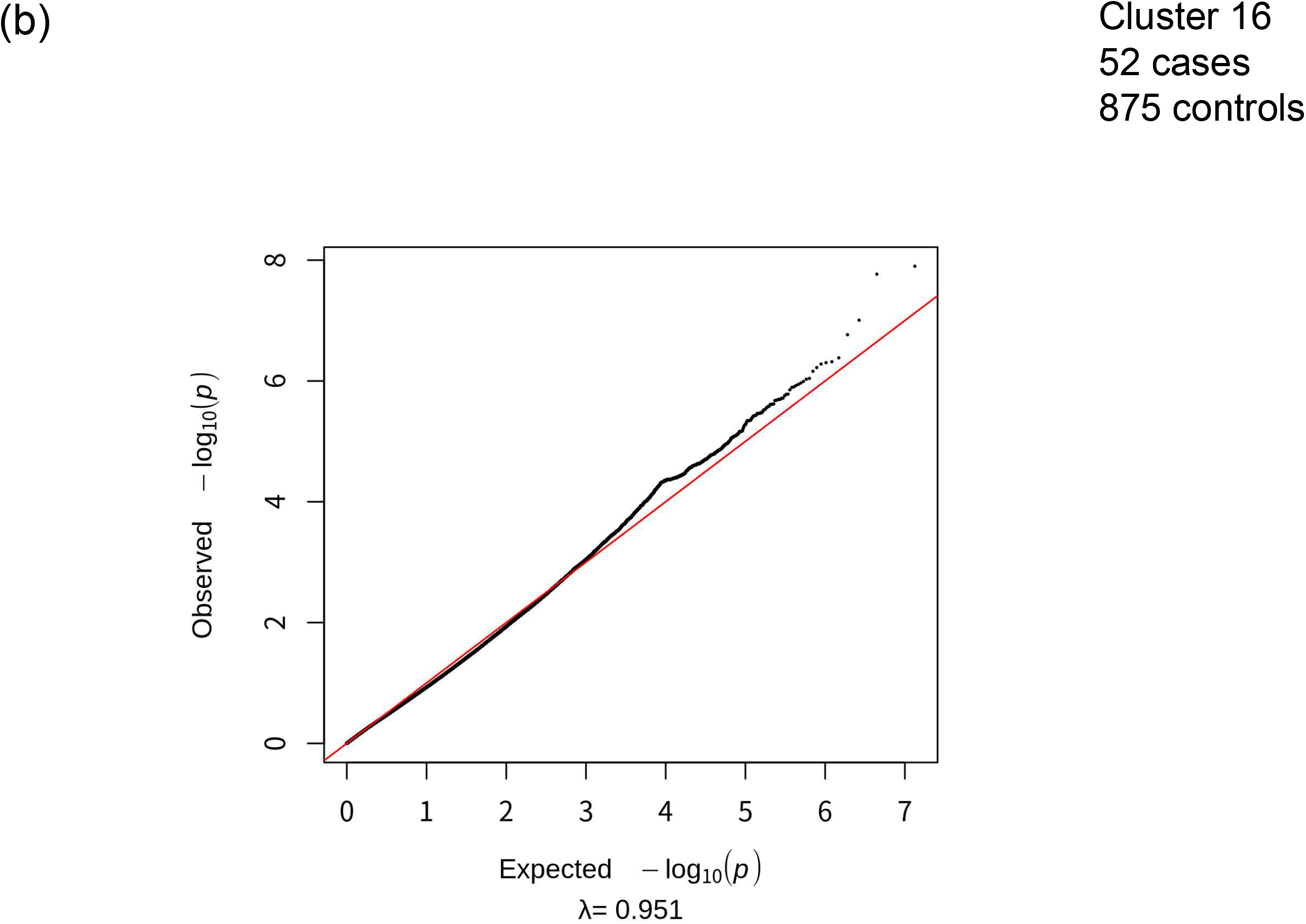
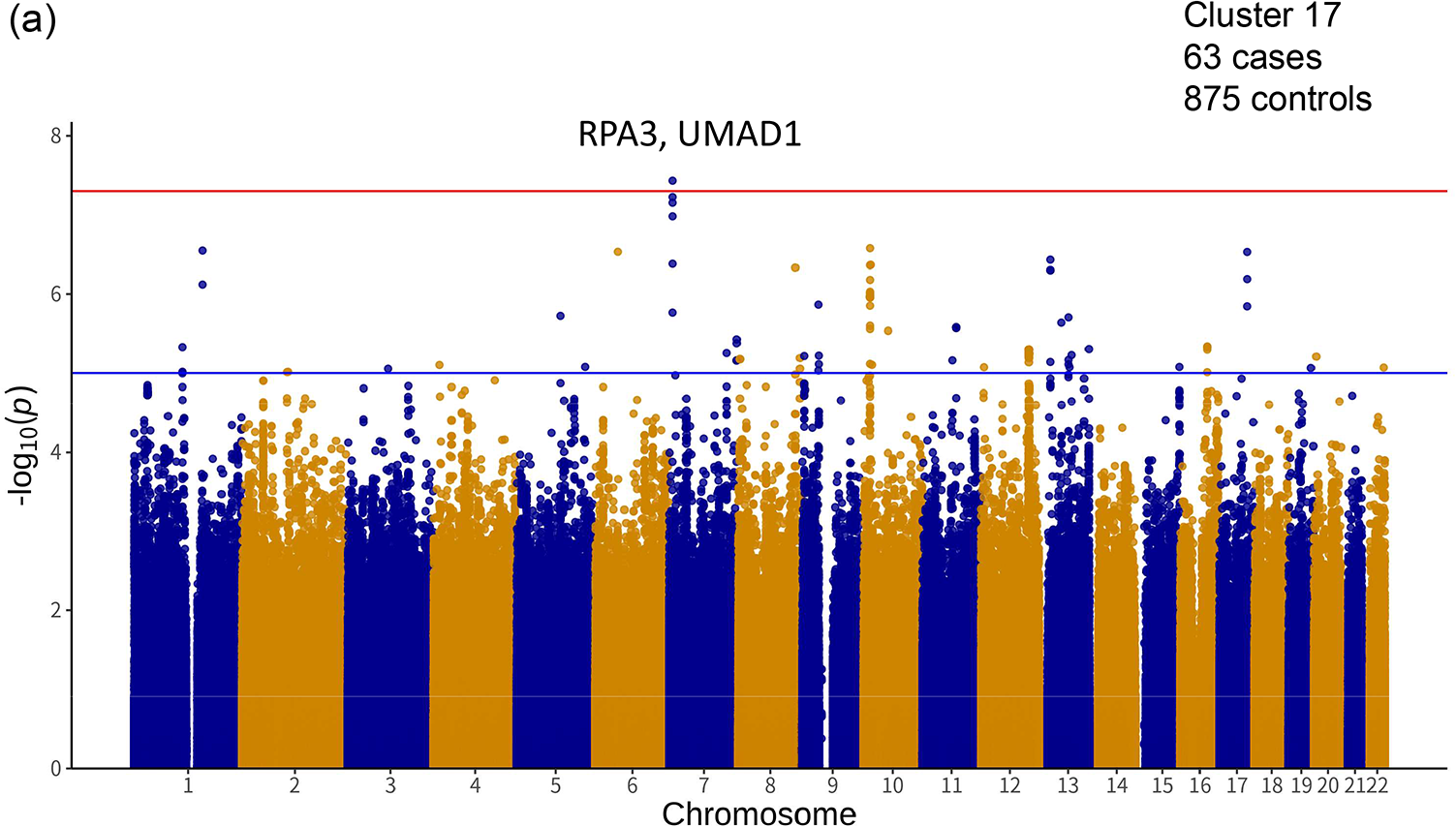
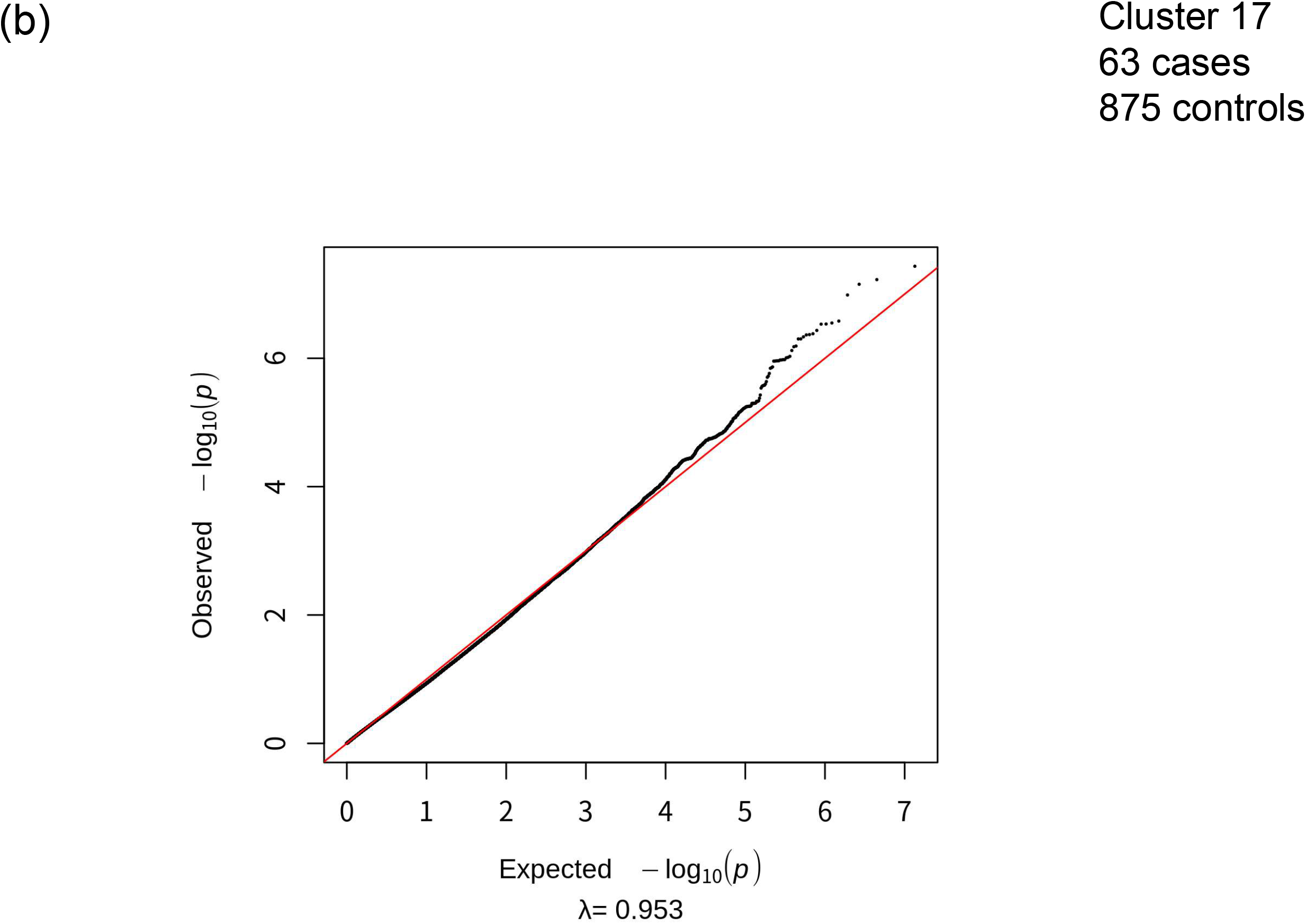
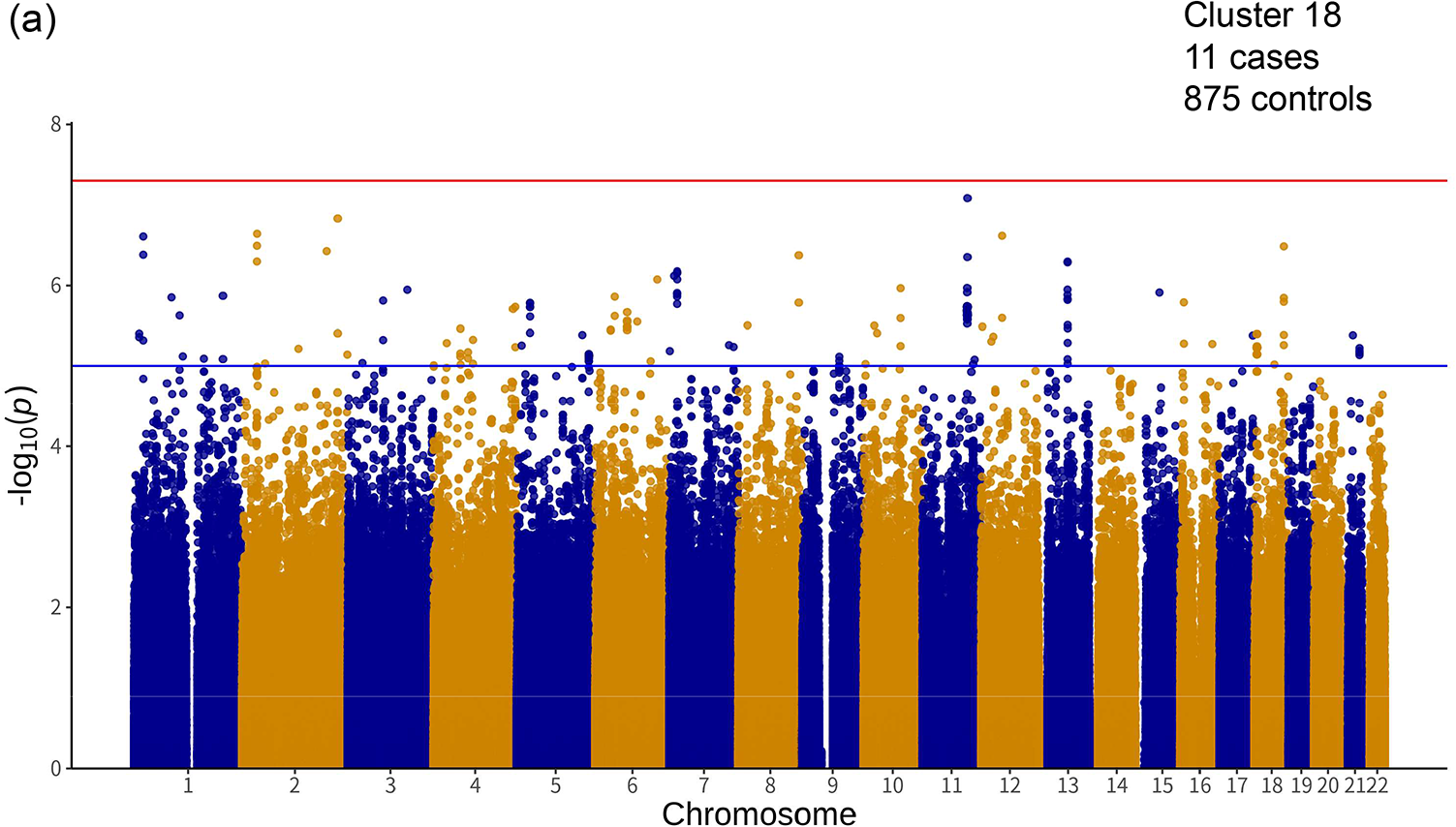
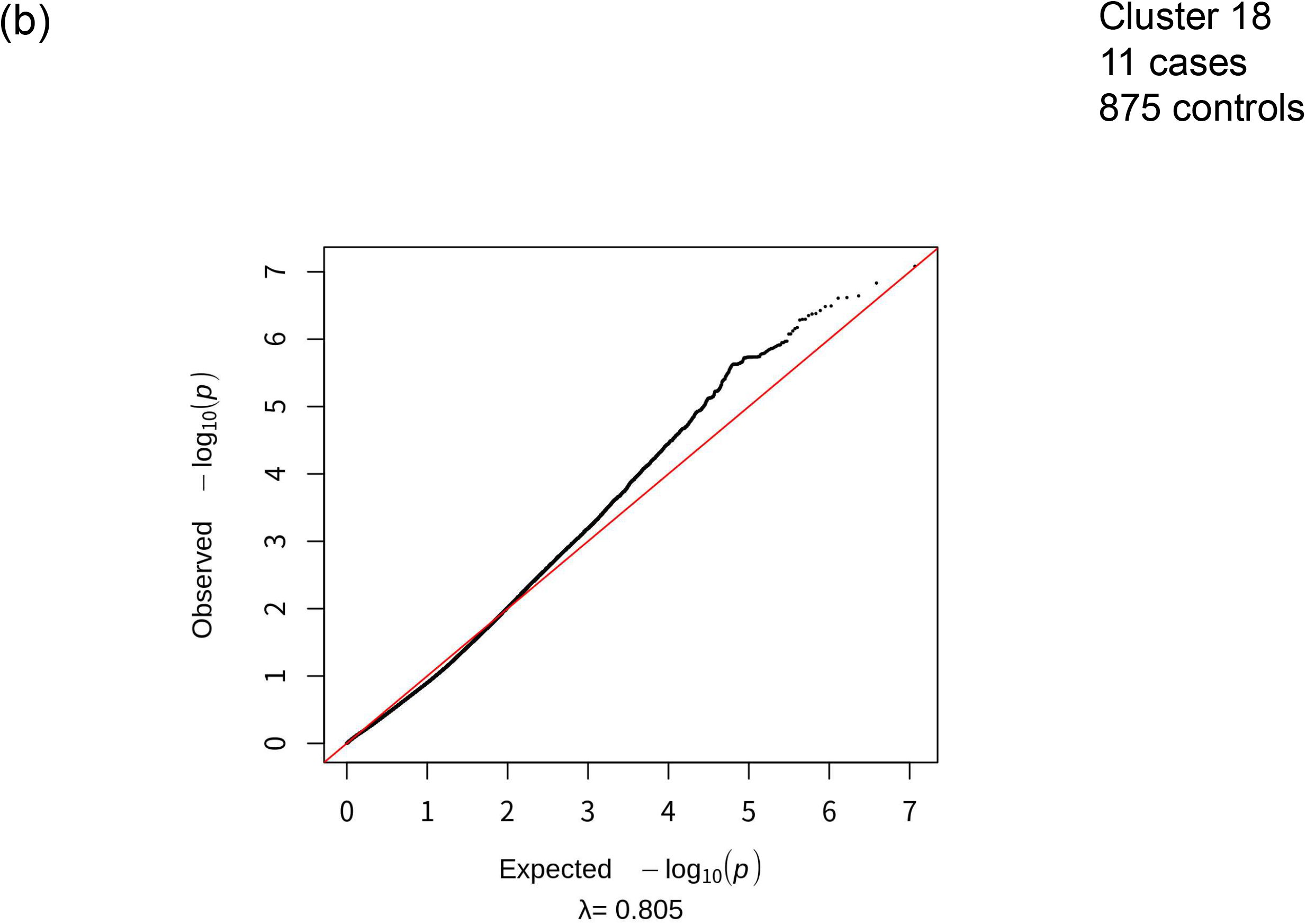
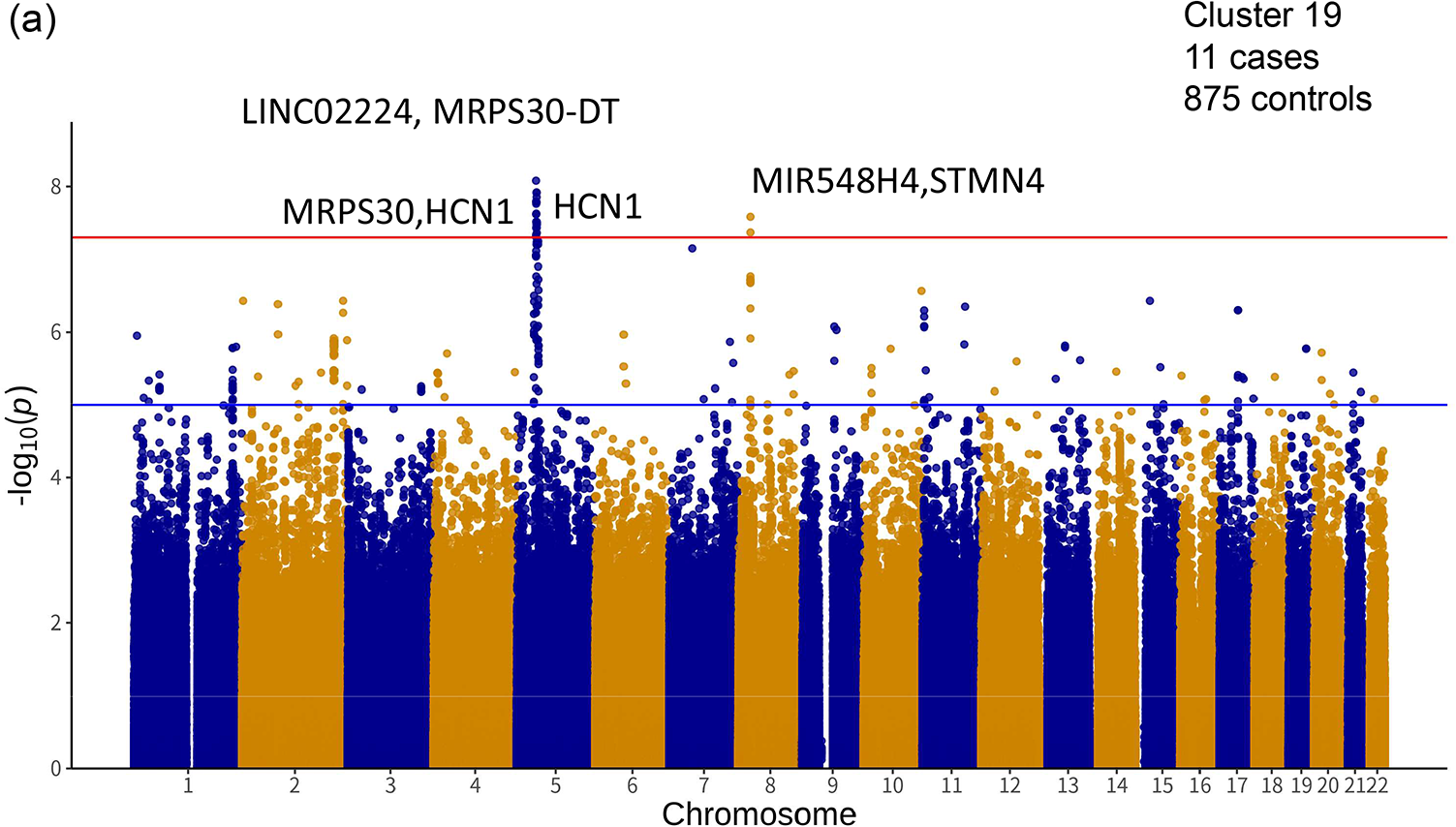
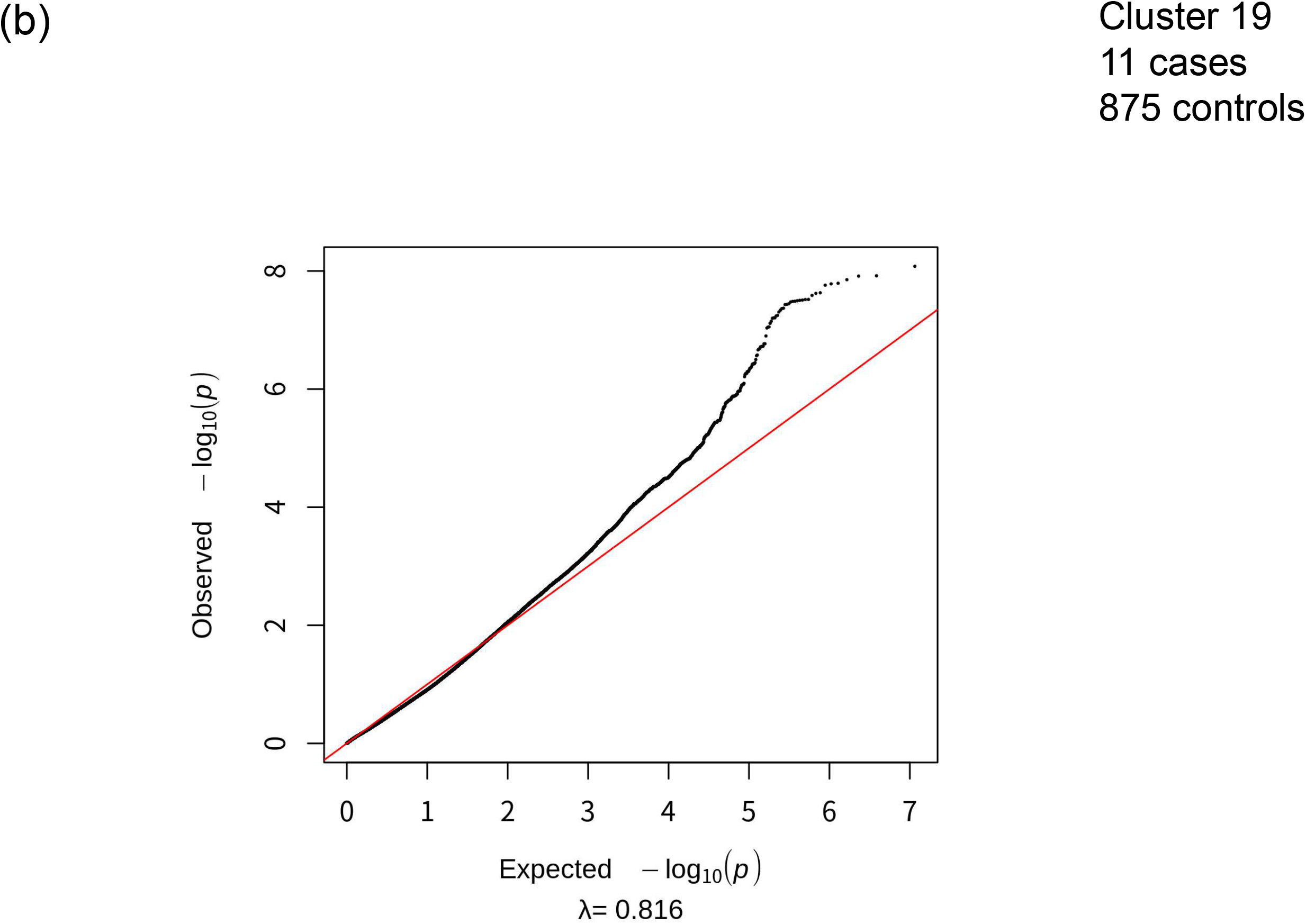
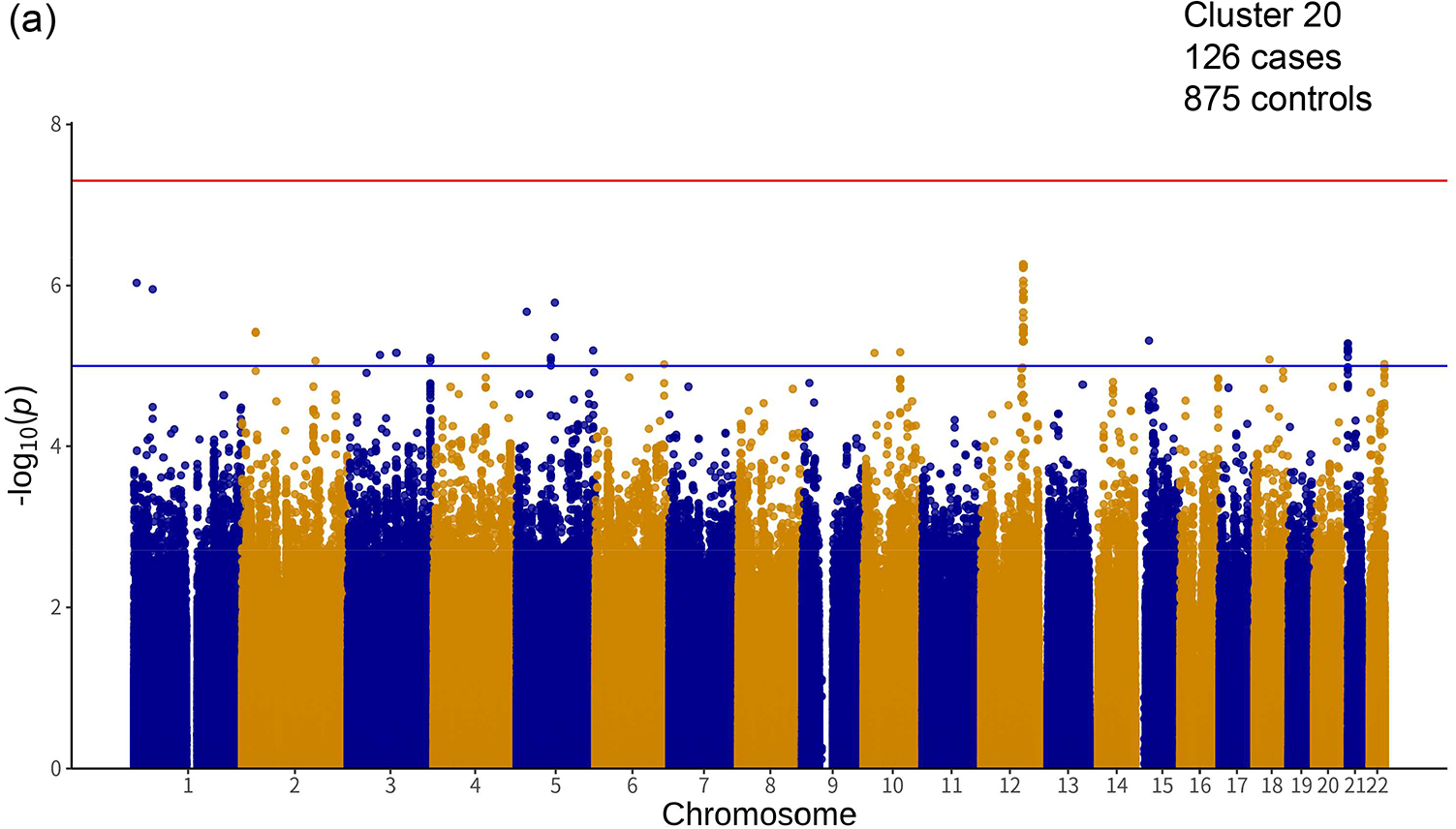
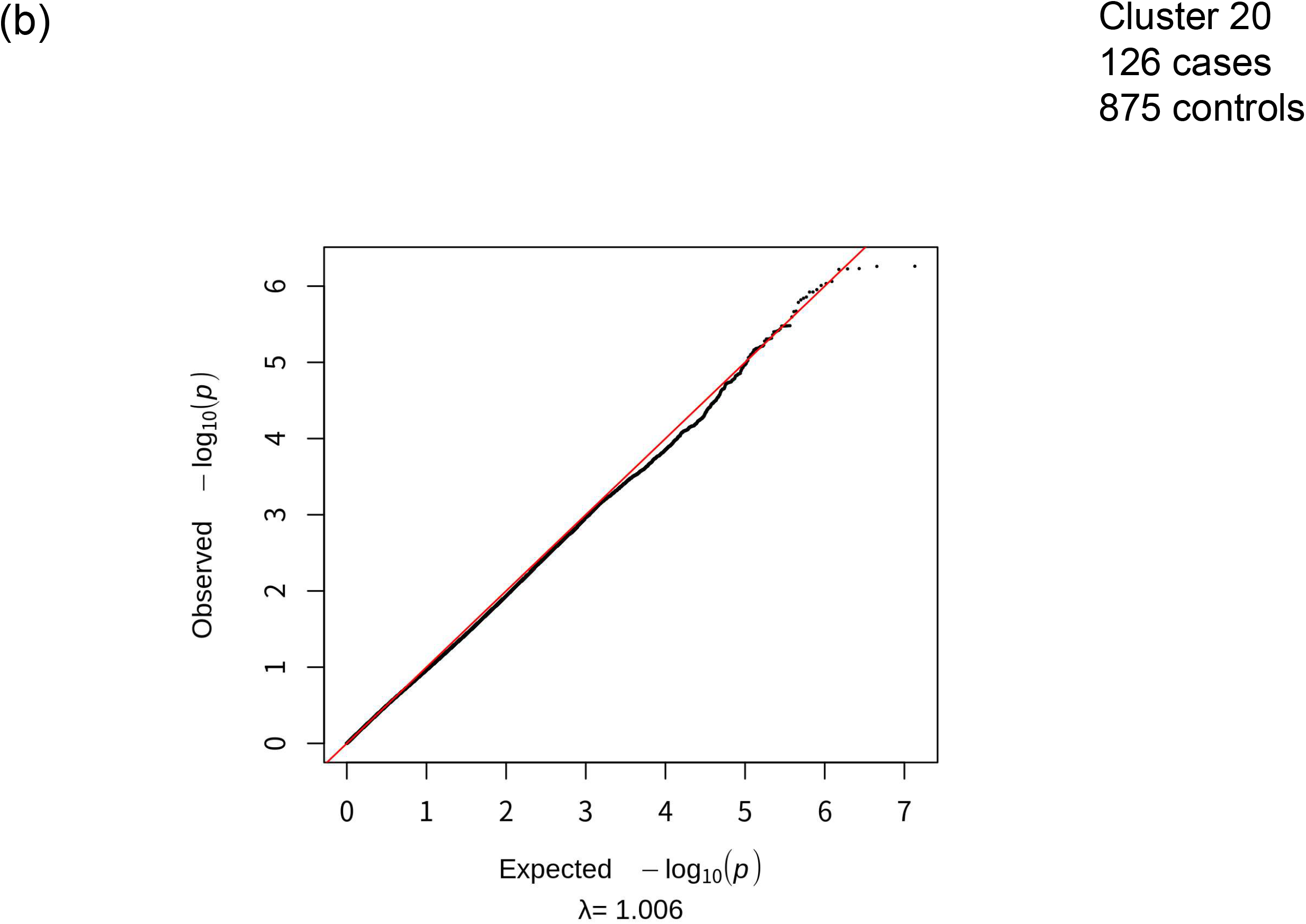
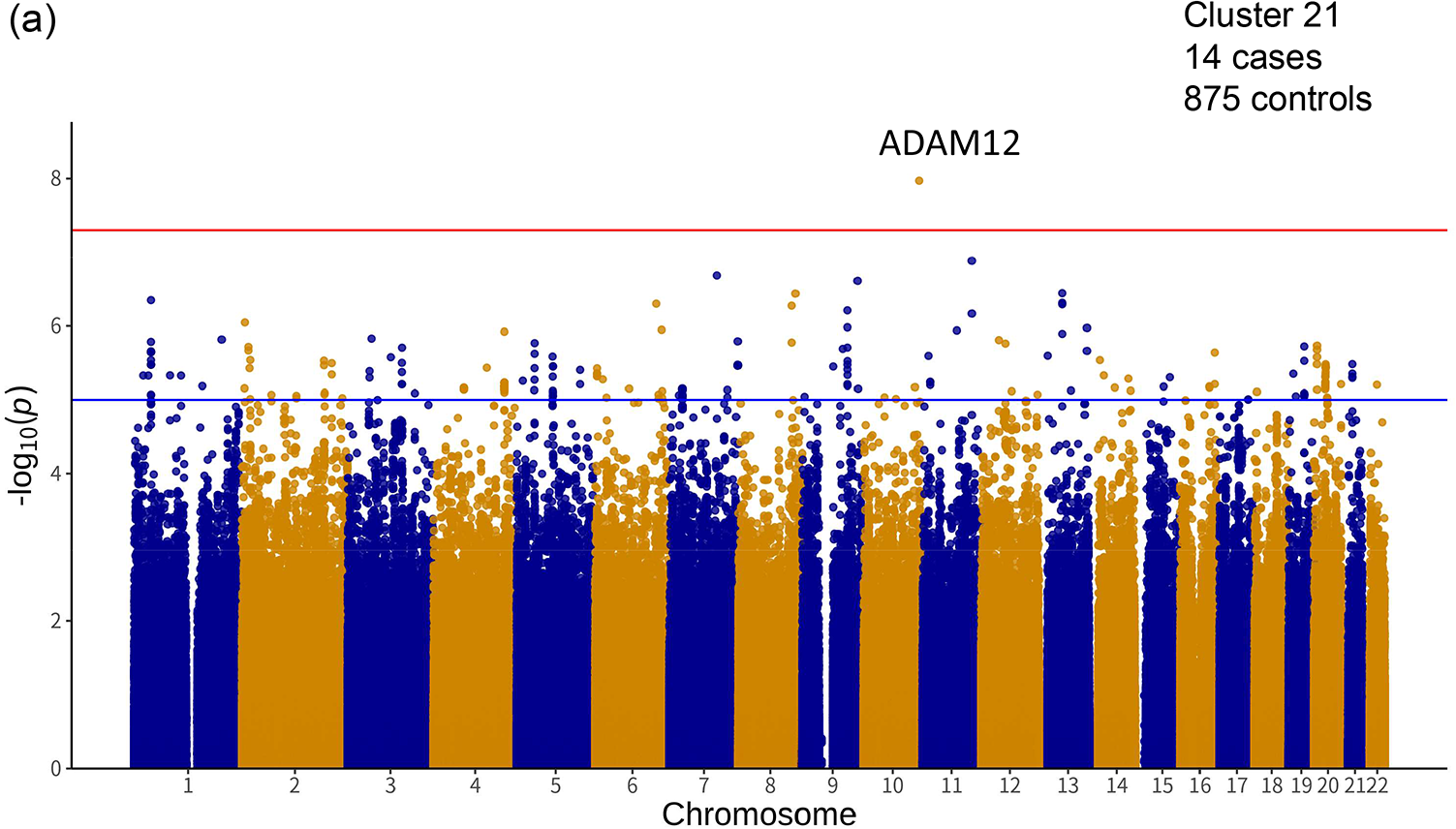
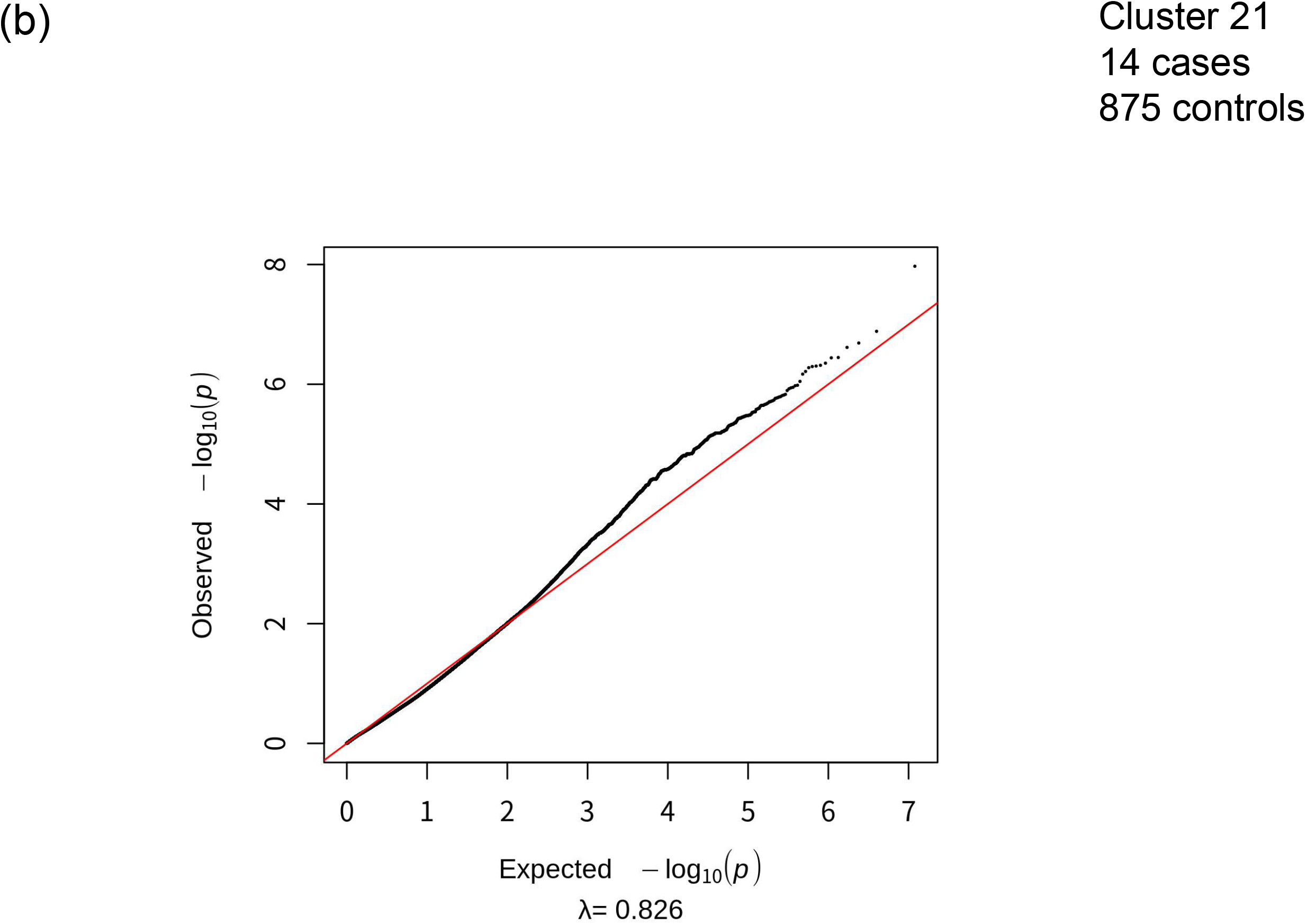
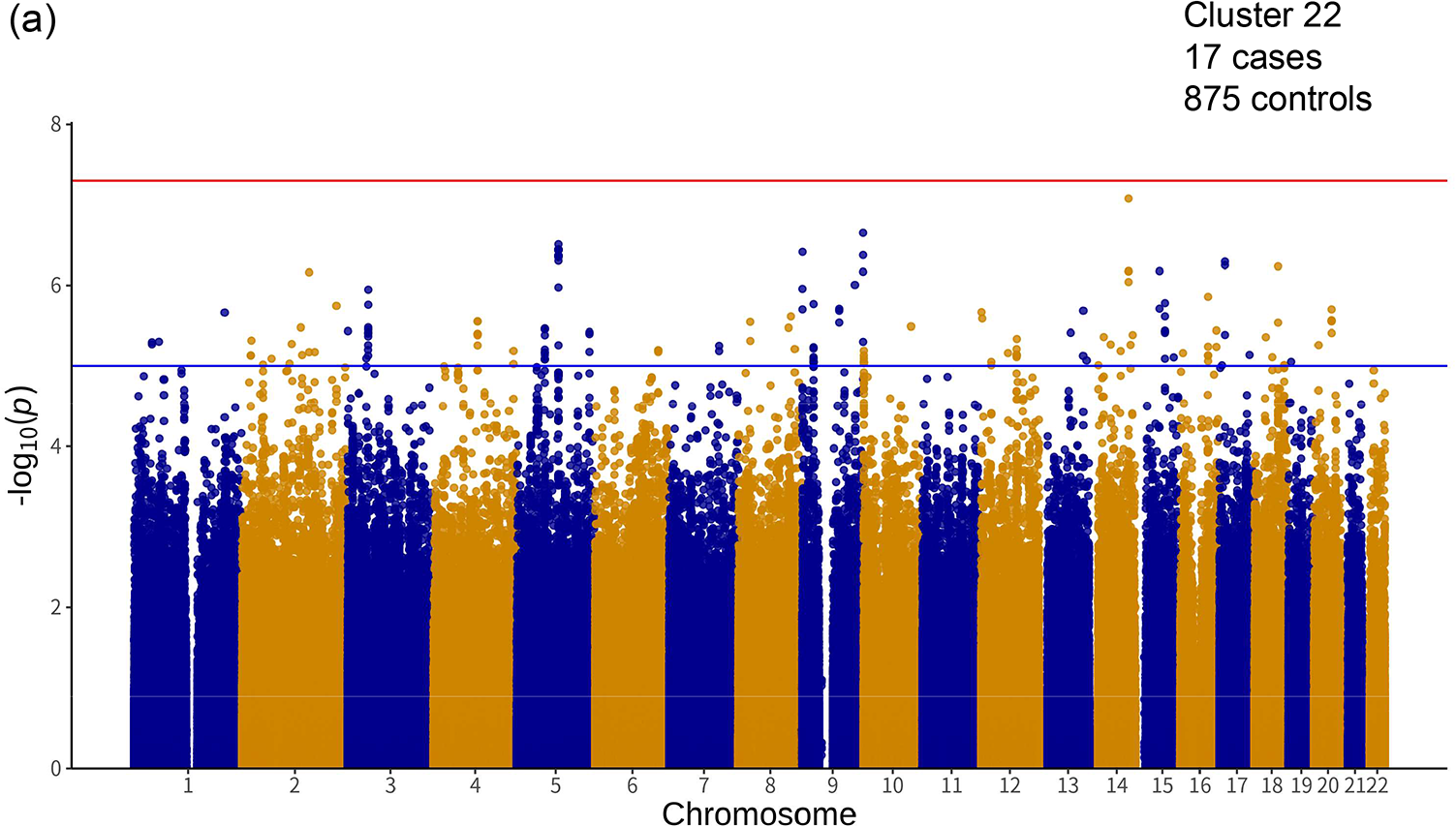
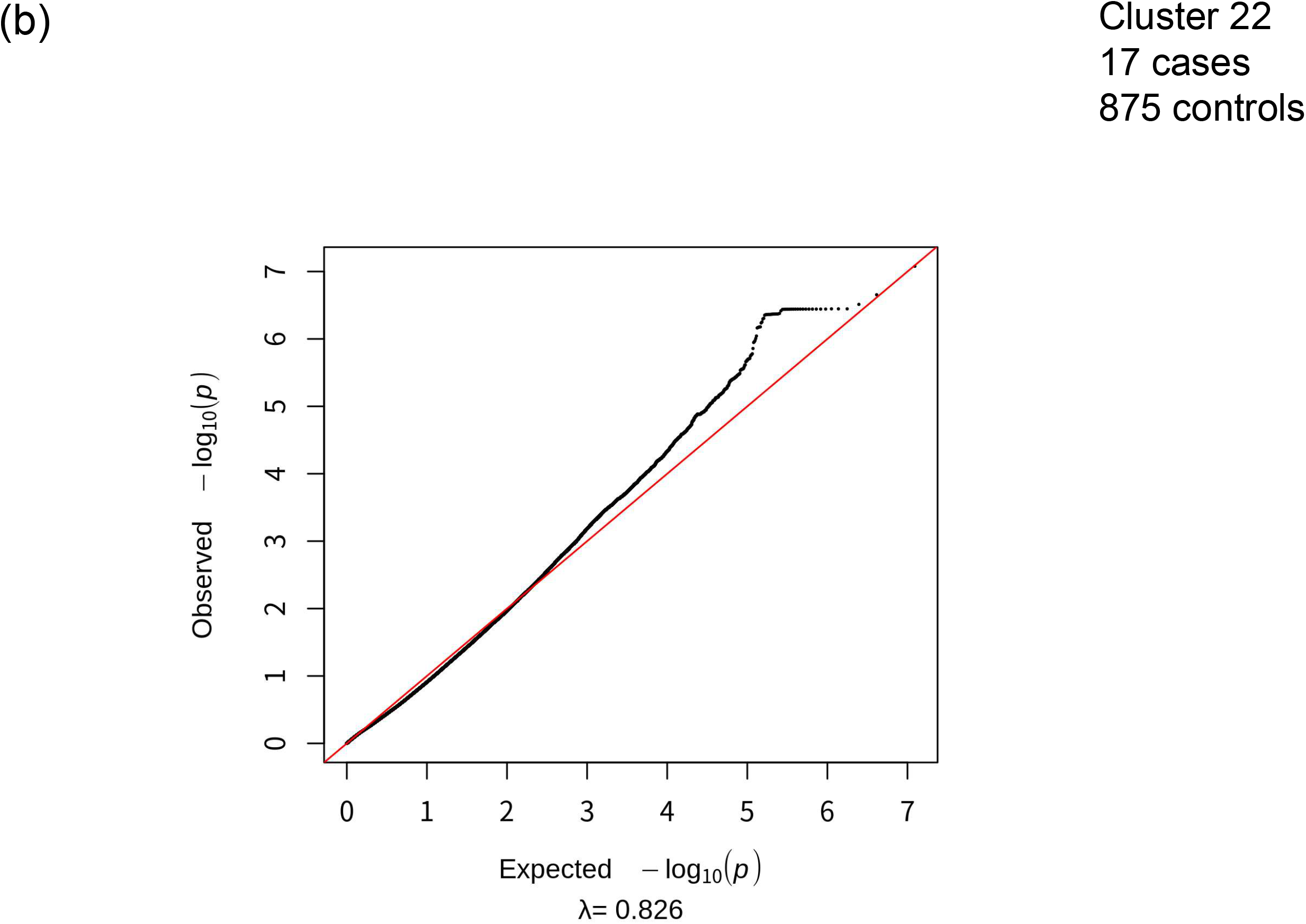
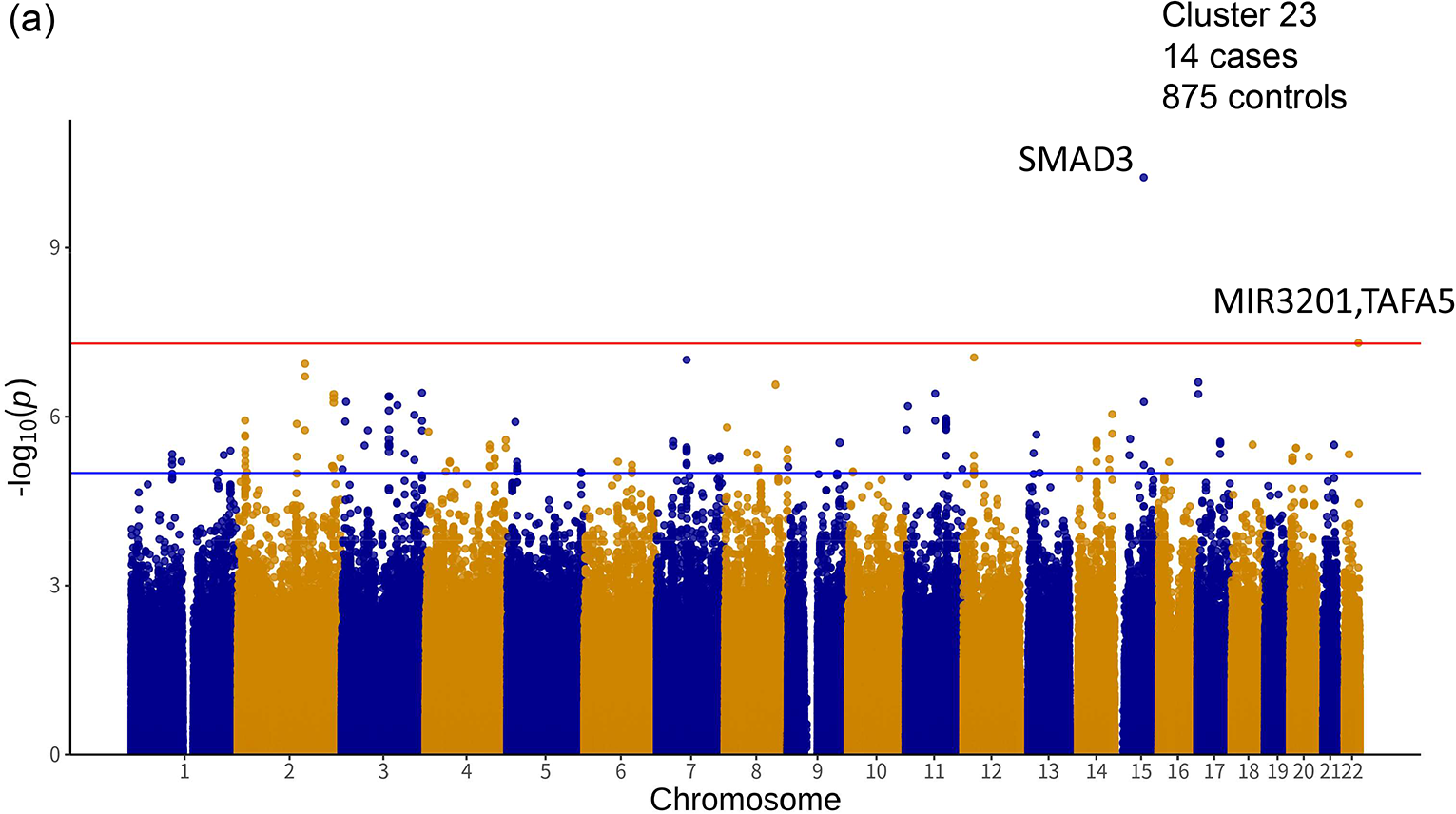
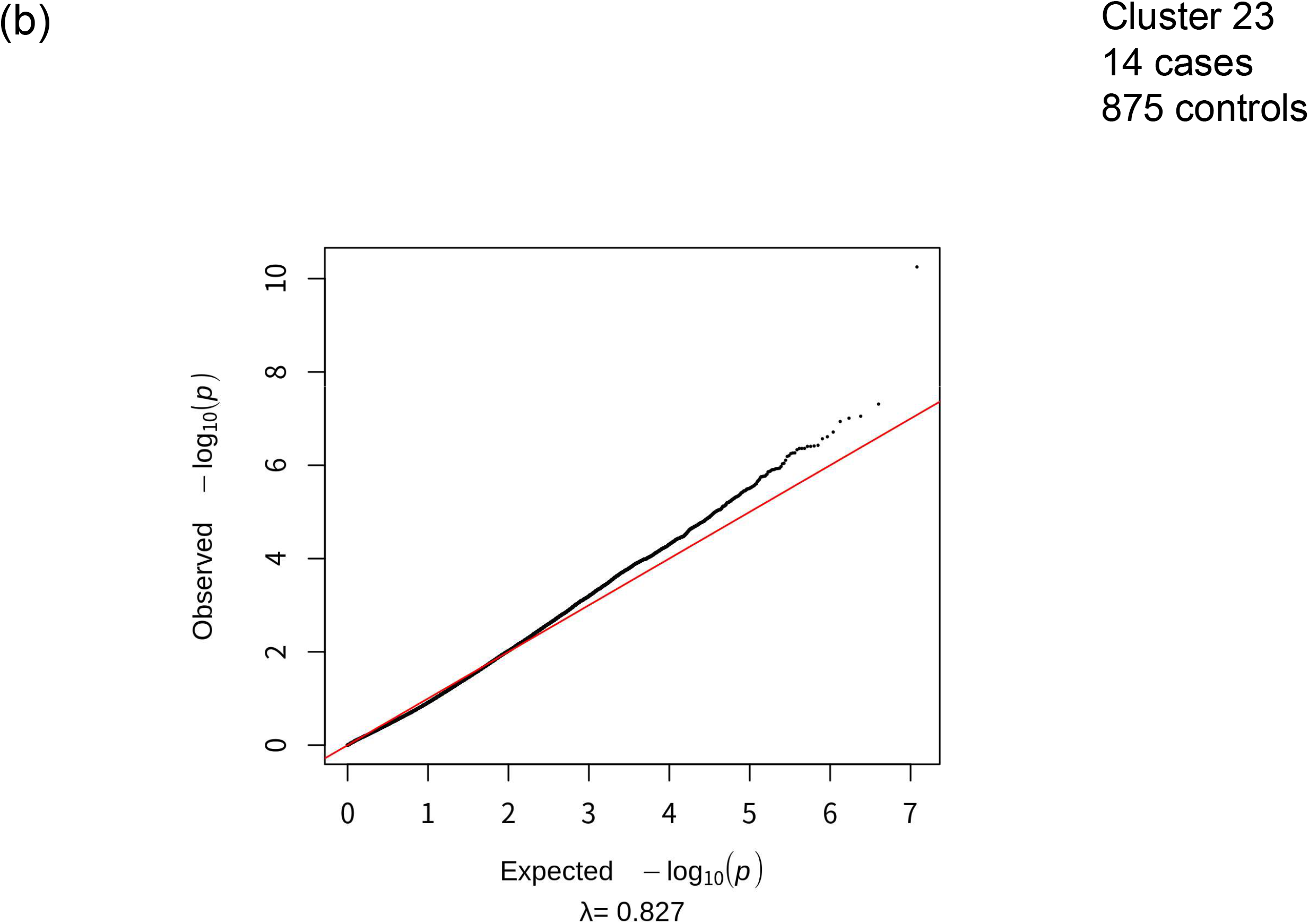
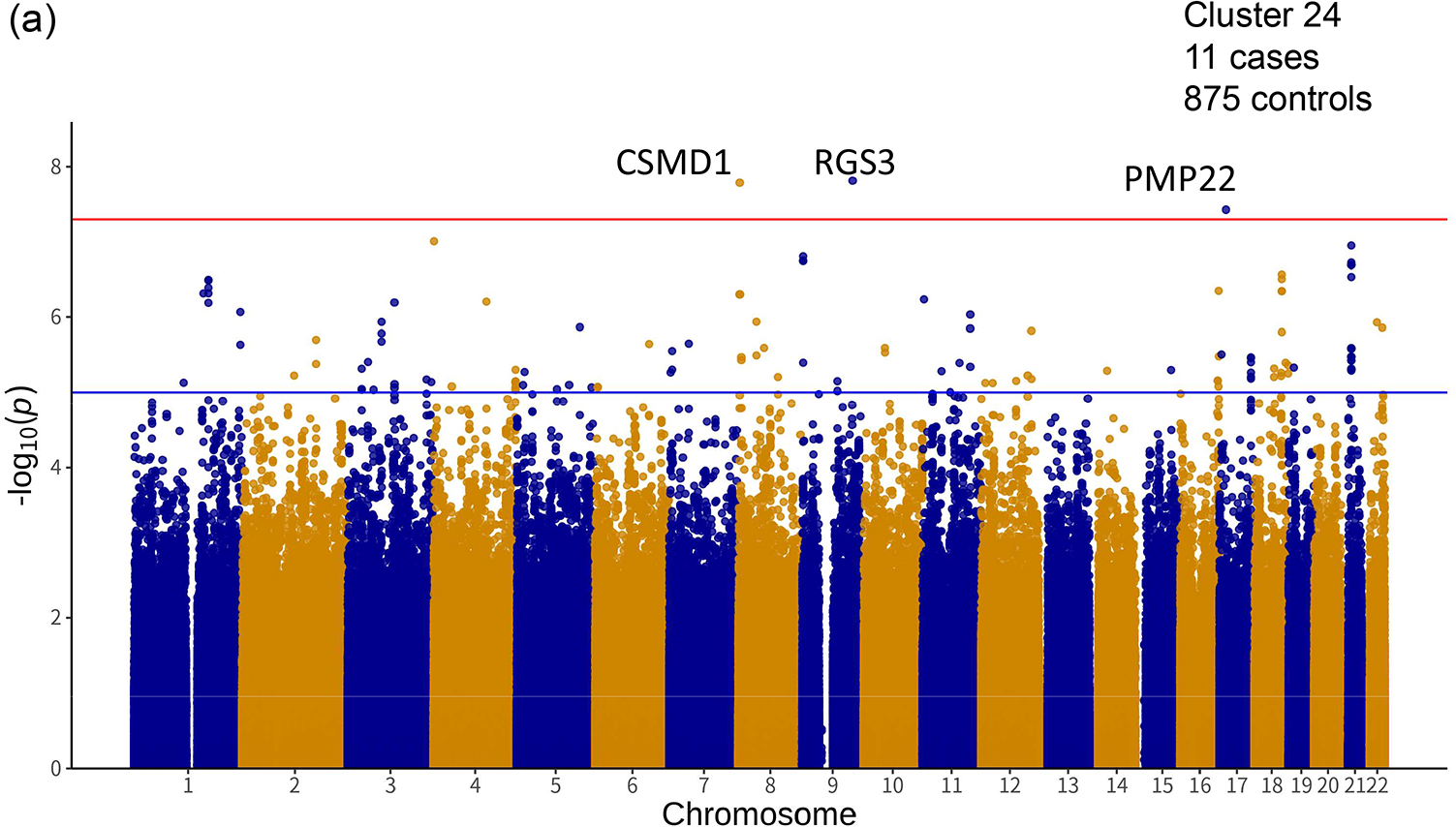
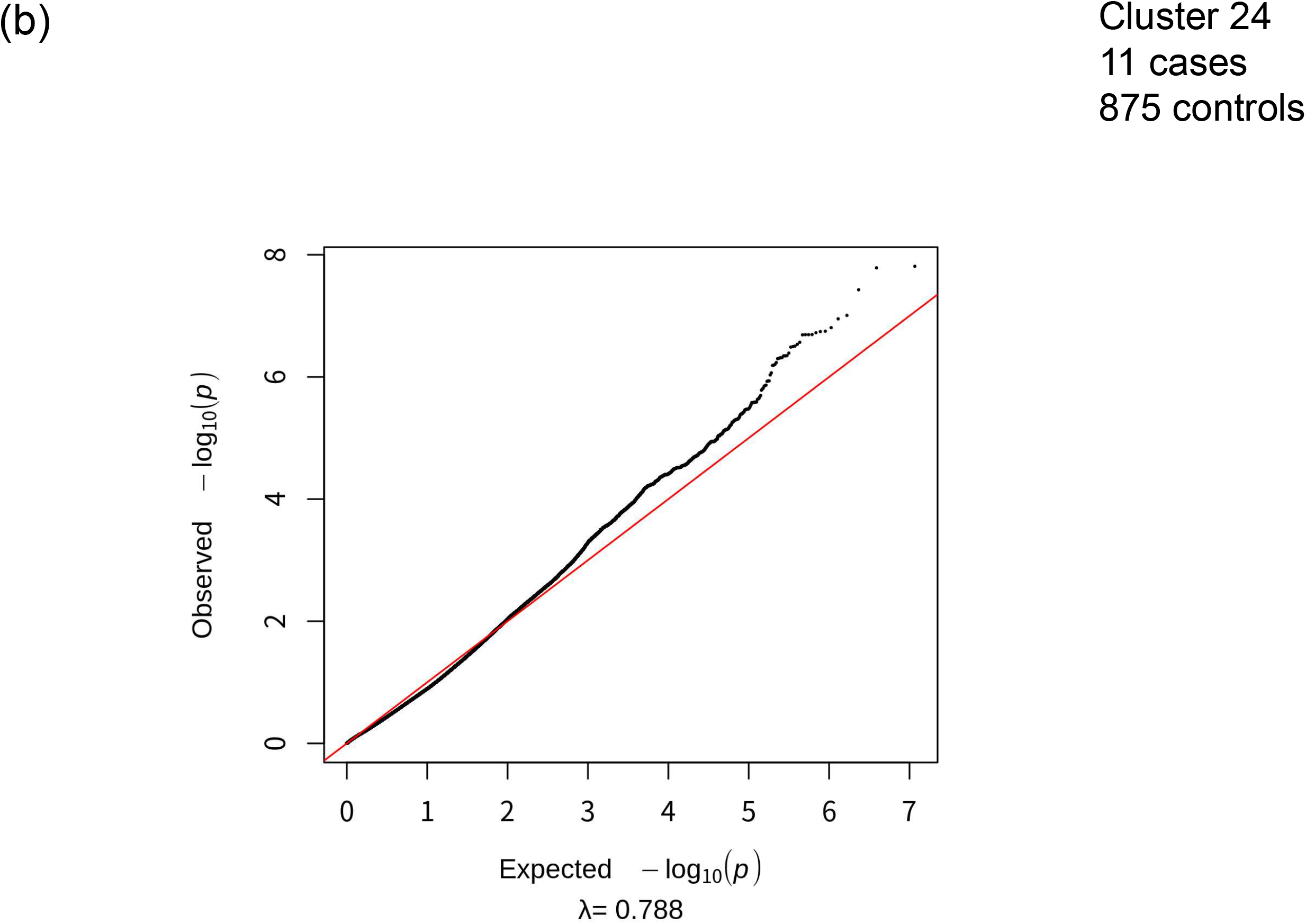
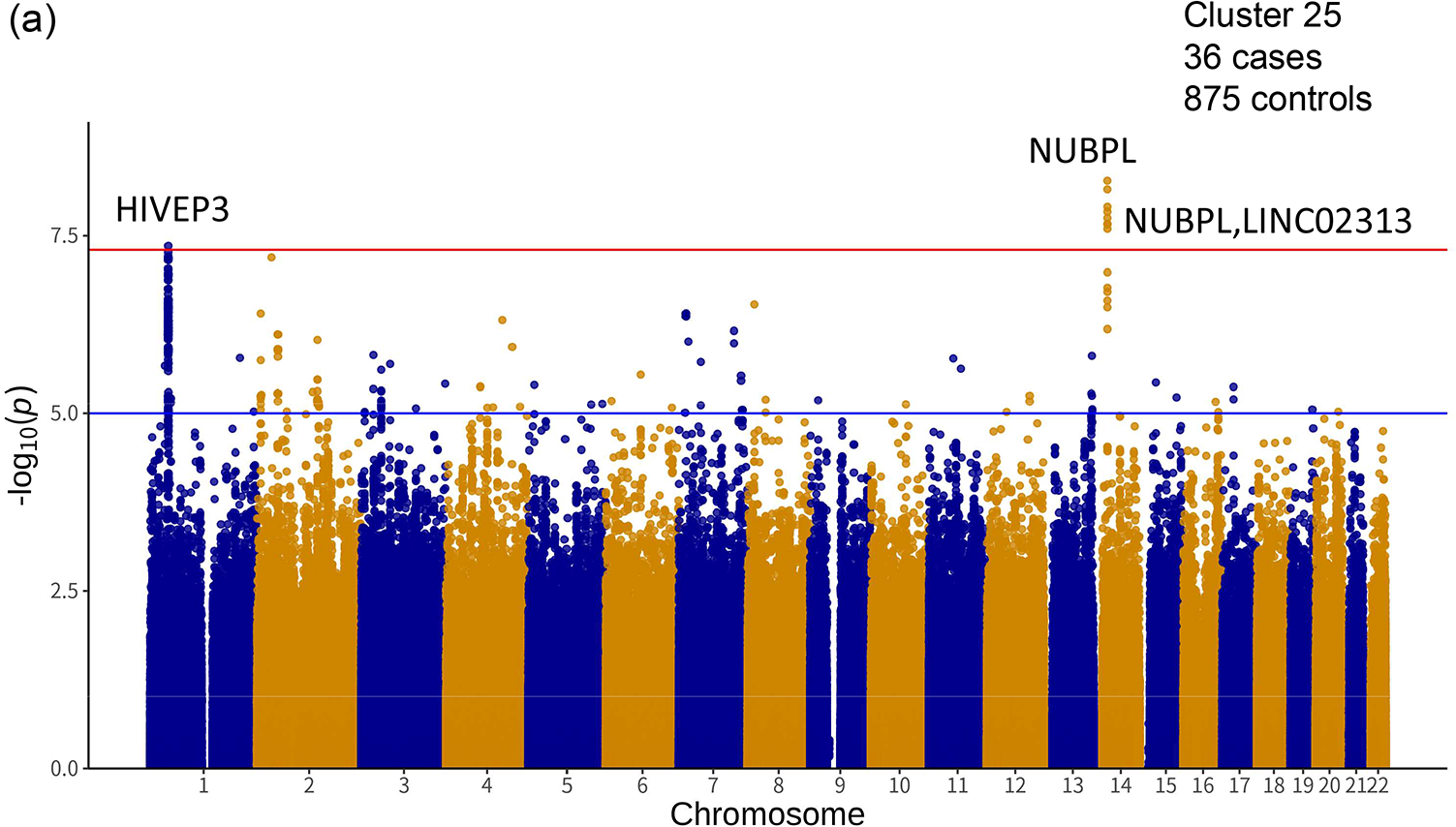
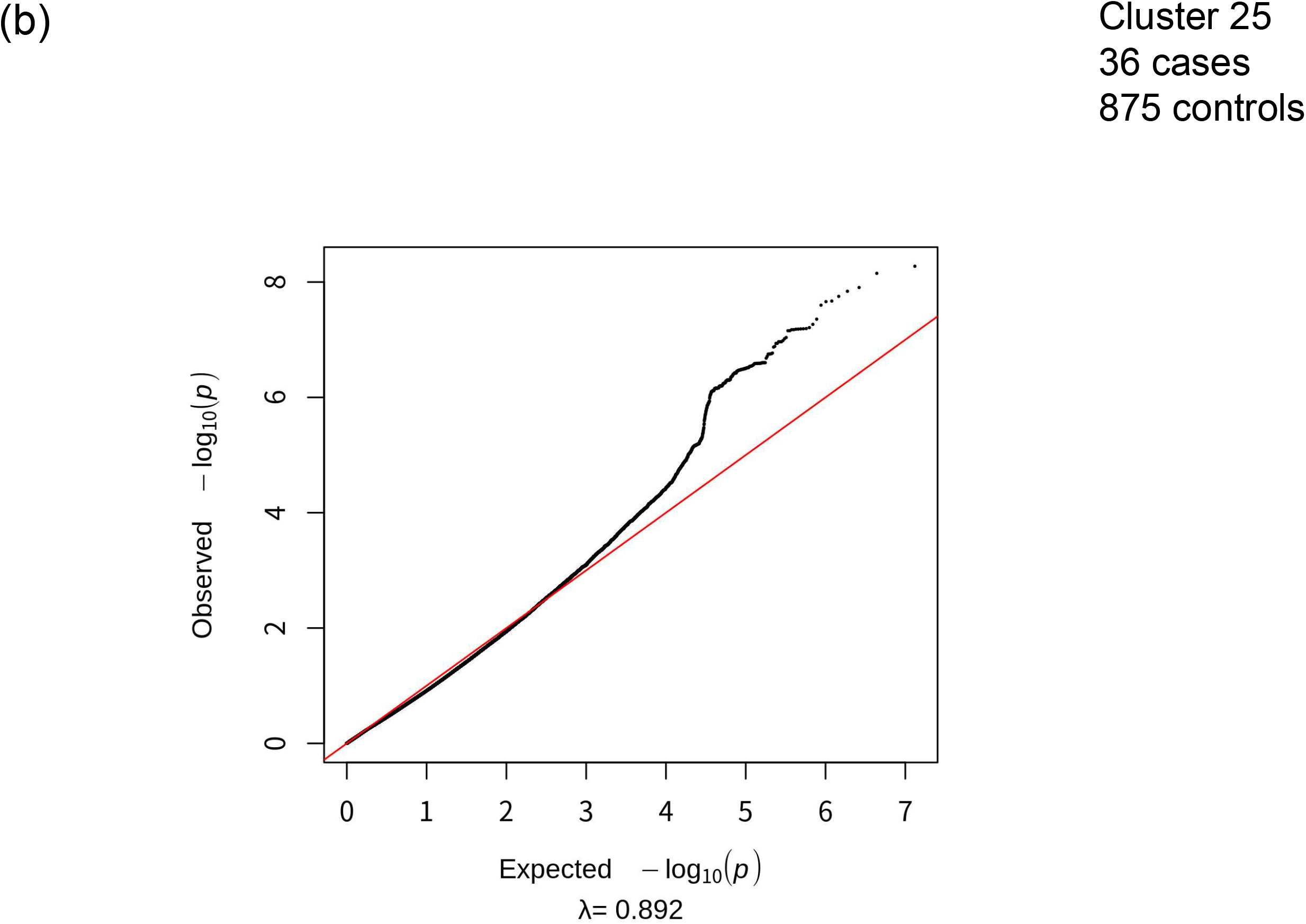
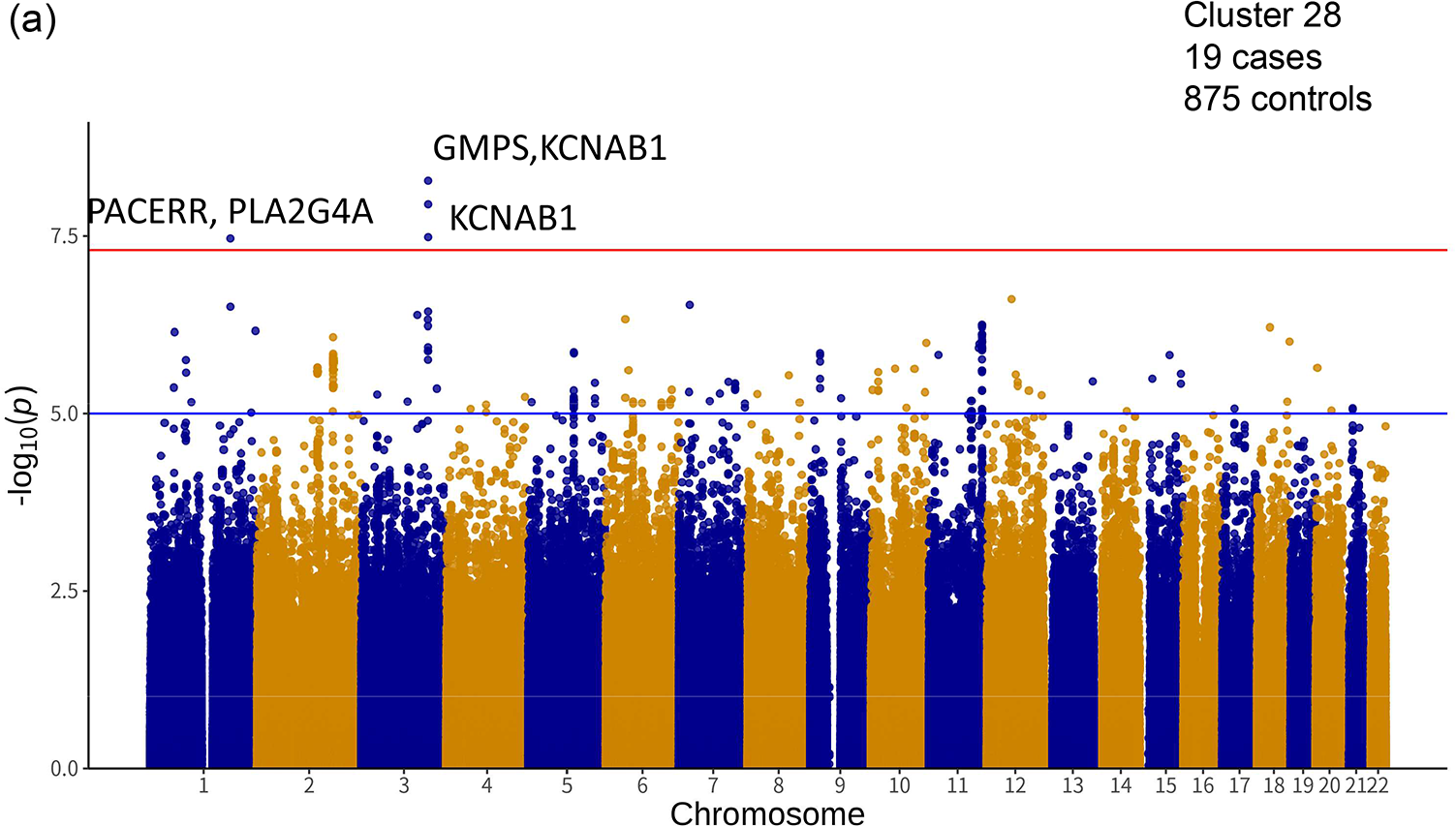
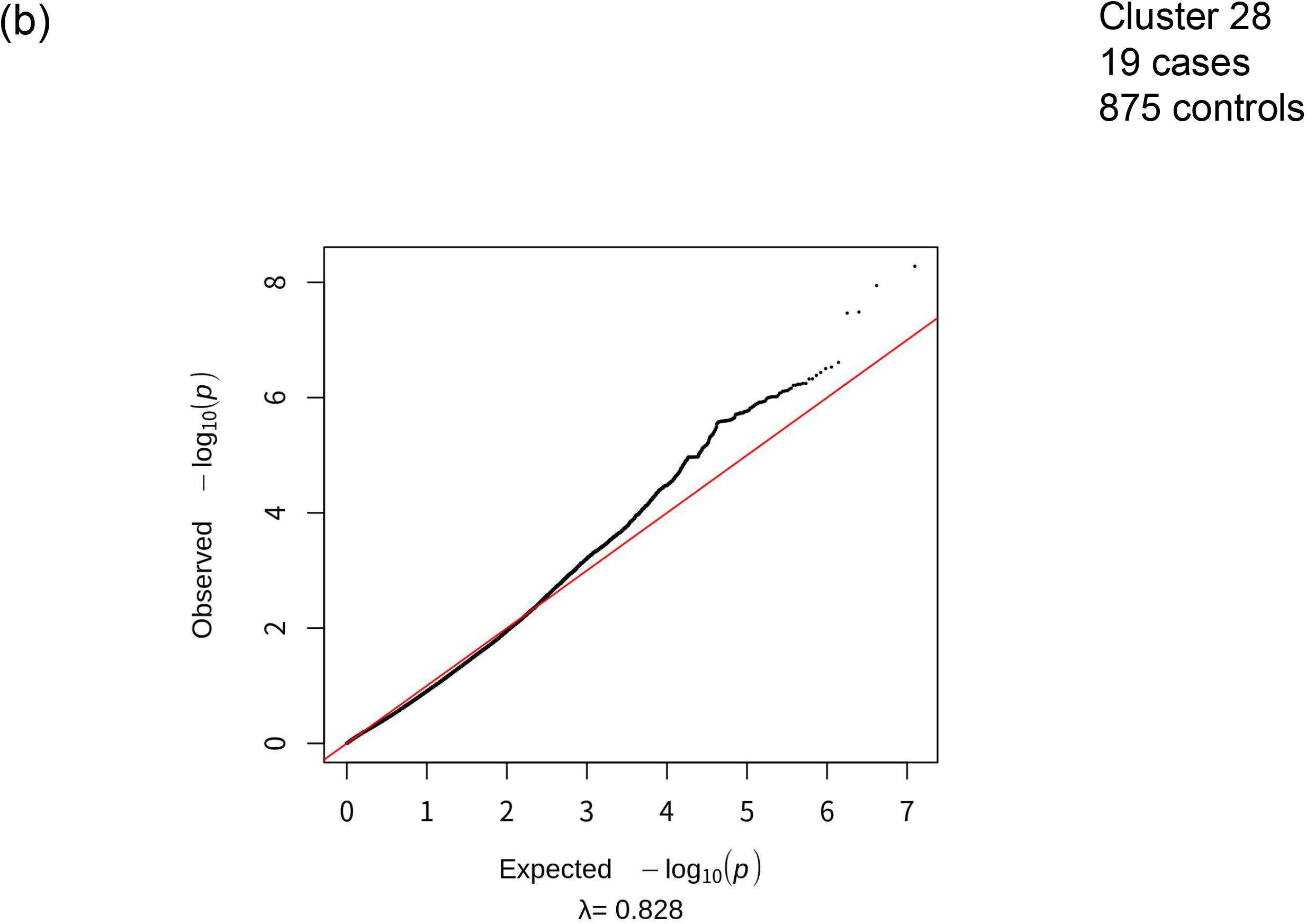
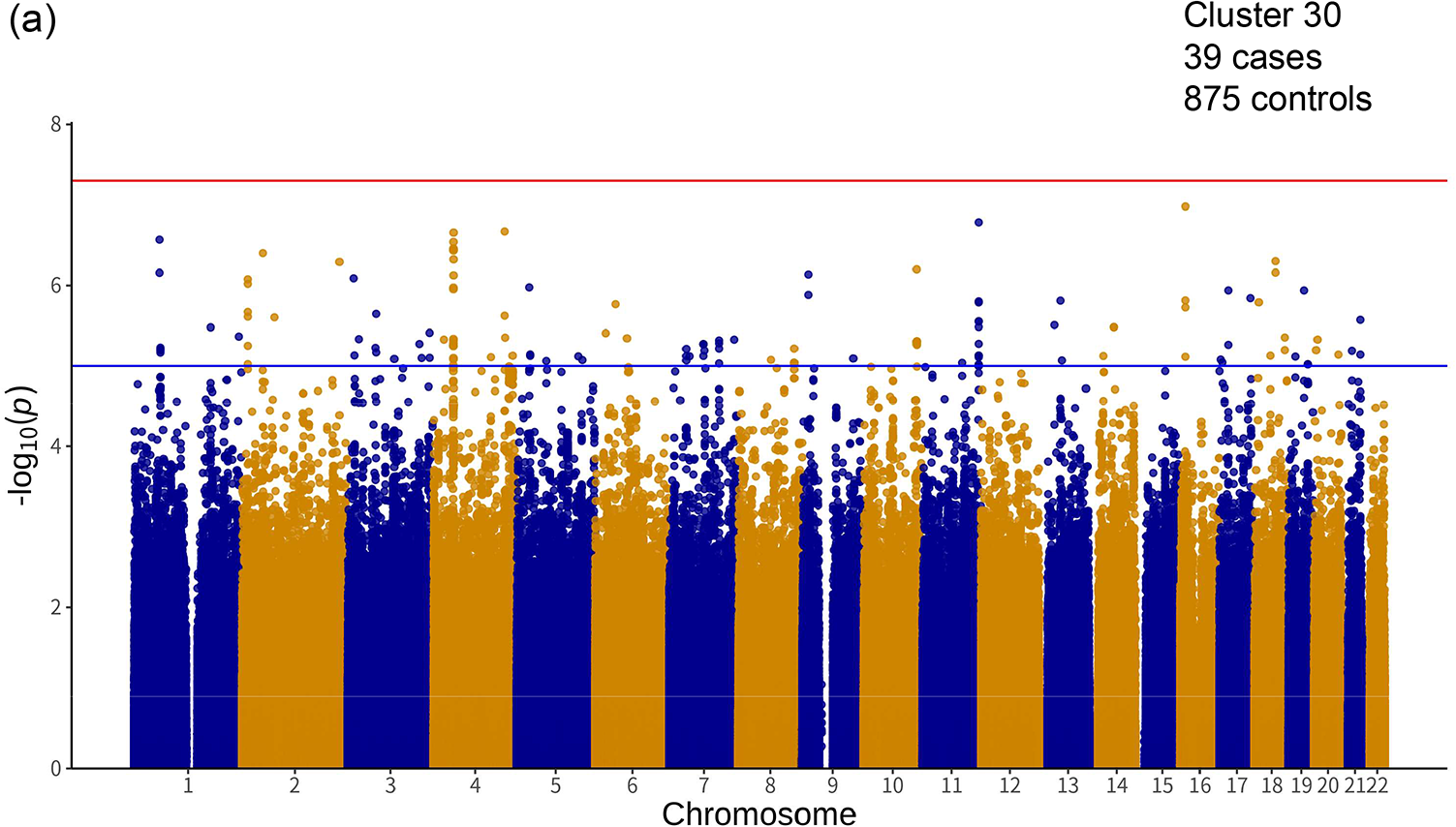
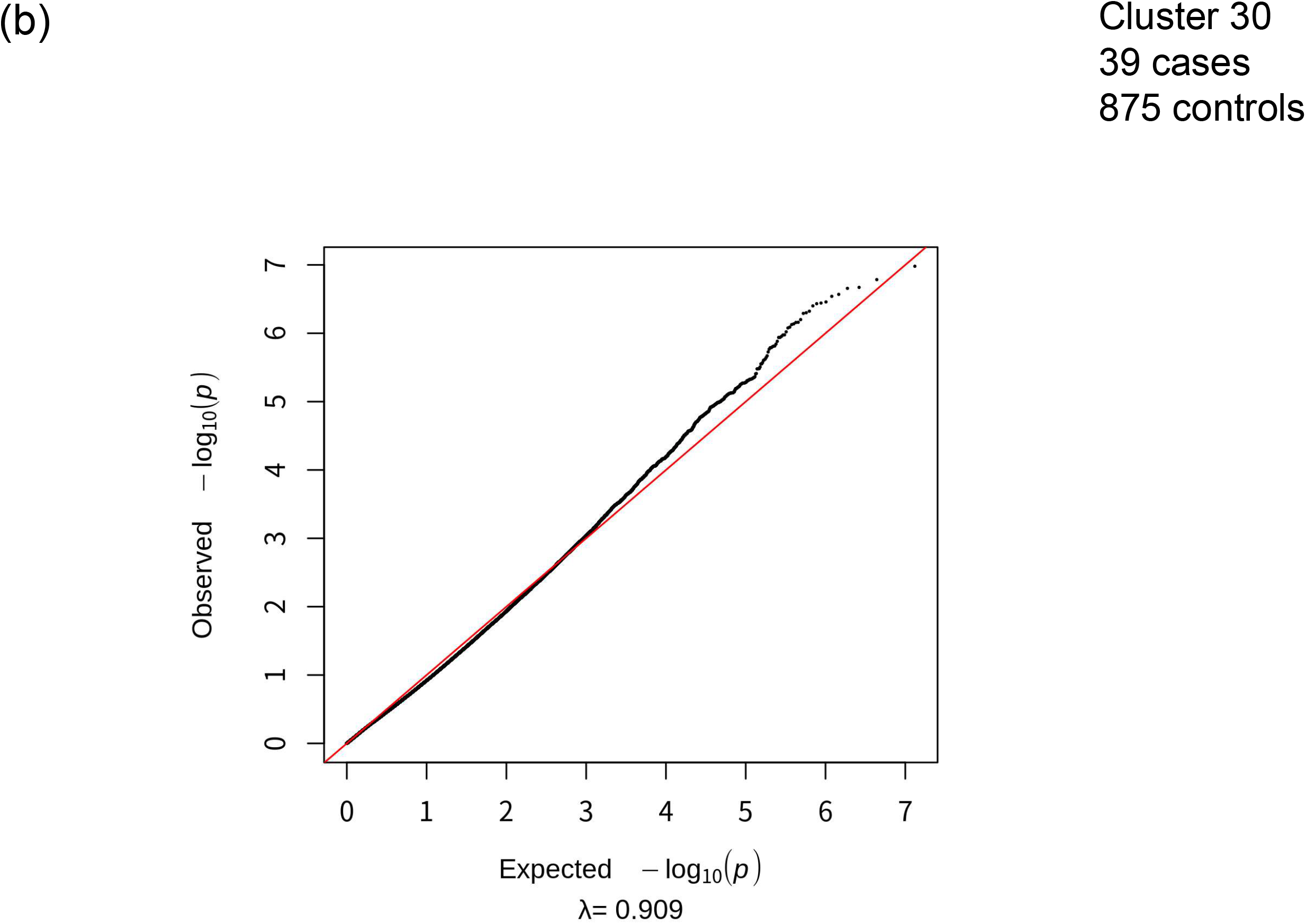
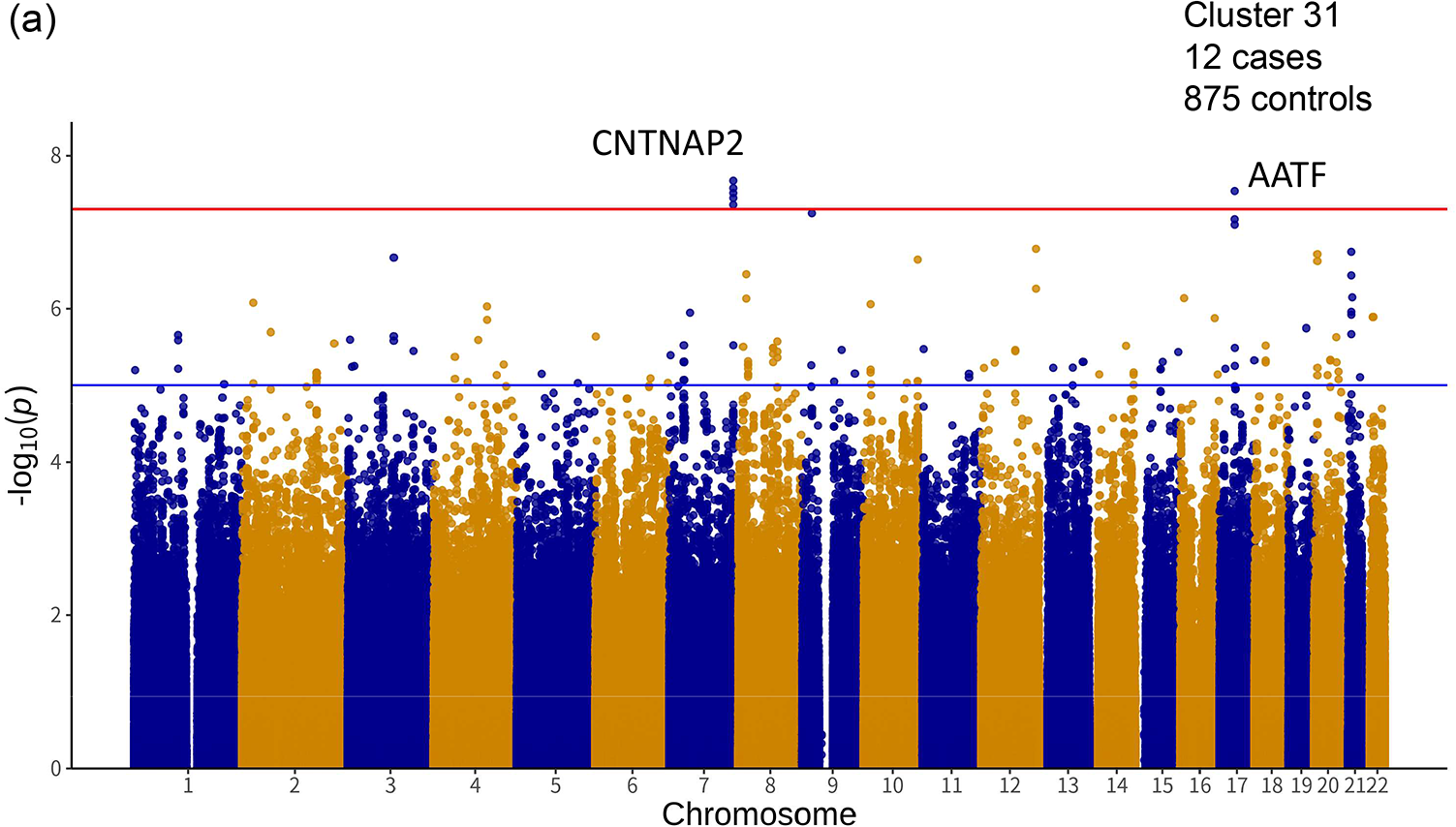
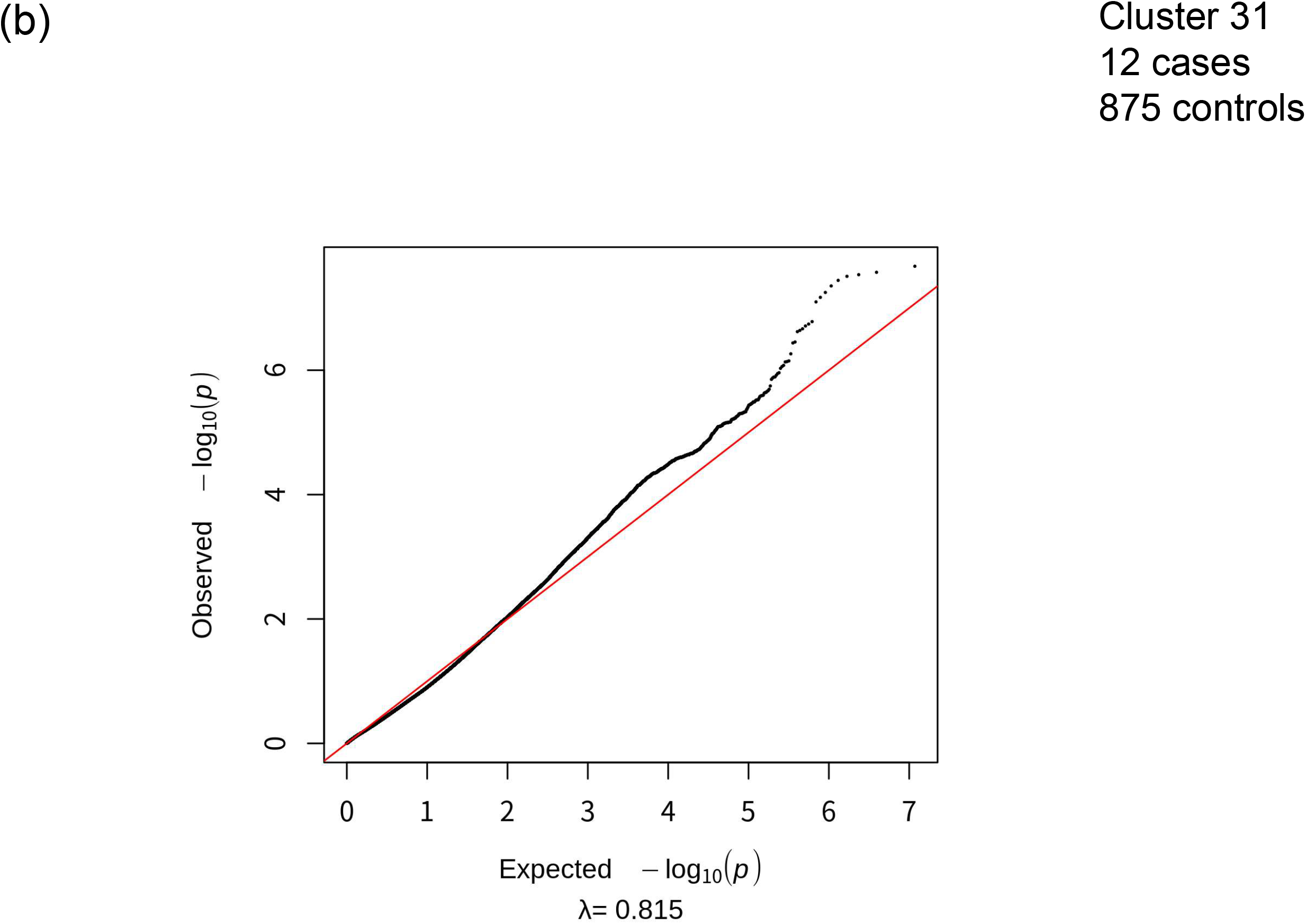
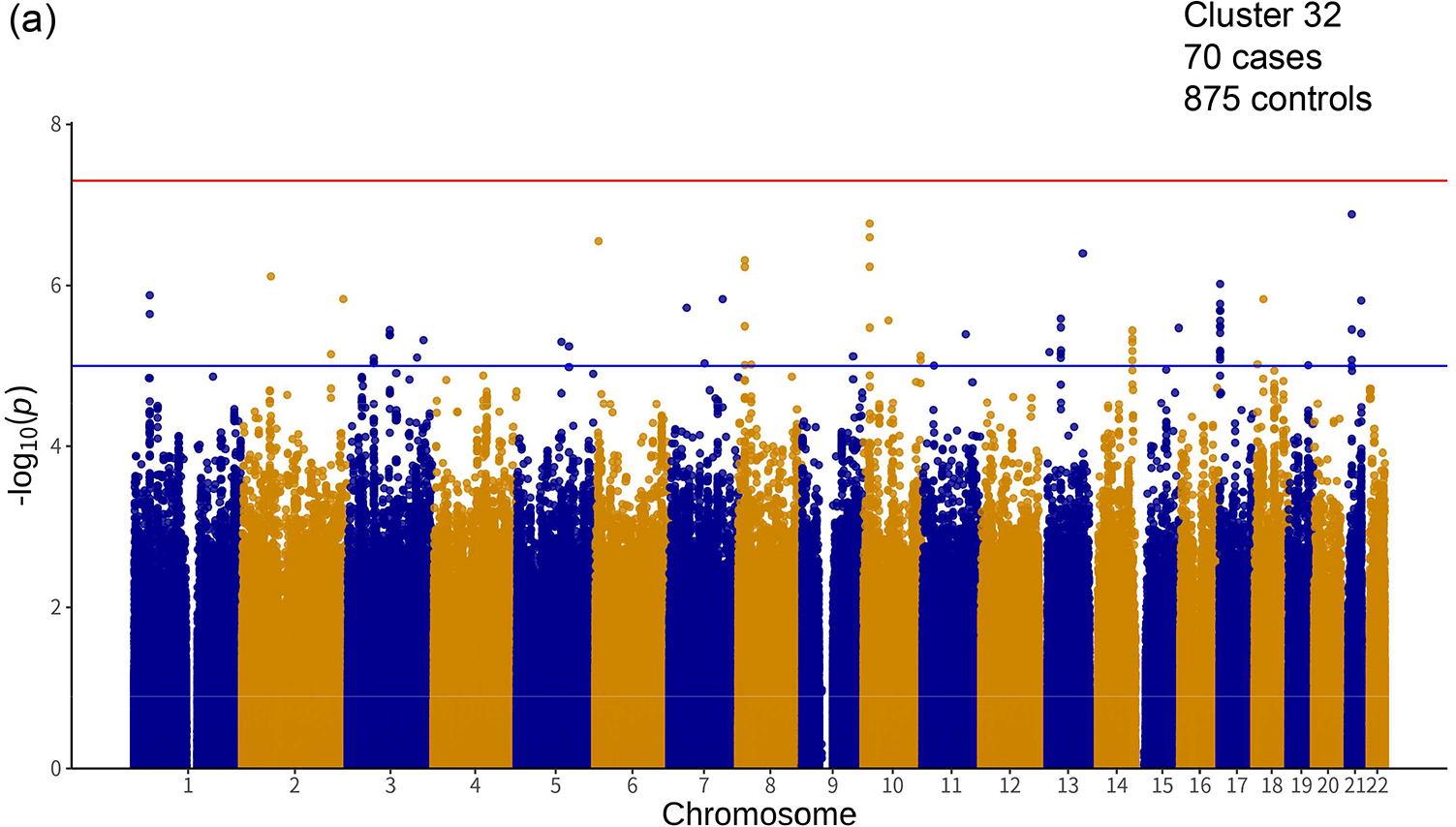
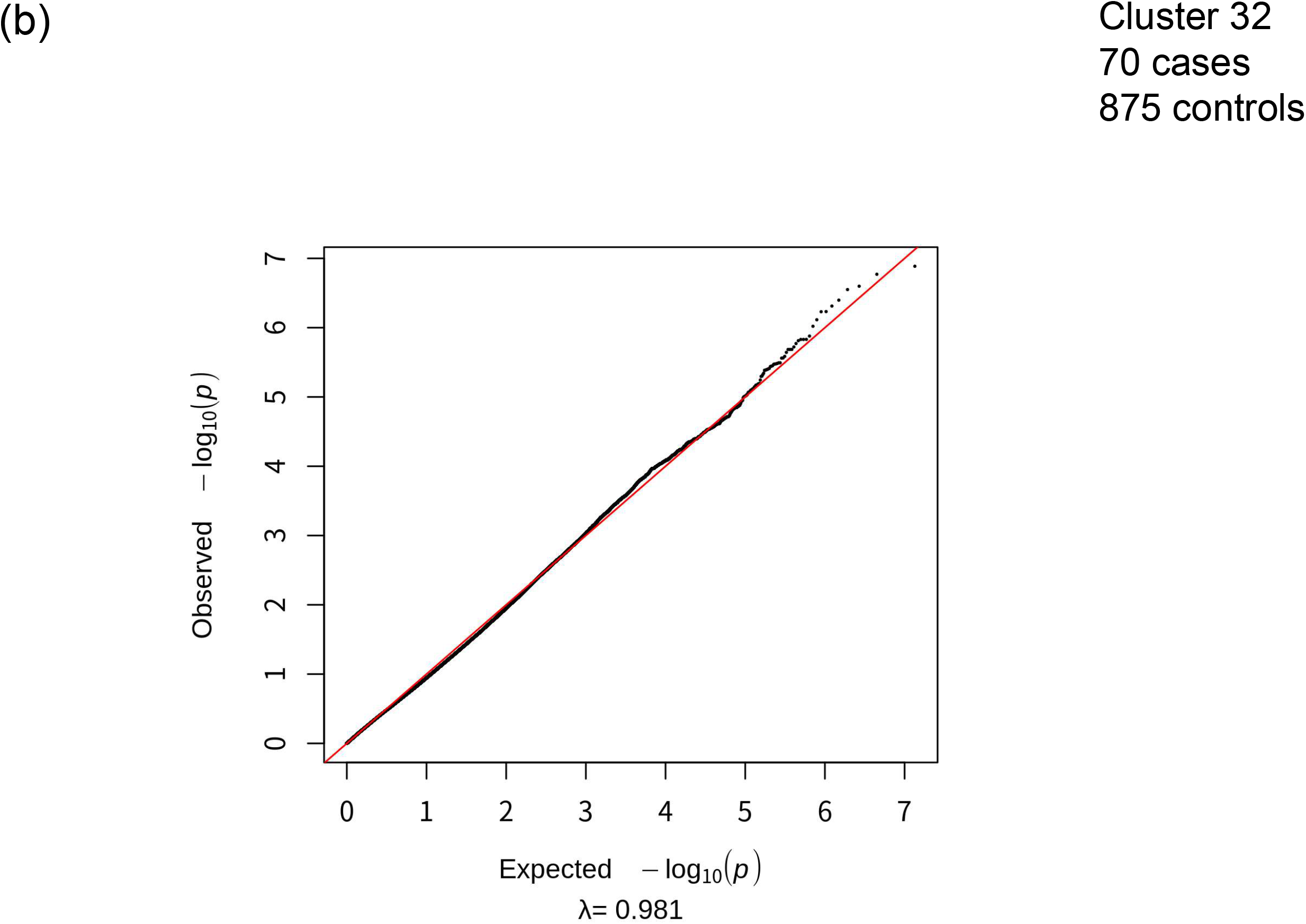
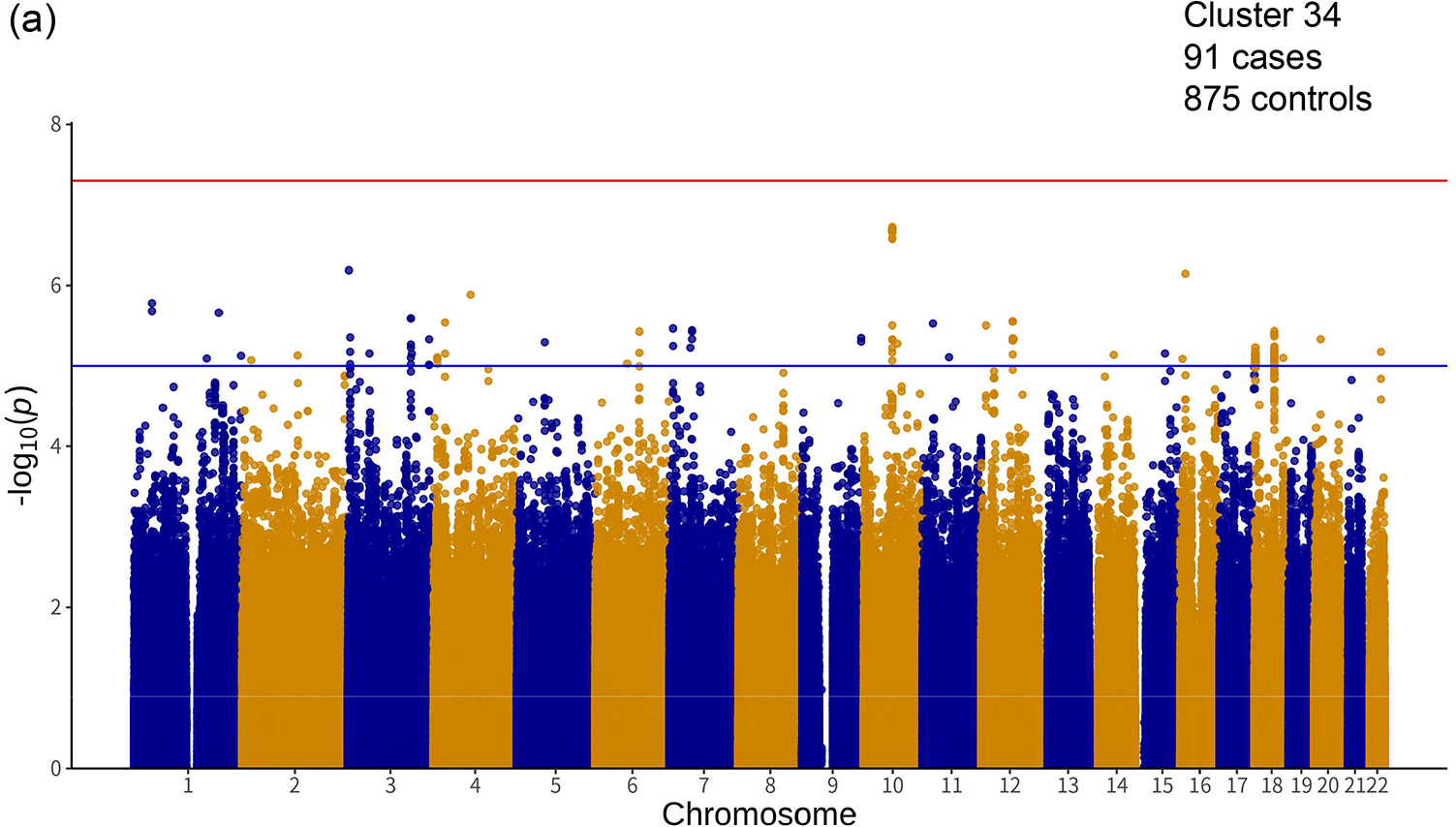
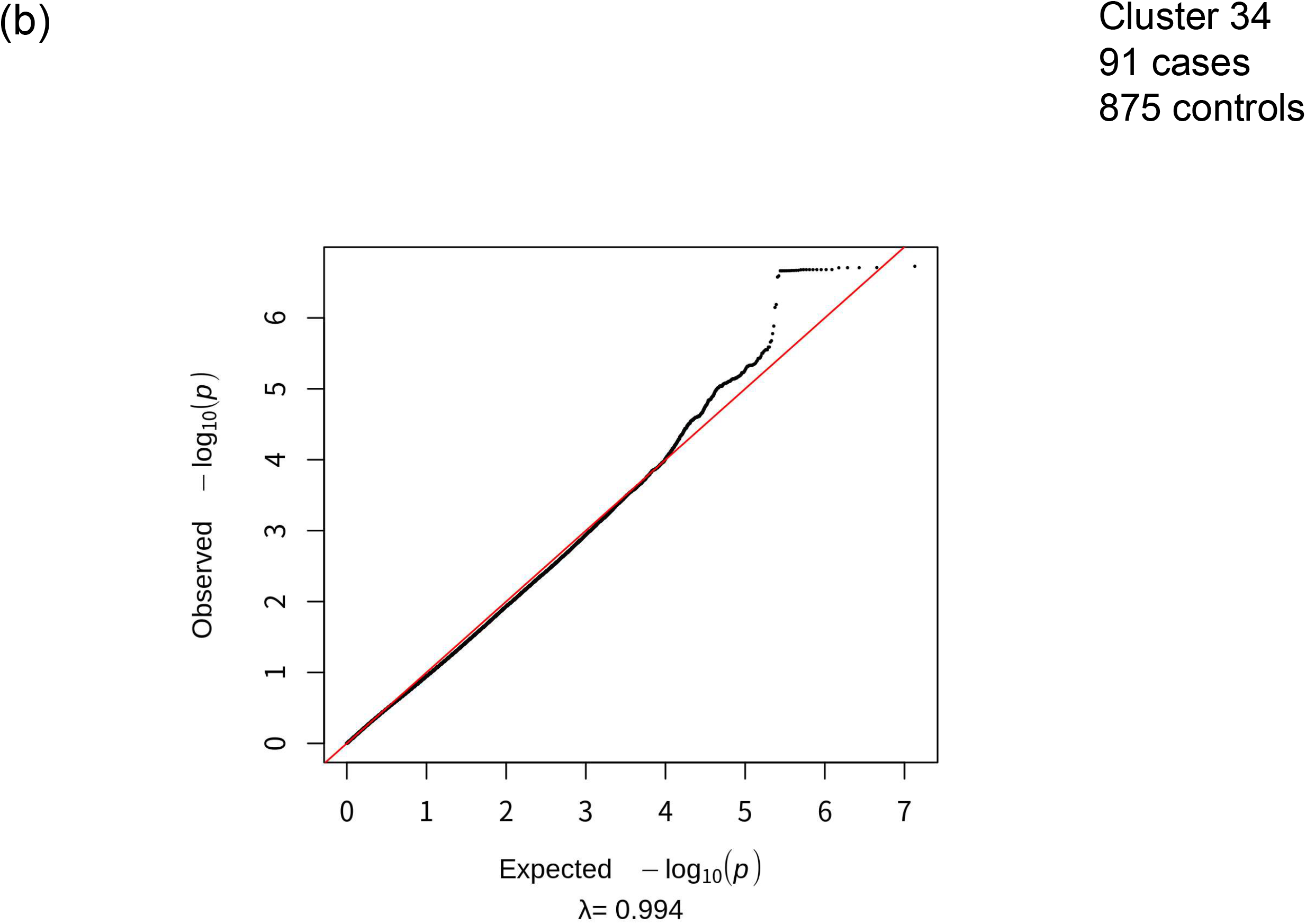
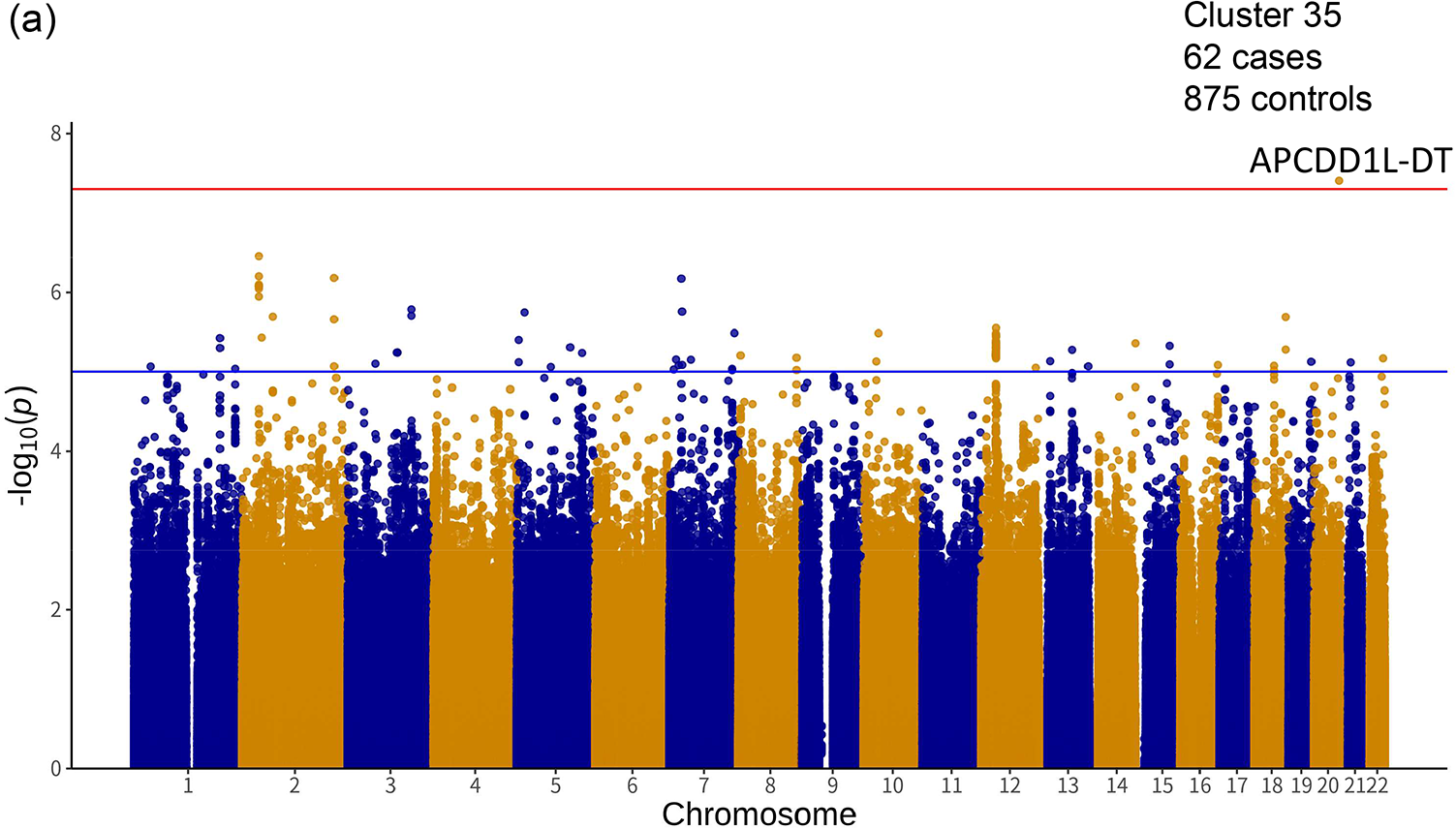
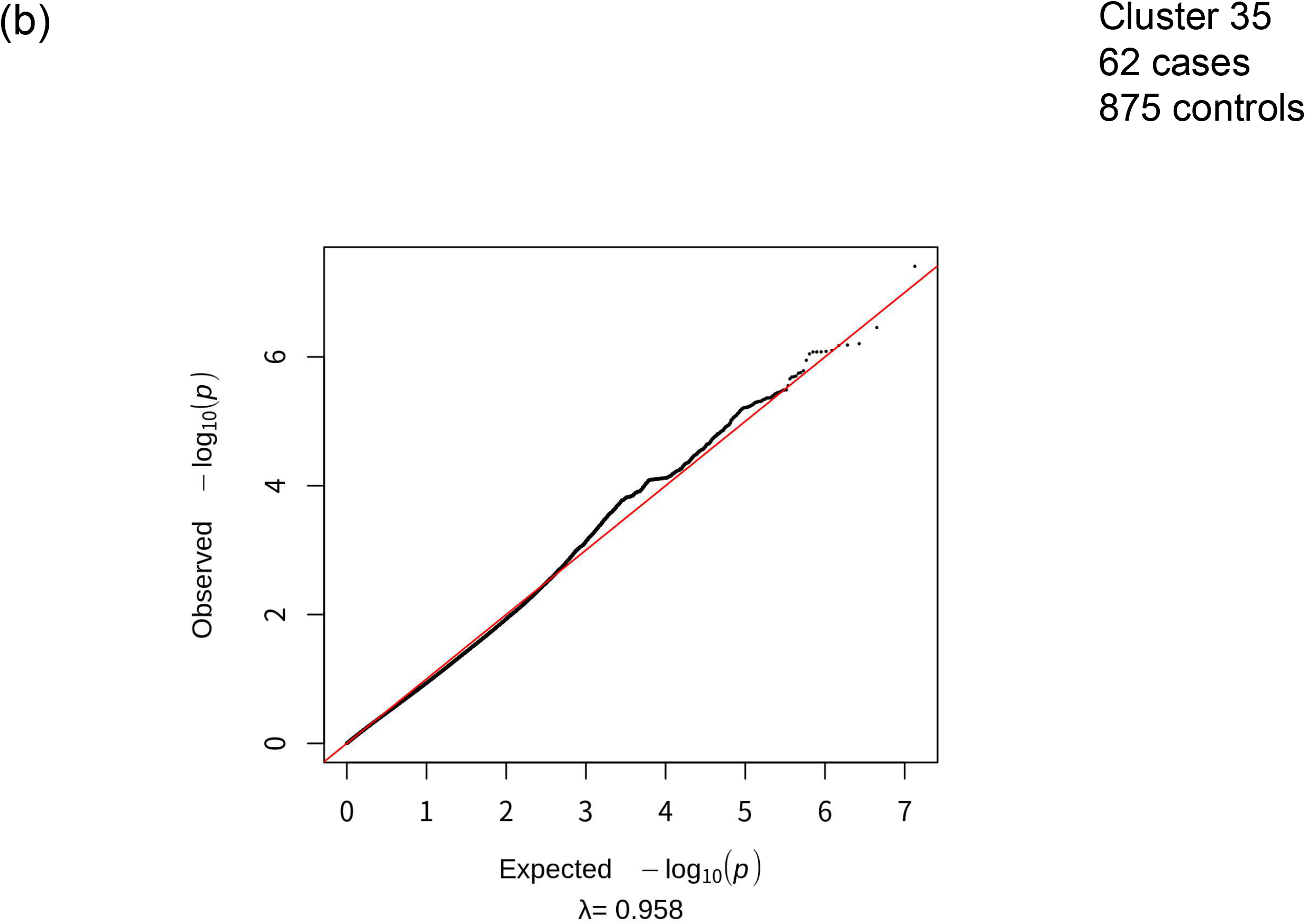
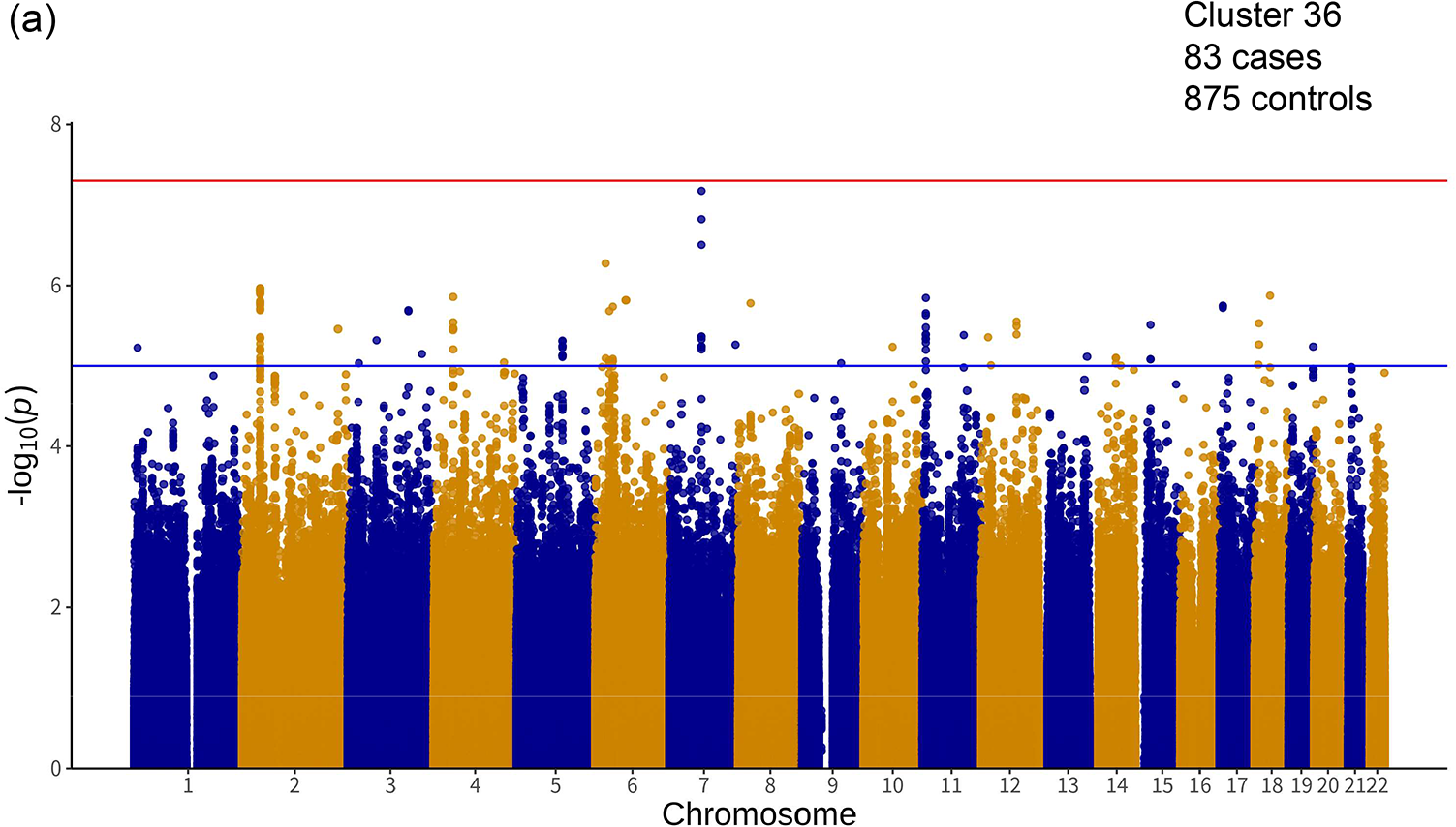
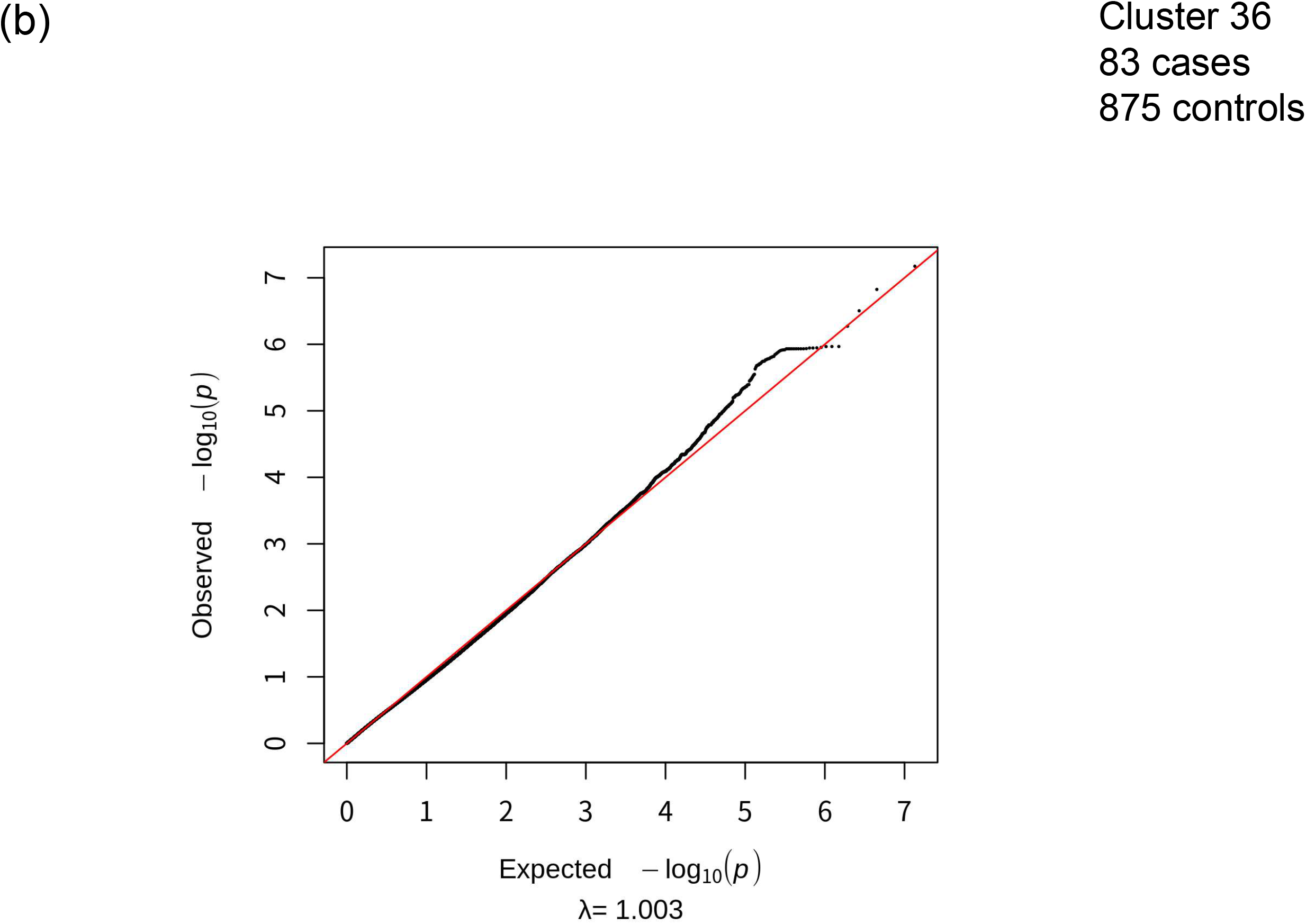
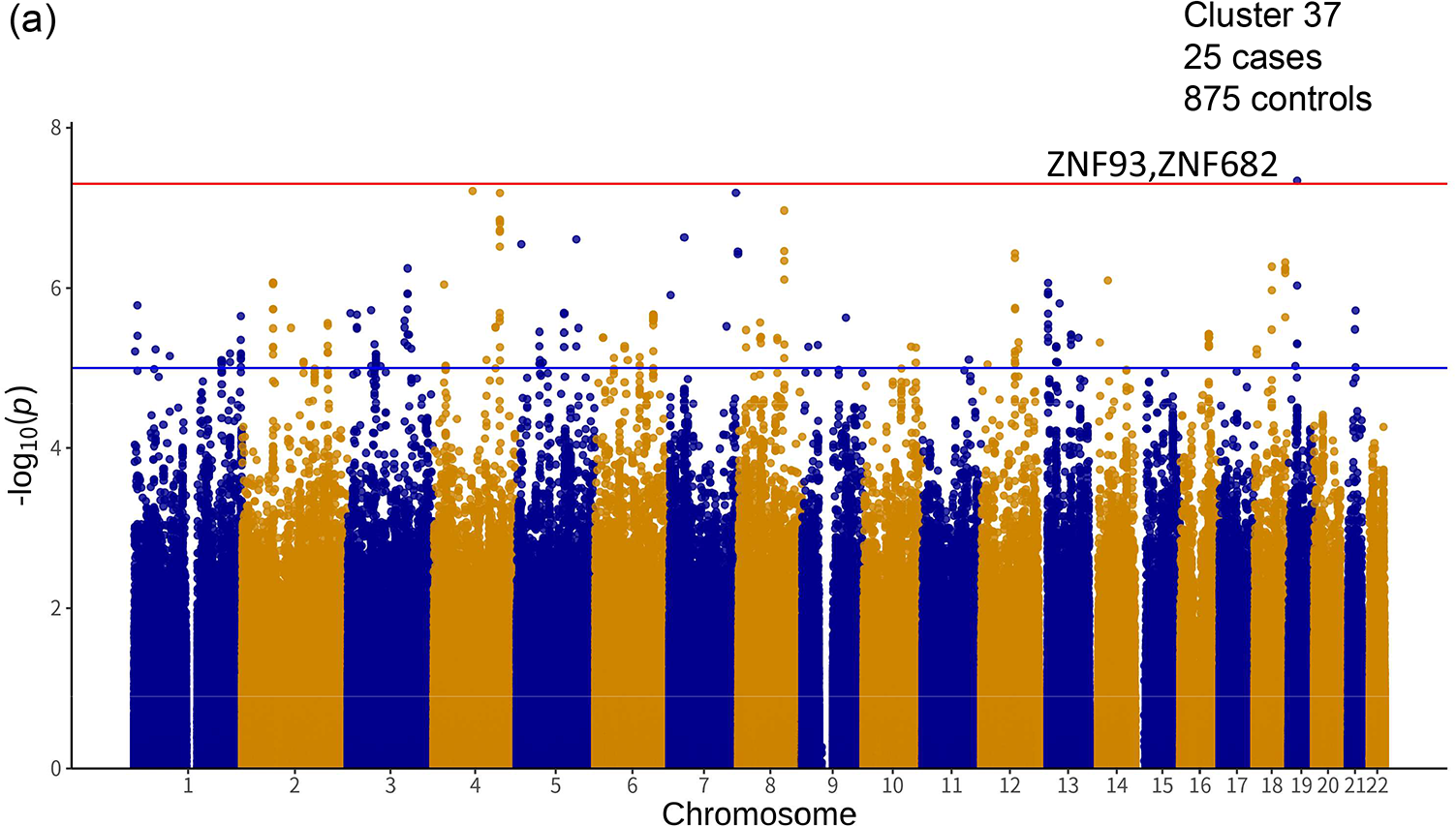
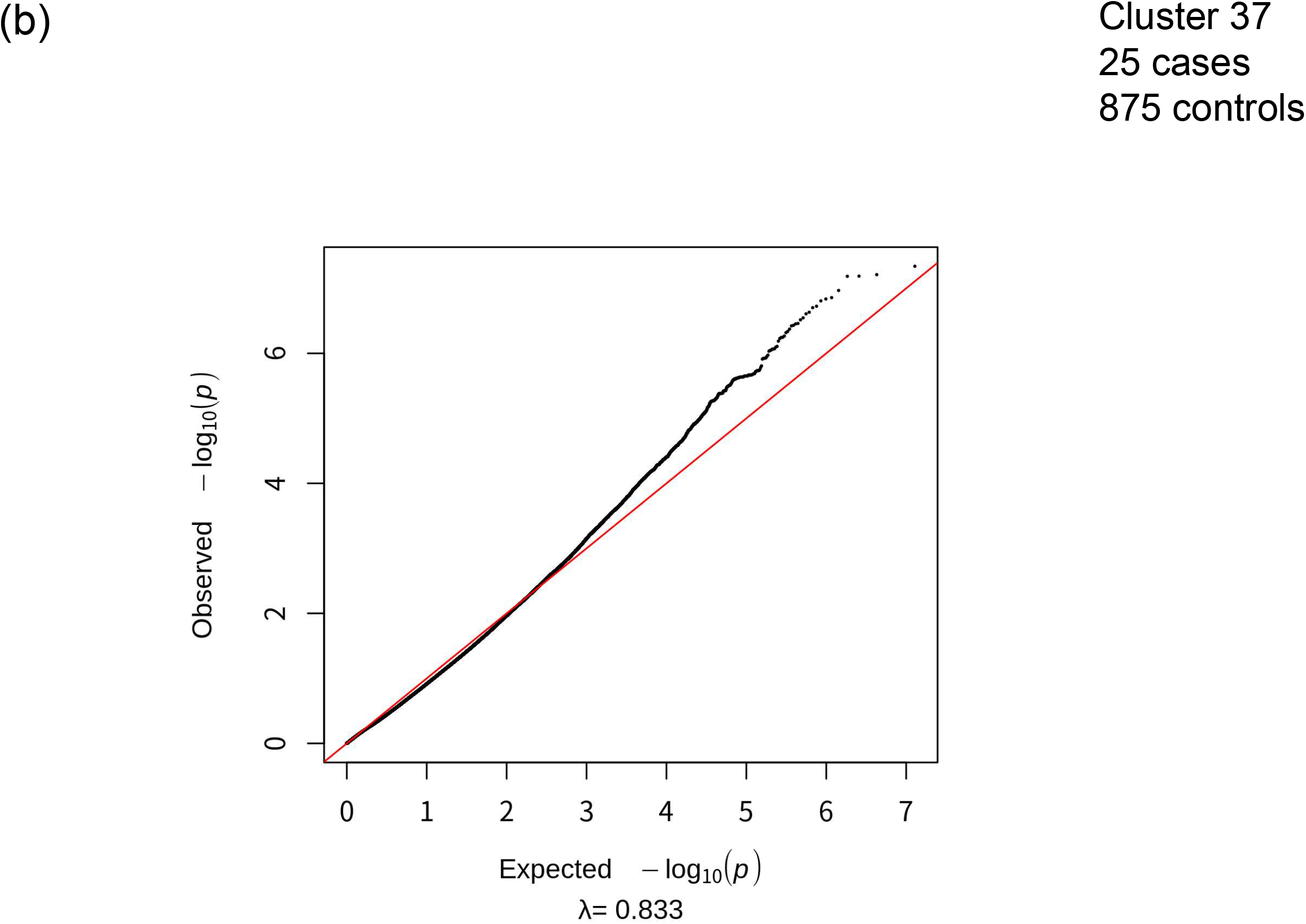
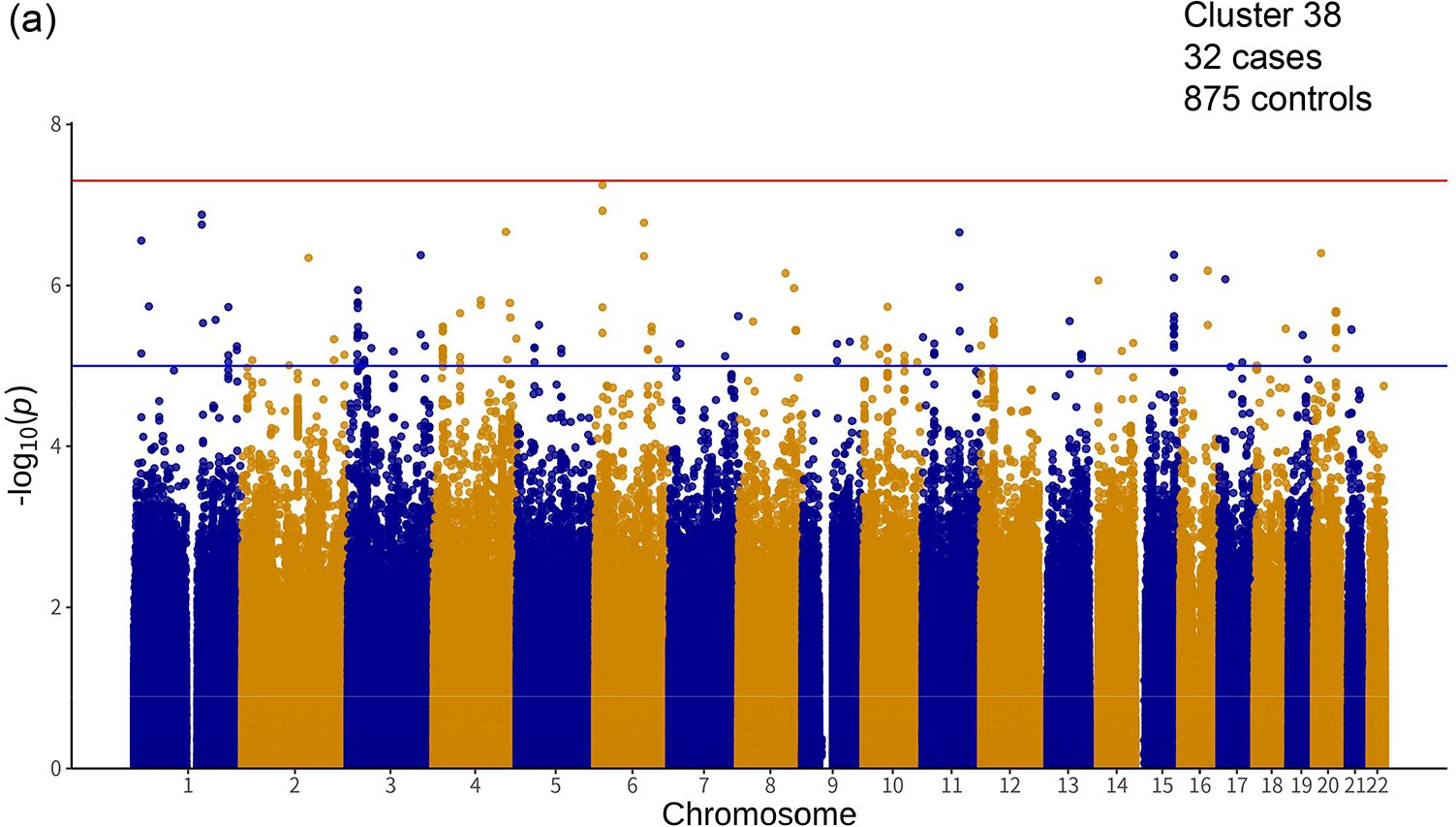
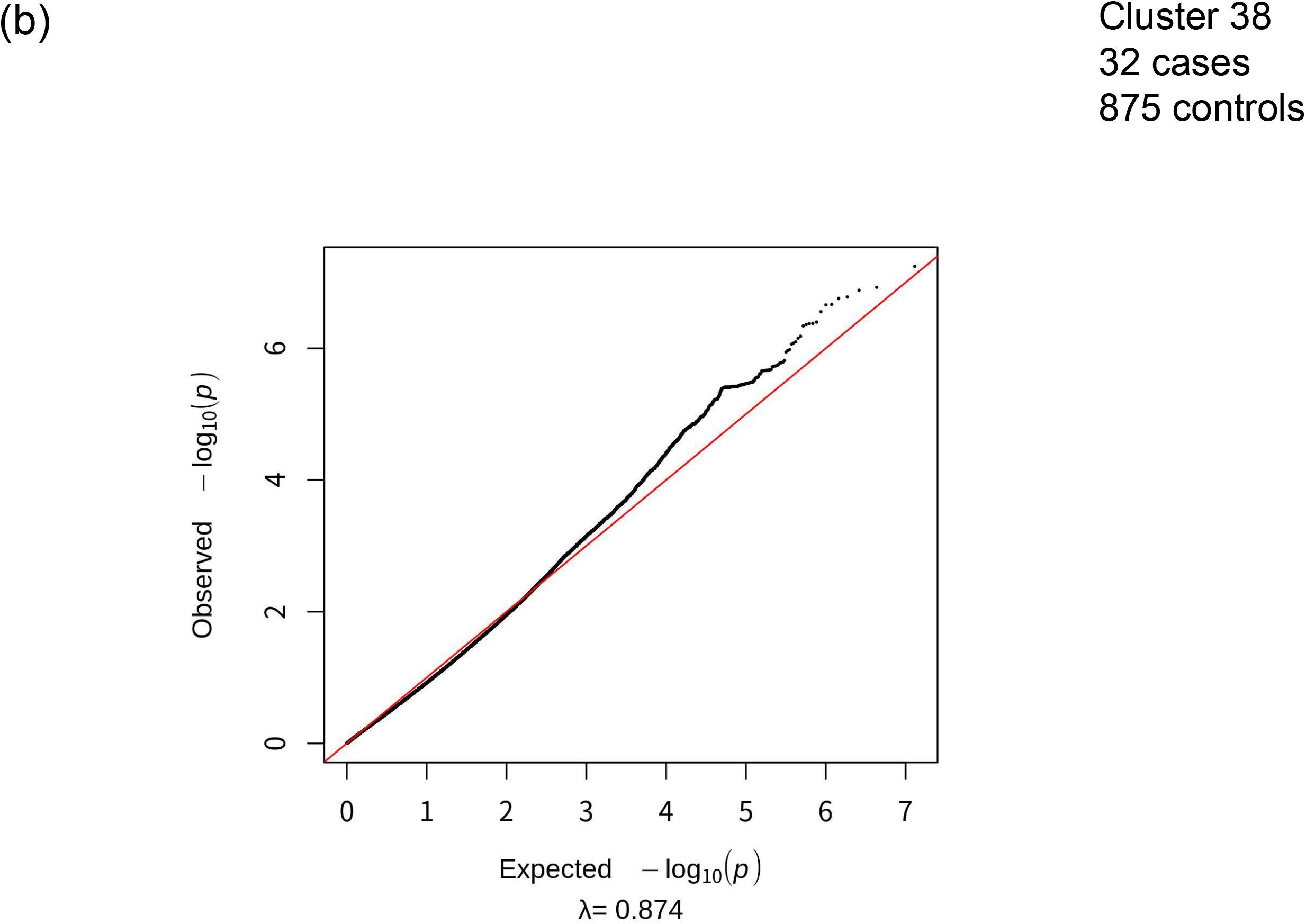
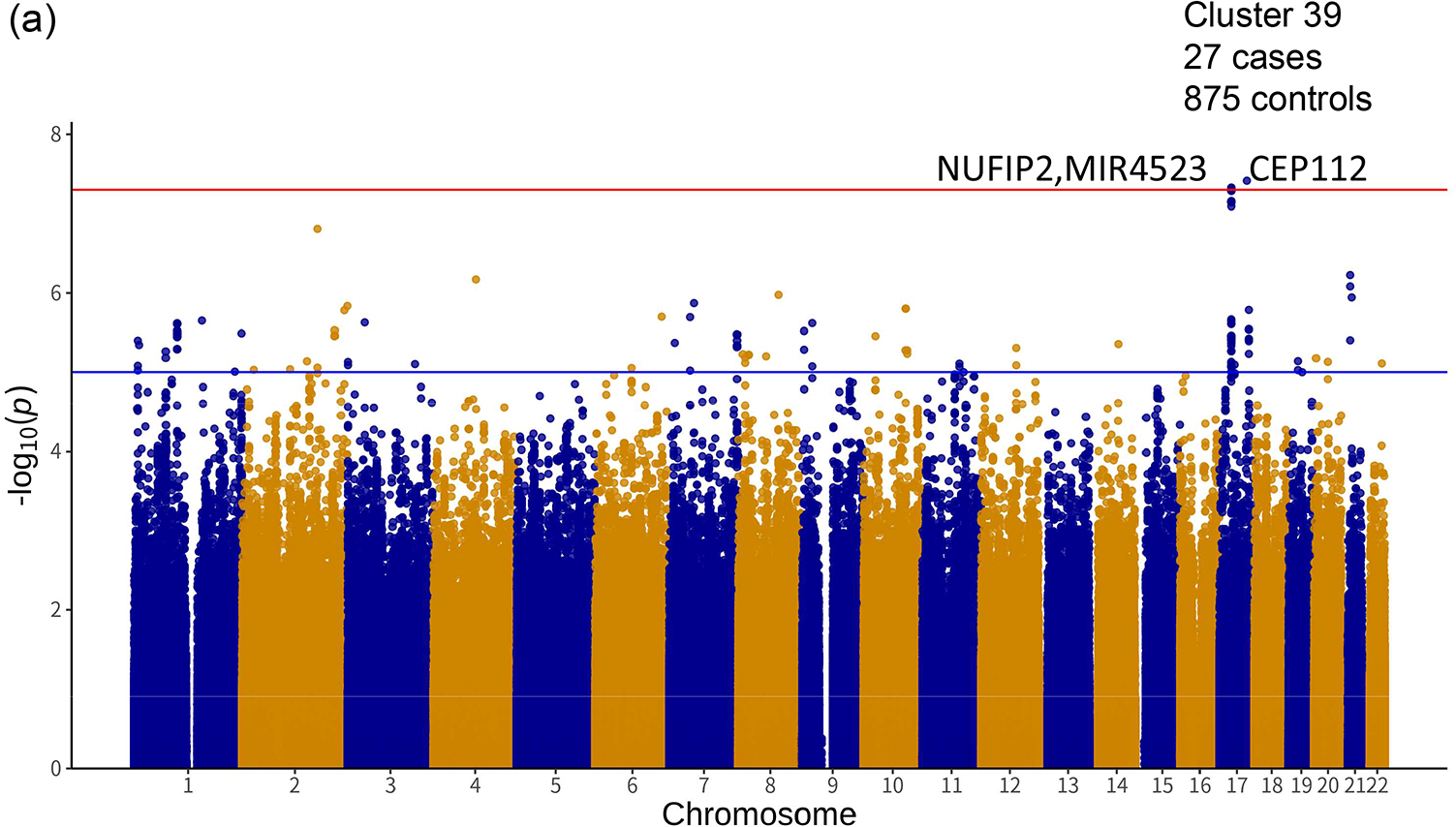
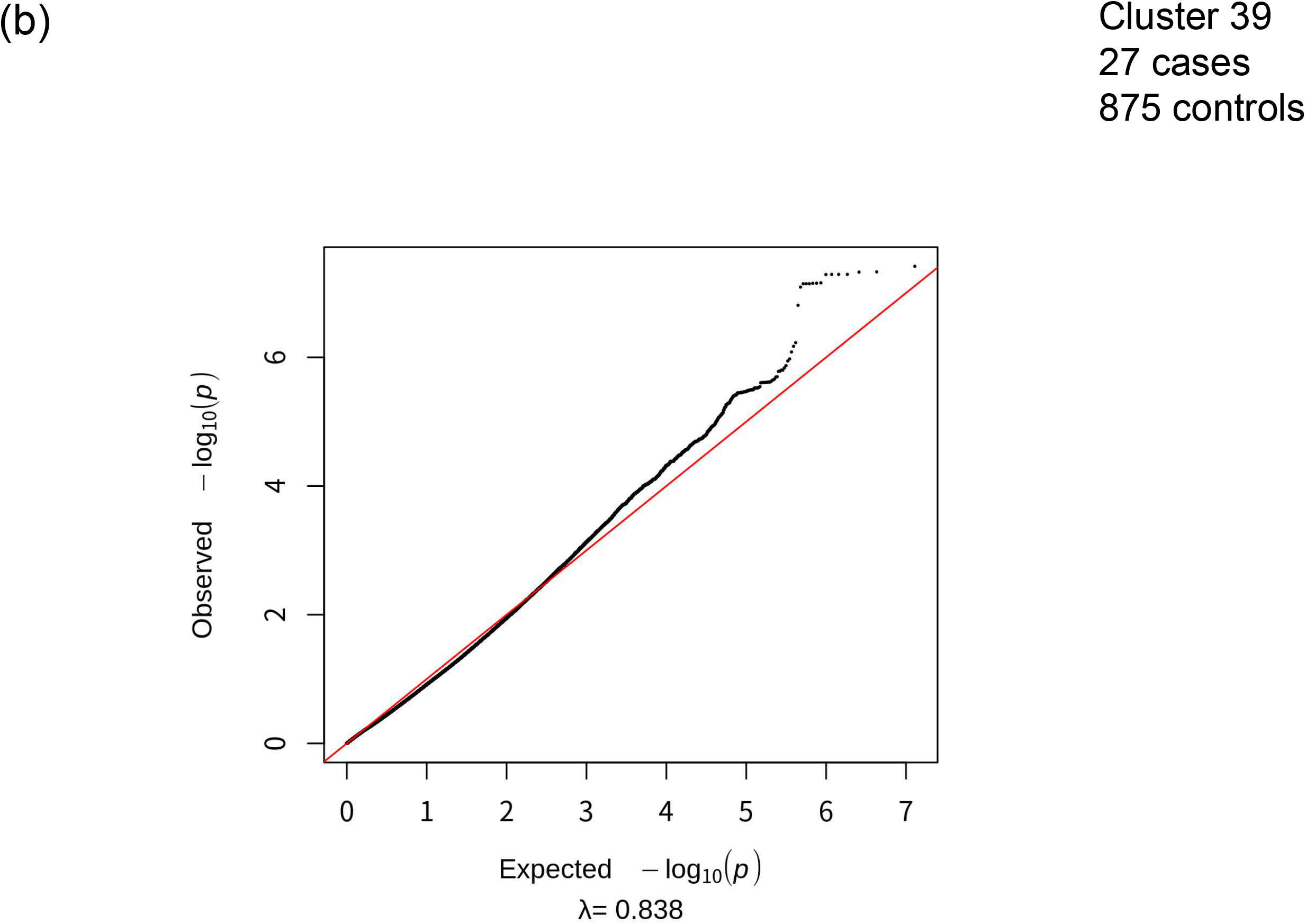
Manhattan plots (a) and corresponding quantile-quantile plots (b) for cluster-based GWASs with a cluster number of 39. We performed cluster analysis using deep embedded clustering (DEC) algorithm with a cluster number of 39 and conducted cluster-based GWAS. Among the 39 clusters, significant associations were observed in 17. In total, we observed 90 chromosomal loci, labeled in the figure, that satisfied the threshold of *P* < 5.0 × 10^−8^. The red lines indicate the threshold for genome-wide significance (*P* < 5.0 × 10^−8^). The result with more than 9 cases in each analysis is shown.

**Table 2.**
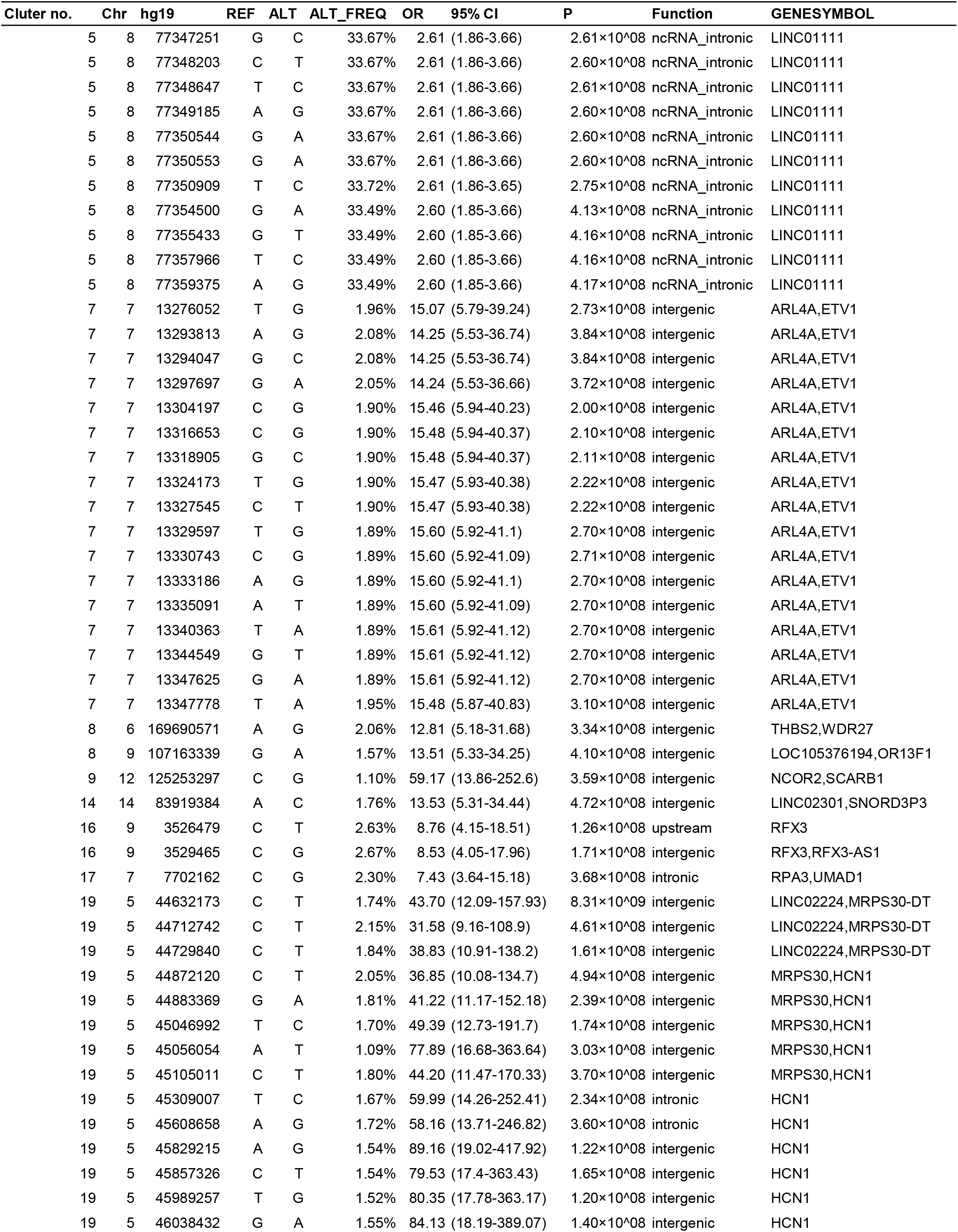
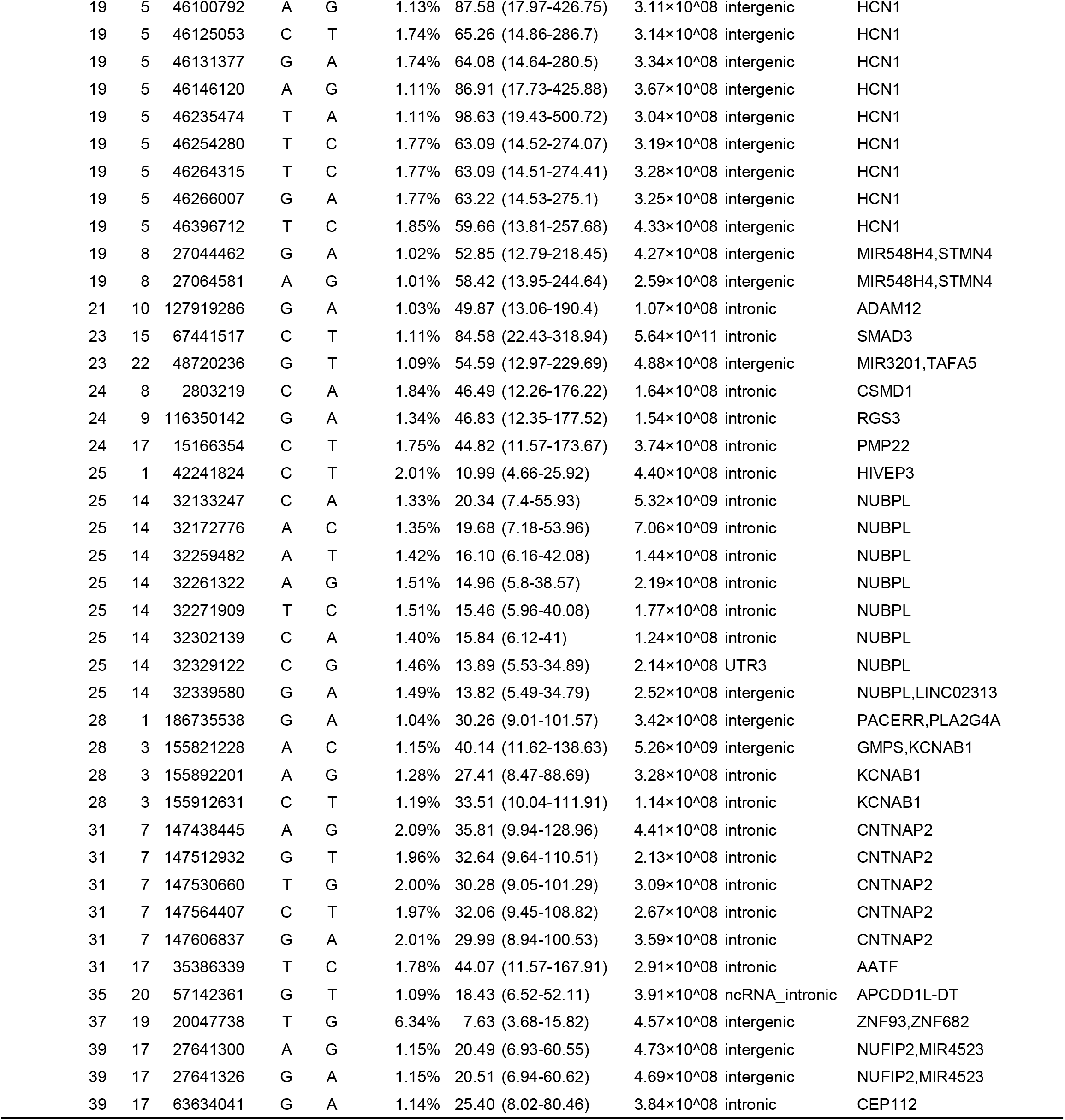
Association table of the cluster-based GWAS.

The SFARI Gene scoring system ranges from “Category 1,” which indicates “High Confidence,” to “Category 3,” which denotes “Suggestive Evidence.” Genes associated with syndromic disorders related to ASD are categorized in a different category as “#S” (e.g., 2S, 3S). Rare single gene variants, disruptions/mutations, and submicroscopic deletions/duplications related to ASD are categorized as “Rare Single Gene Mutation.”

In addition to genes in the Human Gene module of the SFARI Gene, several other important genes associated with ASD or other related disorders from previous reports were included in our findings, listed as follows (Table 3): *PLA2G4A* in Cluster 28, *STMN4* in Cluster 19, *PMP22* in Cluster 24, and *ADAM12* in Cluster 21 previously associated with ASD (*21–30*); *NCOR2* in Cluster 9, *RFX3* in Cluster16, and *CEP112* in Cluster 39 previously associated with attention deficit hyperactivity disorder (ADHD) (*28, 31, 32*); *UMAD1* in Cluster 17, *HCN1* in Cluster 19, *PACERR*, and *KCNAB1*in Cluster 28 previously associated with epilepsy (*33–37*); *KCNAB1* in Cluster 28 previously associated with mental retardation (*33*); *ADAM12* in Cluster 21 previously associated with Down syndrome (*29*); *PLA2G4A* in Cluster 28, *CSMD1* in Cluster 24, and *ADAM12* in Cluster 21, previously associated with schizophrenia (*29, 38–40*); *NCOR2* in Cluster 9 and *TAFA5* in Cluster 23 previously associated with depressive disorder (*41, 42*); *HCN1* and *MRPS30* in Cluster 19, *SMAD3* in Cluster 23, *CSMD1* in Cluster 24, and *HIVEP3* and *NUBPL* in Cluster 25 previously associated with Parkinson’s disease (*43–49*); *GMPS* in Cluster 28, *THBS2*, and *WDR27* in Cluster 8, *CSMD1* in Cluster 24, *ADAM12* in Cluster 21, *SCARB1* in Cluster 9, *AATF* in Cluster 31, *TAFA5* in Cluster 23, and *HCN1* in Cluster 19 previously associated with Alzheimer’s disease (*29, 43, 50–55*); *RPA3* in Cluster 17 previously associated with Machado-Joseph disease (*56*); and *NUFIP2* in Cluster 39 previously associated with microcephaly (*57*); *NCOR2* in Cluster 9 previously associated with spinal muscular atrophy (*58*); *PMP22* in Cluster 24 previously associated with neuropathy (*59*).

**Table 3.**
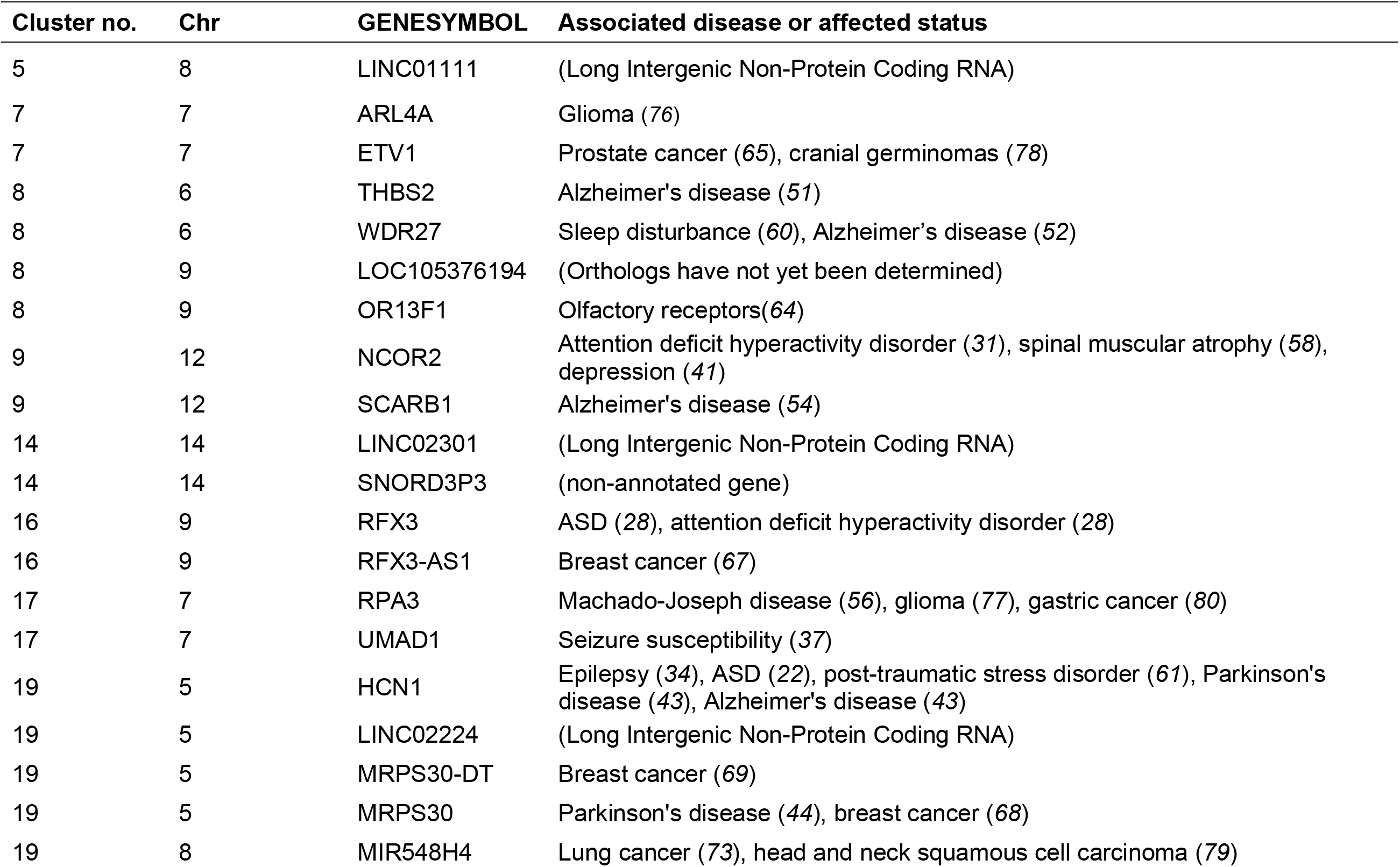
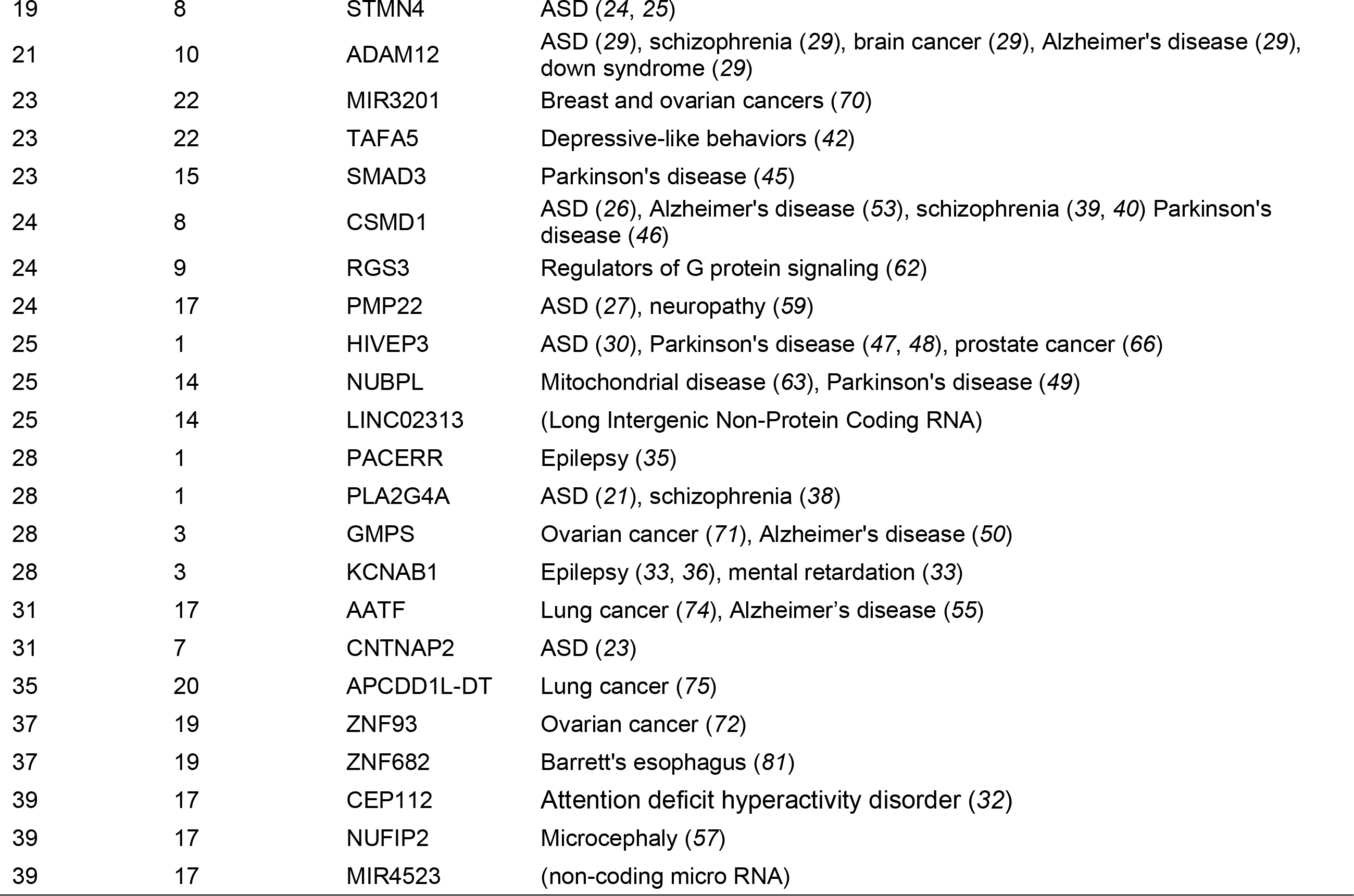
Annotation of the Genome-wide significant genes in the present cluster-based GWAS mainly in relation to autism spectrum disorder.

In addition, several important genes associated with ASD symptoms from previous reports were included in our findings as follows (Table 3): *WDR27* in Cluster 8 previously associated with sleep disturbance (*60*), and *HCN1* in Cluster 19 previously associated with post-traumatic stress disorder (*61*).

Furthermore, we observed signals in important genes associated with ASD pathways (Table 3): *RGS3* in Cluster 24, which encodes a regulator of G protein signaling (*62*); *NUBPL* in Cluster 25 is previously associated with mitochondrial disease (*63*) and *OR13F1* in Cluster 8, which encodes an olfactory receptor (*64*).

We further observed signals in various genes known to be mutated in cancer (Table 3): *ETV1* in Cluster 7 and *HIVEP3* in Cluster 25 previously associated with prostate cancer (*65, 66*); *RFX3-AS1* in Cluster 16*, MRPS30-DT,* and *MRPS30* in Cluster 19, and *MIR3201* in Cluster 23 previously associated with breast cancer (*67–70*); *GMPS* in Cluster 28 and *ZNF93* in Cluster 37 previously associated with ovarian cancer (*71, 72*) ; *MIR548H4* in Cluster 19, *AATF* in Cluster 35 and *APCDD1L-DT* in Cluster 35 previously associated with lung cancer (*73–75*); *ARL4A* in Cluster 7 and *RPA3* in Cluster 17 previously associated with glioma (*76, 77*); *ETV1* in Cluster 7 previously associated with cranial germinomas (*78*); *MIR548H4* in Cluster 19 previously associated with head and neck squamous cell carcinoma (*79*); *RPA3* in Cluster 17 previously associated with gastric cancer (*80*); and *ZNF682* in Cluster 37 previously associated with Barrett’s esophagus (*81*).

### Replication study

We previously conducted a cGWAS using a dataset from the Simons Simplex Collection (SSC) (*11*). We considered the agreement between the SSC and SPARK results as evidence of successful replication. Having analyzed the SSC data, we found statistically significant SNPs in the cGWAS in the following genes: *CDH5, CNTN5, CNTNAP5, DNAH17, DPP10, DSCAM, FOXK1, GABBR2, GRIN2A5, ITPR1, NTM, SDK1, SNCA, SRRM4*, and *ZNF678*. Proteins encoded by *CNTNAP5* and *ZNF678* had CNTNAP and ZNF domains, respectively. In the present study, proteins encoded by *CNTNAP2* in Cluster 30 and *ZNF93* and *ZNF682* in Cluster36 also had CNTNAP and ZNF domains, respectively.

## Discussion

One of the most important contributions of the present study is that we reasonably reduced the sample size while increasing the statistical power based on multiple ASD rating scales using a larger dataset than our previous study (*11*) and a new clustering algorithm. At least two factors likely reduced the possibility of false-positive detections of statistically significant SNPs in the present cGWAS. First, we validated the usefulness and feasibility of the concept proposed in a previous simulation study (*10*), which indicated that homogeneous case subgroups increase the power in the genetic association studies using real-world data. Second, a substantial number of statistically significant SNPs in cGWAS observed in the present study (Table 2, Figure 2) were located within or near previously reported candidate genes for ASD (Table 3). If all statistically significant SNPs in the present study were false positives, then there would be no reasonable explanation for the several concentrated ASD-related genes. Because of the abovementioned reasons, it may be difficult to explain the present results only in terms of false positives.

In addition to genes directly associated with ASD suggested by previous studies, we observed several other genes associated with ASD-related diseases or symptoms (*28, 29, 31–36, 38–57, 60–64*). The ASD phenotype overlaps with other conditions such as ADHD, epilepsy, mental retardation, down syndrome, schizophrenia, depressive symptoms, Parkinson’s disease, Alzheimer’s disease, Machado-Joseph’s disease, and post-traumatic stress disorders. Therefore, genes associated with these diseases were also detected in this study. For example, signals in some of these genes related to Parkinson’s disease might be interpreted as follows: The presence of a certain gene mutation may be seen as an ASD-like symptom in childhood and diagnosed as ASD, but with aging and cumulative exposure to environmental factors, it may change slightly to a Parkinson’s disease-like symptom and be diagnosed as Parkinson’s disease. Alternatively, they might not have been diagnosed with ASD during childhood yet diagnosed with Parkinson’s disease in old age. Furthermore, sleep disturbance and microcephaly are relatively frequently observed in patients with ASD. Dysregulation of G-protein signaling or mitochondrial dysfunctions also have been reported as ASD etiologies (*62, 63*). Almost all statistically significant genes in the present study revealed by the cGWAS were associated with ASD, ASD symptoms, and/or ASD pathways. These findings suggest that clustering might successfully identify subgroups with relatively homogeneous disease etiologies.

Several statistically significant SNPs in the present study have been reported to be associated with cancer. Although, at first glance, such a result may be considered as a false positive, recent genome-wide exome sequencing has revealed extensive overlap of risk genes for ASD and cancer (*65–81*). For example, *ETV1* is associated with prostate cancer; *RFX3-AS1, MRPS30, MRPS30-DT,* and *MIR3201* are associated with breast cancer; *MIR548H4, AATF,* and *APCDD1L-DT* are associated with lung cancer. These cancers are highly associated with ASD (*82, 83*). Considering genes involved in both ASD and cancer, the identification of a cancer-associated gene in some clusters is almost identical to that of ASD-associated genes. Therefore, it would be difficult to explain these findings solely in terms of false positives.

To the best of our knowledge, associations between ASD and variants in loci such as *LINC01111*, *LINC02224*, *LINC02301*, *LINC02313*, and *LOC105376194* have not been previously reported. These results might suggest that novel genetic loci of ASD can be found by identifying better-defined subgroups.

The heat map illustrates the unique characteristics of each cluster obtained in the present study (Table 1). For example, cluster 8 was associated with relatively lower scores on DCDQ than other clusters but higher scores on SCQ and RBS-R, with genes such as *THBS2*, *WDR27*, and *ORF13F1* being associated with Alzheimer’s disease. The characteristics of this cluster, in which repetitive behavior and social communication deficits were more prevalent whereas motor skills were relatively preserved, are consistent with some features of Alzheimer’s disease inferred from functions of these genes. Meanwhile, cluster 25 had relatively lower scores on RBS-R than other clusters but higher scores on DCDQ and SCQ. In addition, *HIVEP3* and *NUBPL* are primarily associated with Parkinson’s disease. The characteristics of this cluster are consistent with Parkinson’s disease features. In cluster 7, no major characterization was observed in DCDQ, SCQ, and RBS-R whereas *ARL4A* and *ETV* are cancer-associated genes. The weakness of ASD characteristics in this cluster may be attributed to its association with cancer. Similar relationships were observed in other clusters. These results suggest the potential association between cluster characteristics and gene function; thus, the clusters obtained were in part functionally valid. Nevertheless, it is important to note that the characteristics of the clusters may not necessarily be human-recognizable. Because artificial intelligence extracts feature that lie in the combination of many variables, the resulting clusters, although more homogeneous, may not always be easy for humans to understand. In other words, artificial intelligence may discover clusters that humans have not been able to discover to date. It will be necessary in the future to define the clusters that artificial intelligence has discovered.

The purpose of dividing a disease into subgroups is to provide optimal treatment to cure or mitigate the disease in that group. There are at least two ways to divide a disease into subgroups. One is to identify subgroups that can predict disease severity. This approach can be used to create subgroups by combining existing severity factors and other factors. However, while this approach may predict disease severity, but does not necessarily prevent or reduce the severity of the disease, since the subgroups may have different etiology. The severity factor may also be heterogeneous. Conversely, the other approach is to divide the patients into subgroups based on genetic architecture. In this approach, subgroups with different etiologies can be identified, and the genetic architecture of the subgroups can be clarified to determine targets for therapy. Therefore, when dividing a disease into subgroups, it is necessary to identify the subgroups so that the genetic architecture, as the etiology, becomes clear to identify etiologically different subgroups and elucidate each genetic architecture for precision therapy.

The present study has some limitations. First, the replication was not sufficient between the findings of our previous study (*11*) and those of the present study. No significant signal was observed in conventional GWAS, but significant signals were observed in GWAS after patients were divided into clusters. Also, the significant signals of the two studies were partially consistent. In both the present and previous studies, signals in genes encoding proteins with CNTNAP and ZNF domains were consistently observed. These domains are associated with a contact in the associated protein family and zinc finger protein superfamily, respectively hence the replication was partially obtained. Nevertheless, this does not suggest that reproducibility has been achieved. However, since the SPARK cohort is one of the largest genetic cohorts dealing with ASD, it is difficult to obtain replication datasets of similar sample sizes for sufficient statistical power. Furthermore, if nine genes in one study are identified out of 1,000 genes, the probability of conducting the same study again without selecting those genes is (_1000_C_9*991_C_9_) / (_1000_C_9*1000_C_9_) = 0.92. Also, the differences in variables in our previous studies (*11*) and those of the present study should be taken into account. Our previous study used ADI-R (*84*) scores and history of vitamin treatment (*85, 86*), whereas the present study used DCDQ, SCQ, and RBS-R scores. In addition, machine learning does not always produce consistent results. In future studies, algorithms that yield the same clusters, even when repeated, may also be considered. The cluster number was fixed at 39 in the present study in reference to the λ value. If ASD arises as a result of >1,000 etiology terms, in the future, it will be necessary to confirm the reproducibility with larger data sets that yield more subtypes.

Second, it remains unclear whether the selection of variables, algorithm, and cluster number in the present study was optimal. We selected phenotypic variables of DCDQ, SCQ, and RBS-R scores, age of individual at initial registration in months, ethnicity, dominant hand, history of medication, history of biomedical intervention, and history of intensive behavioral intervention. ASD has several other symptoms and characteristics (*2, 85, 86*). In future studies, as many variables as possible should be considered and narrow them down appropriately, considering their relationship to each other. Regarding the algorithms, we selected DEC algorithm (*15, 16*), which simultaneously learns feature representations and cluster assignments using deep neural networks, one of the most up-to-date techniques. However, although DEC was useful, different types of algorithms will likely emerge in the future. Such algorithms should be considered in due course on a case-by-case basis.

Third, the present study could not clarify whether each cluster is polygenic, oligogenic, or both. For example, six genome-wide significant genes were detected in cluster 19. It is unclear whether this cluster is composed of six further subtypes or whether these six genes are required simultaneously. By clarifying whether a genetic etiology of a cluster is polygenic, oligogenic, or both and characterizing individual clusters, we could elucidate the genetic architecture of ASD.

Although further research is warranted to fully characterize ASD, our study provides novel clues toward facilitating the development of precision medicine for this disorder. For clusters that can be explained by the polygenic model, it is not always easy to explore the pathogenesis more deeply in animal experiments, and there may be many difficulties in identifying target molecules and linking them to drug discovery. However, if the clusters can be explained by an oligogenic model, it may be relatively easy to conduct animal experiments and identify target molecules, so molecular exploration of the clusters that we identified in the present study could be proceeded. Once the clusters are identified, a classifier will be created using the cluster numbers as teacher data, which will be used to classify patients broadly diagnosed with ASD into subgroups based on their symptoms and other factors. It may be possible to examine the effectiveness of treatments for each of the clusters, in this way, a part of precision medicine for ASD might be realized. Although some subtypes of ASD may still be explained by the polygenic models, the present study demonstrated that ASD is an aggregation of etiologically heterogeneous subtypes, each of which has its own genetic architecture involving a smaller number of genetic loci than previously thought.

## Acknowledgments

The authors would like to thank the families at the participating Simons Foundation Powering Autism Research for Knowledge (SPARK), as well as the staff at the Simons Foundation Autism Research Initiative (SFARI). We are also grateful to Shoji Tanaka for assistance with the present study.

## Funding

The present study was supported by the Ministry of Education, Culture, Sports, Science and Technology (MEXT) KAKENHI Grant Numbers 19H03894, 21K17304 and 22H03346. MEXT had no role in the design or execution of the study

## Author contributions

Conceptualization: Fumihiko Ueno, Tomomi Onuma, Ippei Takahashi, Hisashi Ohseto, Shinichi Kuriyama

Methodology: Fumihiko Ueno, Tomomi Onuma, Ippei Takahashi, Hisashi Ohseto, Akira Narita, Taku Obara, Mami Ishikuro, Keiko Murakami, Aoi Noda, Fumiko Matsuzaki, Hirohito Metoki, Gen Tamiya, Shigeo Kure, Shinichi Kuriyama

Visualization: Fumihiko Ueno

Funding acquisition: Fumihiko Ueno, Shinichi Kuriyama

Project administration: Fumihiko Ueno, Taku Obara, Ippei Takahashi

Supervision: Shinichi Kuriyama

Writing – original draft: Fumihiko Ueno, Shinichi Kuriyama

Writing – review & editing: Fumihiko Ueno, Tomomi Onuma, Ippei Takahashi, Hisashi Ohseto, Akira Narita, Taku Obara, Mami Ishikuro, Keiko Murakami, Aoi Noda, Fumiko Matsuzaki, Hirohito Metoki, Gen Tamiya, Shigeo Kure, Shinichi Kuriyama

## Competing interests

Authors declare that they have no competing interests.

## Data and materials availability

All data used in this study are available only to those who are granted access by the Simons Foundation.

## Supplementary Materials

Supplementary Figure S1

Supplementary Figure S2

Supplementary Figure S3

## Methods

### Participants

We performed the present study in accordance with the guidelines of the Declaration of Helsinki (*87*) and all other applicable guidelines. The protocol was reviewed and approved by the Institutional Review Board of the Tohoku University Graduate School of Medicine (2020-1-826, 2020-1-991), and informed consent was obtained from the participants by SFARI for the SPARK study (*14*). In SPARK, phenotypic data and biospecimens were collected remotely, so that participants could complete the study protocol online at their convenience. Individuals living in the United States with a professional diagnosis of ASD made by a physician, psychologist, or therapist, as well as their parents and unaffected siblings were eligible to participate in SPARK. Phenotype information and ASD diagnoses in SPARK are self-or parent-reported. The Interactive Autism Network suggests that parent-reported diagnosis of ASD is highly valid (*88*).

### Datasets

We used phenotypic variables, background history, and genotypic data from the SPARK database, which was publicly released in October 2017 and is directly available from SFARI (*14*). From the SPARK WES1 (27K) dataset, we used data from 2,685 affected white male probands for whom data from all three tests of the DCDQ (*18*), RBS-R (*19*), and SCQ (*20*) were available and 891 unaffected brothers of probands for whom genotype data generated by the Illumina Infinium Global Screening Array (GSA) v1.0 were available. From the 2,685 affected probands, we excluded 606 who were biologically related to the unaffected brothers. Therefore, 2,079 probands and 891 unaffected brothers were available for subsequent analyses. We excluded participants whose ancestries were estimated to be different from other participants using principal component analyses performed using EIGENSOFT version 7.2.1 (*89*) for the genotype data. Based on the principal component analyses, we excluded 33 persons whose data points beyond six standard deviations of principal components 1 or 2 (Supplementary Figure S2). Therefore, 2,062 probands and 875 unaffected brothers were available for subsequent clustering and genetic analyses.

### Clustering

We conducted cluster analyses using phenotypic variables of DCDQ (15 items), SCQ (40 items), RBS-R (43 items), age of the individual at initial registration in months, ethnicity, dominant hand, history of medication, history of biomedical intervention (e.g., diet, alternative medicine, supplements), and history of intensive behavioral intervention (e.g., Applied Behavior Analysis, Verbal Behavior, Pivotal Response Treatment). Missing data were imputed with the mean value of each variable, and all categorical data were transformed into dummy variables.

We applied DEC (*15, 16*), which uses the algorithm of deep learning, to conduct a cluster analysis to divide the dataset, including data from individuals with ASD, into subgroups using phenotypic variables. The DEC algorithm requires a cluster number (k), iteration number, epoch number, and network dimension determined by researchers. We set an a priori k of 40 assuming that ASD consists of hundreds of subgroups (*6*) and considering statistical power following sample size calculations (*90*). Other hyperparameters for DEC were batch size of 256,300 pre-training epochs, 400 max iterations, 30 update intervals, and 0.001 tolerance for the stopping criterion. We performed the analyses using the scikit-learn toolkit in Python 2.7.

Clustering is an exploratory data analysis technique, and the validity of clustering results may be judged by external knowledge, such as the purpose of the segmentation (*91*). Several methods have been proposed to prespecify a cluster number of k, such as visual examination of the data and likelihood and error-based approaches; however, these methods do not necessarily provide mutually consistent results (*13*). Although there are measures for evaluating the quality of the clusters (*92*), the number of clusters should also be determined according to the research purposes. Therefore, we calculated the inflation factor (λ) of quantile-quantile (Q-Q) plots using the logarithm of the *P*-value to base 10 (−log_10_P) for each cluster number. Under the hypothesis that ASD consists of hundreds of subgroups (*6*), we compared λ values giving larger numbers of clusters as priority. We then set hyperparameter k of DEC.

### Genotype data and quality control

We used the SPARK WES1(27K) genotypic dataset, in which probands and unaffected brothers had already been genotyped previously (https://gpf.sfari.org/hg38/datasets/SFARI_SPARK_WES_1_CONSORTIUM/dataset-description). We used the dataset genotyped by the Illumina GSA v1.0 array, which has 642,824 probes. We excluded SNPs with a minor allele frequency < 0.01, call rate < 0.95, and Hardy-Weinberg equilibrium test *P* < 0.000001.

We imputed SPARK WES1(27K) genotypic data independently from phenotype data using the Michigan imputation server. The human genome reference build of the genotypic data was converted from hg38 to hg19 using LiftOver, a tool provided by the UCSC Genome browser (http://genome.ucsc.edu/cgi-bin/hgLiftOver). On the Michigan imputation server, we selected “HRC r1.1 2016 (GRCh37/hg19)” for Reference Panel, “0.3” for Rsq Filter, “Eagle v2.4” for Phasing, and “Other/Mixed” for population options, and performed quality control and imputation.

After genotype imputation, the SPARK WES1(27K) genotypic dataset included 33,717,335 SNPs in the autosome.

### Statistical analysis

For a preliminary study, we conducted a conventional GWAS comparing all patients to all controls in the whole SPARK WES1(27K) genotypic dataset, with 2,062 male probands and 875 unaffected brothers. The control group did not include the brothers of affected participants. Unaffected brothers were selected from other families of the present probands, including siblings of probands who did not respond to the all survey forms and siblings whose sisters were probands. In the second step, we conducted a cGWAS in each subgroup of the cases, which were divided using the DEC algorithm (*15, 16*), and controls. Details of the study design are shown in Supplementary Figure S3. We applied the logistic regression model to calculate the additive allele dosage effect, which examines the risk of disease in those with the alternative allele of interest compared to those with the reference allele.

Association analyses were performed with PLINK software package (*93*). The detected SNPs were subsequently annotated using ANNOVAR (*94*). Manhattan plots and Q-Q plots were generated using R (version 4.1.0) (*95*).

## Notes

### Competing Interest Statement

The authors have declared no competing interest.

## References

1. A. P. Association, Diagnostic and Statistical Manual of Mental Disorders, 5th *Edition:* DSM-5 (American Psychiatric Publishing, Washington, D.C, 5th edition., 2013).

2. C. Lord, M. Elsabbagh, G. Baird, J. Veenstra-Vanderweele, Autism spectrum disorder. Lancet. 392, 508–520 (2018).

3. D. H. Geschwind, M. W. State, Gene hunting in autism spectrum disorder: on the path to precision medicine. Lancet Neurol. 14, 1109–1120 (2015).

4. A. Bailey, A. Le Couteur, I. Gottesman, P. Bolton, E. Simonoff, E. Yuzda, M. Rutter, Autism as a strongly genetic disorder: evidence from a British twin study. Psychol Med. 25, 63–77 (1995).

5. M. B. Lauritsen, C. B. Pedersen, P. B. Mortensen, Effects of familial risk factors and place of birth on the risk of autism: a nationwide register-based study. J Child Psychol Psychiatry. 46, 963–971 (2005).

6. Gene Scoring Module. SFARI Gene, (available at https://gene.sfari.org/database/gene-scoring/).

7. J. Grove, S. Ripke, T. D. Als, M. Mattheisen, R. K. Walters, H. Won, J. Pallesen, E. Agerbo, O. A. Andreassen, R. Anney, S. Awashti, R. Belliveau, F. Bettella, J. D. Buxbaum, J. Bybjerg-Grauholm, M. Bækvad-Hansen, F. Cerrato, K. Chambert, J. H. Christensen, C. Churchhouse, K. Dellenvall, D. Demontis, S. De Rubeis, B. Devlin, S. Djurovic, A. L. Dumont, J. I. Goldstein, C. S. Hansen, M. E. Hauberg, M. V. Hollegaard, S. Hope, D. P. Howrigan, H. Huang, C. M. Hultman, L. Klei, J. Maller, J. Martin, A. R. Martin, J. L. Moran, M. Nyegaard, T. Nærland, D. S. Palmer, A. Palotie, C. B. Pedersen, M. G. Pedersen, T. dPoterba, J. B. Poulsen, B. S. Pourcain, P. Qvist, K. Rehnström, A. Reichenberg, J. Reichert, E. B. Robinson, K. Roeder, P. Roussos, E. Saemundsen, S. Sandin, F. K. Satterstrom, G. Davey Smith, H. Stefansson, S. Steinberg, C. R. Stevens, P. F. Sullivan, P. Turley, G. B. Walters, X. Xu, Autism Spectrum Disorder Working Group of the Psychiatric Genomics Consortium, BUPGEN, Major Depressive Disorder Working Group of the Psychiatric Genomics Consortium, 23andMe Research Team, K. Stefansson, D. H. Geschwind, M. Nordentoft, D. M. Hougaard, T. Werge, O. Mors, P. B. Mortensen, B. M. Neale, M. J. Daly, A. D. Børglum, Identification of common genetic risk variants for autism spectrum disorder. Nat Genet. 51, 431–444 (2019).

8. P. M. Visscher, M. E. Goddard, From R.A. Fisher’s 1918 Paper to GWAS a Century Later. Genetics. 211, 1125–1130 (2019).

9. T. Gaugler, L. Klei, S. J. Sanders, C. A. Bodea, A. P. Goldberg, A. B. Lee, M. Mahajan, D. Manaa, Y. Pawitan, J. Reichert, S. Ripke, S. Sandin, P. Sklar, O. Svantesson, A. Reichenberg, C. M. Hultman, B. Devlin, K. Roeder, J. D. Buxbaum, Most genetic risk for autism resides with common variation. Nat Genet. 46, 881– 885 (2014).

10. M. Traylor, H. Markus, C. M. Lewis, Homogeneous case subgroups increase power in genetic association studies. Eur J Hum Genet. 23, 863–869 (2015).

11. A. Narita, M. Nagai, S. Mizuno, S. Ogishima, G. Tamiya, M. Ueki, R. Sakurai, S. Makino, T. Obara, M. Ishikuro, C. Yamanaka, H. Matsubara, Y. Kuniyoshi, K. Murakami, F. Ueno, A. Noda, T. Kobayashi, M. Kobayashi, T. Usuzaki, H. Ohseto, A. Hozawa, M. Kikuya, H. Metoki, S. Kure, S. Kuriyama, Clustering by phenotype and genome-wide association study in autism. Transl Psychiatry. 10, 290 (2020).

12. G. D. Fischbach, C. Lord, The Simons Simplex Collection: a resource for identification of autism genetic risk factors. Neuron. 68, 192–195 (2010).

13. Y. P. Raykov, A. Boukouvalas, F. Baig, M. A. Little, What to Do When K-Means Clustering Fails: A Simple yet Principled Alternative Algorithm. PLoS One. 11, e0162259 (2016).

14. P. Feliciano, A. M. Daniels, L. G. Snyder, A. Beaumont, A. Camba, A. Esler, A. G. Gulsrud, A. Mason, A. Gutierrez, A. Nicholson, A. M. Paolicelli, A. P. McKenzie, A. L. Rachubinski, A. N. Stephens, A. R. Simon, A. Stedman, A. D. Shocklee, A. Swanson, B. Finucane, B. A. Hilscher, B. Hauf, B. J. O’Roak, B. McKenna, B. E. Robertson, B. Rodriguez, B. M. Vernoia, B. V. Metre, C. Bradley, C. Cohen, C. A. Erickson, C. Harkins, C. Hayes, C. Lord, C. L. Martin, C. Ortiz, C. Ochoa-Lubinoff, C. Peura, C. E. Rice, C. R. Rosenberg, C. J. Smith, C. Thomas, C. M. Taylor, L. C. White, C. H. Walston, D. G. Amaral, D. L. Coury, D. E. Sarver, D. Istephanous, D. Li, D. C. Nugyen, E. A. Fox, E. M. Butter, E. Berry-Kravis, E. Courchesne, E. J. Fombonne, E. Hofammann, E. Lamarche, E. L. Wodka, E. T. Matthews, E. O’Connor, E. Palen, F. Miller, G. S. Dichter, G. Marzano, G. Stein, H. Hutter, H. E. Kaplan, H. Li, H. Lechniak, H. L. Schneider, H. Zaydens, I. Arriaga, J. A. Gerdts, J. F. Cubells, J. M. Cordova, J. Gunderson, J. Lillard, J. Manoharan, J. T. McCracken, J. J. Michaelson, J. Neely, J. Orobio, J. Pandey, J. Piven, J. Scherr, J. S. Sutcliffe, J. Tjernagel, J. Wallace, K. Callahan, K. Dent, K. A. Schweers, K. E. Hamer, J. K. Law, K. Lowe, K. O’Brien, K. Smith, K. G. Pawlowski, K. L. Pierce, K. Roeder, L. J. Abbeduto, L. N. Berry, L. A. Cartner, L. A. Coppola, L. Carpenter, L. Cordeiro, L. DeMarco, L. P. Grosvenor, L. Higgins, L. Y. Huang-Storms, L. Hosmer-Quint, L. M. Herbert, L. Kasparson, L. M. Prock, L. D. Pacheco, L. Raymond, L. Simon, L. V. Soorya, L. Wasserburg, M. Lazar, M. Alessandri, M. Brown, M. H. Currin, M. F. Gwynette, M. Heyman, M. N. Hale, M. Jones, M. Jordy, M. J. Morrier, M. Sahin, M. S. Siegel, M. Verdi, M. V. Parlade, M. Yinger, N. Bardett, N. Hanna, N. Harris, N. Pottschmidt, N. Russo-Ponsaran, N. Takahashi, O. Y. Ousley, A. P. Juarez, P. Manning, R. D. Annett, R. A. Bernier, R. D. Clark, R. J. Landa, R. P. Goin-Kochel, R. Remington, R. T. Schultz, S. J. Brewster, S. Booker, S. Carpenter, S. Eldred, S. Francis, S. L. Friedman, S. Horner, S. Hepburn, S. Jacob, S. Kanne, S. J. Lee, S. A. Mastel, S. Plate, S. Qiu, S. Sandhu, S. Thompson, S. White, V. J. Myers, V. Singh, W. S. Yang, Z. Warren, A. Amatya, A. J. Ace, A. S. Chatha, A. E. Lash, B. Negron, C. Rigby, C. Ridenour, C. M. Stock, D. Schmidt, I. Fisk, J. Acampado, J. L. Nestle, J. A. Nestle, K. Layman, M. E. Butler, M. Kent, M. D. Mallardi, N. Carriero, N. Lawson, N. Volfovsky, R. Edgar, R. Marini, R. Rana, S. Ganesan, S. Shah, T. Ramsey, W. Chin, W. Jensen, A. D. Krentz, A. J. Gruber, A. Sabo, A. Salomatov, C. Eng, D. Muzny, I. Astrovskaya, R. A. Gibbs, X. Han, Y. Shen, L. F. Reichardt, W. K. Chung, SPARK: A US Cohort of 50,000 Families to Accelerate Autism Research. Neuron. 97, 488–493 (2018).

15. J. Xie, R. Girshick, A. Farhadi, “Unsupervised deep embedding for clustering analysis” in International conference on machine learning (PMLR, 2016), pp. 478–487.

16. N. Rohani, C. Eslahchi, Classifying Breast Cancer Molecular Subtypes by Using Deep Clustering Approach. Front Genet. 11, 553587 (2020).

17. Y. Wang, X. Ding, Z. Tan, C. Ning, K. Xing, T. Yang, Y. Pan, D. Sun, C. Wang, Genome-Wide Association Study of Piglet Uniformity and Farrowing Interval. Front Genet. 8, 194 (2017).

18. M. M. Schoemaker, B. Flapper, N. P. Verheij, B. N. Wilson, H. A. Reinders-Messelink, A. de Kloet, Evaluation of the Developmental Coordination Disorder Questionnaire as a screening instrument. Dev Med Child Neurol. 48, 668–673 (2006).

19. K. S. L. Lam, M. G. Aman, The Repetitive Behavior Scale-Revised: independent validation in individuals with autism spectrum disorders. J Autism Dev Disord. 37, 855–866 (2007).

20. L. C. Eaves, H. D. Wingert, H. H. Ho, E. C. R. Mickelson, Screening for autism spectrum disorders with the social communication questionnaire. J Dev Behav Pediatr. 27, S95–S103 (2006).

21. C. S. Leblond, F. Cliquet, C. Carton, G. Huguet, A. Mathieu, T. Kergrohen, J. Buratti, N. Lemière, L. Cuisset, T. Bienvenu, A. Boland, J.-F. Deleuze, T. Stora, R. Biskupstoe, J. Halling, G. Andorsdóttir, E. Billstedt, C. Gillberg, T. Bourgeron, Both rare and common genetic variants contribute to autism in the Faroe Islands. NPJ Genom Med. 4, 1 (2019).

22. S.-Y. Lee, T. A. Vuong, H.-K. So, H.-J. Kim, Y. B. Kim, J.-S. Kang, I. Kwon, H. Cho, PRMT7 deficiency causes dysregulation of the HCN channels in the CA1 pyramidal cells and impairment of social behaviors. Exp Mol Med. 52, 604–614 (2020).

23. S. Zare, F. Mashayekhi, E. Bidabadi, The association of CNTNAP2 rs7794745 gene polymorphism and autism in Iranian population. J Clin Neurosci. 39, 189–192 (2017).

24. Y. Tao, H. Gao, B. Ackerman, W. Guo, D. Saffen, Y. Y. Shugart, Evidence for contribution of common genetic variants within chromosome 8p21.2-8p21.1 to restricted and repetitive behaviors in autism spectrum disorders. BMC Genomics. 17, 163 (2016).

25. H. M. Ozgen, W. G. Staal, J. C. Barber, M. V. de Jonge, M. J. Eleveld, F. A. Beemer, R. Hochstenbach, M. Poot, A novel 6.14 Mb duplication of chromosome 8p21 in a patient with autism and self mutilation. J Autism Dev Disord. 39, 322–329 (2009).

26. X. Liu, T. Shimada, T. Otowa, Y.-Y. Wu, Y. Kawamura, M. Tochigi, Y. Iwata, T. Umekage, T. Toyota, M. Maekawa, Y. Iwayama, K. Suzuki, C. Kakiuchi, H. Kuwabara, Y. Kano, H. Nishida, T. Sugiyama, N. Kato, C.-H. Chen, N. Mori, K. Yamada, T. Yoshikawa, K. Kasai, K. Tokunaga, T. Sasaki, S. S.-F. Gau, Genome-wide Association Study of Autism Spectrum Disorder in the East Asian Populations. Autism Res. 9, 340–349 (2016).

27. J. L. Roberts, K. Hovanes, M. Dasouki, A. M. Manzardo, M. G. Butler, Chromosomal microarray analysis of consecutive individuals with autism spectrum disorders or learning disability presenting for genetic services. Gene. 535, 70–78 (2014).

28. H. K. Harris, T. Nakayama, J. Lai, B. Zhao, N. Argyrou, C. S. Gubbels, A. Soucy, C. A. Genetti, V. Suslovitch, L. H. Rodan, G. E. Tiller, G. Lesca, K. W. Gripp, R. Asadollahi, A. Hamosh, C. D. Applegate, P. D. Turnpenny, M. E. H. Simon, C. M. L. Volker-Touw, K. L. I. van Gassen, E. van Binsbergen, R. Pfundt, T. Gardeitchik, B. B. A. de Vries, L. L. Immken, C. Buchanan, M. Willing, T. L. Toler, E. Fassi, L. Baker, F. Vansenne, X. Wang, J. L. Ambrus, M. Fannemel, J. E. Posey, E. Agolini, A. Novelli, A. Rauch, P. Boonsawat, C. R. Fagerberg, M. J. Larsen, M. Kibaek, A. Labalme, A. Poisson, K. K. Payne, L. E. Walsh, K. A. Aldinger, J. Balciuniene, C. Skraban, C. Gray, J. Murrell, C. P. Bupp, G. Pascolini, P. Grammatico, M. Broly, S. Küry, M. Nizon, I. G. Rasool, M. Y. Zahoor, C. Kraus, A. Reis, M. Iqbal, K. Uguen, S. Audebert-Bellanger, C. Ferec, S. Redon, J. Baker, Y. Wu, G. Zampino, S. Syrbe, I. Brosse, R. A. Jamra, W. B. Dobyns, L. L. Cohen, A. Blomhoff, C. Mignot, B. Keren, T. Courtin, P. B. Agrawal, A. H. Beggs, T. W. Yu, Disruption of RFX family transcription factors causes autism, attention-deficit/hyperactivity disorder, intellectual disability, and dysregulated behavior. Genet Med. 23, 1028–1040 (2021).

29. H.-G. Bernstein, G. Keilhoff, H. Dobrowolny, U. Lendeckel, J. Steiner, From putative brain tumor marker to high cognitive abilities: Emerging roles of a disintegrin and metalloprotease (ADAM) 12 in the brain. J Chem Neuroanat. 109, 101846 (2020).

30. H. A. F. Stessman, B. Xiong, B. P. Coe, T. Wang, K. Hoekzema, M. Fenckova, M. Kvarnung, J. Gerdts, S. Trinh, N. Cosemans, L. Vives, J. Lin, T. N. Turner, G. Santen, C. Ruivenkamp, M. Kriek, A. van Haeringen, E. Aten, K. Friend, J. Liebelt, C. Barnett, E. Haan, M. Shaw, J. Gecz, B.-M. Anderlid, A. Nordgren, A. Lindstrand, C. Schwartz, R. F. Kooy, G. Vandeweyer, C. Helsmoortel, C. Romano, A. Alberti, M. Vinci, E. Avola, S. Giusto, E. Courchesne, T. Pramparo, K. Pierce, S. Nalabolu, D. G. Amaral, I. E. Scheffer, M. B. Delatycki, P. J. Lockhart, F. Hormozdiari, B. Harich, A. Castells-Nobau, K. Xia, H. Peeters, M. Nordenskjöld, A. Schenck, R. A. Bernier, E. E. Eichler, Targeted sequencing identifies 91 neurodevelopmental-disorder risk genes with autism and developmental-disability biases. Nat Genet. 49, 515–526 (2017).

31. J. Finik, J. Buthmann, W. Zhang, K. Go, Y. Nomura, Placental Gene Expression and Offspring Temperament Trajectories: Predicting Negative Affect in Early Childhood. J Abnorm Child Psychol. 48, 783–795 (2020).

32. E. Mick, A. Todorov, S. Smalley, X. Hu, S. Loo, R. D. Todd, J. Biederman, D. Byrne, B. Dechairo, A. Guiney, J. McCracken, J. McGough, S. F. Nelson, A. M. Reiersen, T. E. Wilens, J. Wozniak, B. M. Neale, S. V. Faraone, Family-Based Genome-Wide Association Scan of Attention-Deficit/Hyperactivity Disorder. J Am Acad Child Adolesc Psychiatry. 49, 898–905.e3 (2010).

33. Y. Zhang, W. Kong, Y. Gao, X. Liu, K. Gao, H. Xie, Y. Wu, Y. Zhang, J. Wang, F. Gao, X. Wu, Y. Jiang, Gene Mutation Analysis in 253 Chinese Children with Unexplained Epilepsy and Intellectual/Developmental Disabilities. PLoS One. 10, e0141782 (2015).

34. C. Marini, A. Porro, A. Rastetter, C. Dalle, I. Rivolta, D. Bauer, R. Oegema, C. Nava, E. Parrini, D. Mei, C. Mercer, R. Dhamija, C. Chambers, C. Coubes, J. Thévenon, P. Kuentz, S. Julia, L. Pasquier, C. Dubourg, W. Carré, A. Rosati, F. Melani, T. Pisano, M. Giardino, A. M. Innes, Y. Alembik, S. Scheidecker, M. Santos, S. Figueiroa, C. Garrido, C. Fusco, D. Frattini, C. Spagnoli, A. Binda, T. Granata, F. Ragona, E. Freri, S. Franceschetti, L. Canafoglia, B. Castellotti, C. Gellera, R. Milanesi, M. M. Mancardi, D. R. Clark, F. Kok, K. L. Helbig, S. Ichikawa, L. Sadler, J. Neupauerová, P. Laššuthova, K. Šterbová, A. Laridon, E. Brilstra, B. Koeleman, J. R. Lemke, F. Zara, P. Striano, J. Soblet, G. Smits, N. Deconinck, A. Barbuti, D. DiFrancesco, E. LeGuern, R. Guerrini, B. Santoro, K. Hamacher, G. Thiel, A. Moroni, J. C. DiFrancesco, C. Depienne, HCN1 mutation spectrum: from neonatal epileptic encephalopathy to benign generalized epilepsy and beyond. Brain. 141, 3160–3178 (2018).

35. S. Mirzajani, S. Ghafouri-Fard, J. M. Habibabadi, S. Arsang-Jang, M. D. Omrani, S. S. H. Fesharaki, A. Sayad, M. Taheri, Expression Analysis of lncRNAs in Refractory and Non-Refractory Epileptic Patients. J Mol Neurosci. 70, 689–698 (2020).

36. G. L. Cavalleri, M. E. Weale, K. V. Shianna, R. Singh, J. M. Lynch, B. Grinton, C. Szoeke, K. Murphy, P. Kinirons, D. O’Rourke, D. Ge, C. Depondt, K. G. Claeys, M. Pandolfo, C. Gumbs, N. Walley, J. McNamara, J. C. Mulley, K. N. Linney, L. J. Sheffield, R. A. Radtke, S. K. Tate, S. L. Chissoe, R. A. Gibson, D. Hosford, A. Stanton, T. D. Graves, M. G. Hanna, K. Eriksson, A.-M. Kantanen, R. Kalviainen, T. J. O’Brien, J. W. Sander, J. S. Duncan, I. E. Scheffer, S. F. Berkovic, N. W. Wood, C. P. Doherty, N. Delanty, S. M. Sisodiya, D. B. Goldstein, Multicentre search for genetic susceptibility loci in sporadic epilepsy syndrome and seizure types: a case-control study. Lancet Neurol. 6, 970–980 (2007).

37. I. Cajigas, A. Chakraborty, K. R. Swyter, H. Luo, M. Bastidas, M. Nigro, E. R. Morris, S. Chen, M. J. W. VanGompel, D. Leib, S. J. Kohtz, M. Martina, S. Koh, F. Ay, J. D. Kohtz, The Evf2 Ultraconserved Enhancer lncRNA Functionally and Spatially Organizes Megabase Distant Genes in the Developing Forebrain. Mol Cell. 71, 956–972.e9 (2018).

38. S. Nadalin, A. Buretić-Tomljanović, An association between the BanI polymorphism of the PLA2G4A gene for calcium-dependent phospholipase A2 and plasma glucose levels among females with schizophrenia. Prostaglandins Leukot Essent Fatty Acids. 135, 39–41 (2018).

39. Y. Liu, X. Fu, Z. Tang, C. Li, Y. Xu, F. Zhang, D. Zhou, C. Zhu, Altered expression of the CSMD1 gene in the peripheral blood of schizophrenia patients. BMC Psychiatry. 19, 113 (2019).

40. V. M. Steen, C. Nepal, K. M. Ersland, R. Holdhus, M. Nævdal, S. M. Ratvik, S. Skrede, B. Håvik, Neuropsychological deficits in mice depleted of the schizophrenia susceptibility gene CSMD1. PLoS One. 8, e79501 (2013).

41. J. A. Azevedo, B. S. Carter, F. Meng, D. L. Turner, M. Dai, A. F. Schatzberg, J. D. Barchas, E. G. Jones, W. E. Bunney, R. M. Myers, H. Akil, S. J. Watson, R. C. Thompson, The microRNA network is altered in anterior cingulate cortex of patients with unipolar and bipolar depression. J Psychiatr Res. 82, 58–67 (2016).

42. S. Huang, C. Zheng, G. Xie, Z. Song, P. Wang, Y. Bai, D. Chen, Y. Zhang, P. Lv, W. Liang, S. She, Q. Li, Z. Liu, Y. Wang, G.-G. Xing, Y. Wang, FAM19A5/TAFA5, a novel neurokine, plays a crucial role in depressive-like and spatial memory-related behaviors in mice. Mol Psychiatry. 26, 2363–2379 (2021).

43. X. Chang, J. Wang, H. Jiang, L. Shi, J. Xie, Hyperpolarization-Activated Cyclic Nucleotide-Gated Channels: An Emerging Role in Neurodegenerative Diseases. Front Mol Neurosci. 12, 141 (2019).

44. Y. Hu, L. Deng, J. Zhang, X. Fang, P. Mei, X. Cao, J. Lin, Y. Wei, X. Zhang, R. Xu, A Pooling Genome-Wide Association Study Combining a Pathway Analysis for Typical Sporadic Parkinson’s Disease in the Han Population of Chinese Mainland. Mol Neurobiol. 53, 4302–4318 (2016).

45. M. D. Muñoz, N. de la Fuente, A. Sánchez-Capelo, TGF-β/Smad3 Signalling Modulates GABA Neurotransmission: Implications in Parkinson’s Disease. Int J Mol Sci. 21, E590 (2020).

46. J. Ruiz-Martínez, L. J. Azcona, A. Bergareche, J. F. Martí-Massó, C. Paisán-Ruiz, Whole-exome sequencing associates novel CSMD1 gene mutations with familial Parkinson disease. Neurol Genet. 3, e177 (2017).

47. Y. J. Li, J. Deng, G. M. Mayhew, J. W. Grimsley, X. Huo, J. M. Vance, Investigation of the PARK10 gene in Parkinson disease. Ann Hum Genet. 71, 639–647 (2007).

48. S. A. Oliveira, Y.-J. Li, M. A. Noureddine, S. Zuchner, X. Qin, M. A. Pericak-Vance, J. M. Vance, Identification of risk and age-at-onset genes on chromosome 1p in Parkinson disease. Am J Hum Genet. 77, 252–264 (2005).

49. P. S. Eis, N. Huang, J. W. Langston, E. Hatchwell, B. Schüle, Loss-of-Function NUBPL Mutation May Link Parkinson’s Disease to Recessive Complex I Deficiency. Front Neurol. 11, 555961 (2020).

50. T. Müller, C. Loosse, A. Schrötter, A. Schnabel, S. Helling, R. Egensperger, K. Marcus, The AICD interacting protein DAB1 is up-regulated in Alzheimer frontal cortex brain samples and causes deregulation of proteins involved in gene expression changes. Curr Alzheimer Res. 8, 573–582 (2011).

51. H. N. Cukier, B. K. Kunkle, K. L. Hamilton, S. Rolati, M. A. Kohli, P. L. Whitehead, J. Jaworski, J. M. Vance, M. L. Cuccaro, R. M. Carney, J. R. Gilbert, L. A. Farrer, E. R. Martin, G. W. Beecham, J. L. Haines, M. A. Pericak-Vance, Exome Sequencing of Extended Families with Alzheimer’s Disease Identifies Novel Genes Implicated in Cell Immunity and Neuronal Function. J Alzheimers Dis Parkinsonism. 7, 355 (2017).

52. G. A. Pathak, F. R. Wendt, A. De Lillo, Y. Z. Nunez, A. Goswami, F. De Angelis, M. Fuciarelli, H. R. Kranzler, J. Gelernter, R. Polimanti, Epigenomic Profiles of African-American Transthyretin Val122Ile Carriers Reveals Putatively Dysregulated Amyloid Mechanisms. Circ Genom Precis Med. 14, e003011 (2021).

53. A. Parcerisas, S. E. Rubio, A. Muhaisen, A. Gómez-Ramos, L. Pujadas, M. Puiggros, D. Rossi, J. Ureña, F. Burgaya, M. Pascual, D. Torrents, A. Rábano, J. Avila, E. Soriano, Somatic signature of brain-specific single nucleotide variations in sporadic Alzheimer’s disease. J Alzheimers Dis. 42, 1357–1382 (2014).

54. R. A. K. Srivastava, J. C. Jain, Scavenger receptor class B type I expression and elemental analysis in cerebellum and parietal cortex regions of the Alzheimer’s disease brain. J Neurol Sci. 196, 45–52 (2002).

55. A. Raina, D. Kaul, LXR-α genomics programmes neuronal death observed in Alzheimer’s disease. Apoptosis. 15, 1461–1469 (2010).

56. S. Martins, C. E. Pearson, P. Coutinho, S. Provost, A. Amorim, M.-P. Dubé, J. Sequeiros, G. A. Rouleau, Modifiers of (CAG)(n) instability in Machado-Joseph disease (MJD/SCA3) transmissions: an association study with DNA replication, repair and recombination genes. Hum Genet. 133, 1311–1318 (2014).

57. B. Xie, X. Fan, Y. Lei, R. Chen, J. Wang, C. Fu, S. Yi, J. Luo, S. Zhang, Q. Yang, S. Chen, Y. Shen, A novel de novo microdeletion at 17q11.2 adjacent to NF1 gene associated with developmental delay, short stature, microcephaly and dysmorphic features. Mol Cytogenet. 9, 41 (2016).

58. G. Y. Zheleznyakova, E. K. Nilsson, A. V. Kiselev, M. A. Maretina, L. I. Tishchenko, R. Fredriksson, V. S. Baranov, H. B. Schiöth, Methylation levels of SLC23A2 and NCOR2 genes correlate with spinal muscular atrophy severity. PLoS One. 10, e0121964 (2015).

59. H. Pantera, M. E. Shy, J. Svaren, Regulating PMP22 expression as a dosage sensitive neuropathy gene. Brain Res. 1726, 146491 (2020).

60. J. M. Lane, J. Liang, I. Vlasac, S. G. Anderson, D. A. Bechtold, J. Bowden, R. Emsley, S. Gill, M. A. Little, A. I. Luik, A. Loudon, F. A. J. L. Scheer, S. M. Purcell, S. D. Kyle, D. A. Lawlor, X. Zhu, S. Redline, D. W. Ray, M. K. Rutter, R. Saxena, Genome-wide association analyses of sleep disturbance traits identify new loci and highlight shared genetics with neuropsychiatric and metabolic traits. Nat Genet. 49, 274–281 (2017).

61. L. Ni, Y. Xu, S. Dong, Y. Kong, H. Wang, G. Lu, Y. Wang, Q. Li, C. Li, Z. Du, H. Sun, L. Sun, The potential role of the HCN1 ion channel and BDNF-mTOR signaling pathways and synaptic transmission in the alleviation of PTSD. Transl Psychiatry. 10, 101 (2020).

62. P. Tosetti, N. Pathak, M. H. Jacob, K. Dunlap, RGS3 mediates a calcium-dependent termination of G protein signaling in sensory neurons. Proc Natl Acad Sci U S A. 100, 7337–7342 (2003).

63. V. Kimonis, R. Al Dubaisi, A. E. Maclean, K. Hall, L. Weiss, A. E. Stover, P. H. Schwartz, B. Berg, C. Cheng, S. Parikh, B. R. Conner, S. Wu, A. N. Hasso, D. A. Scott, M. K. Koenig, R. Karam, S. Tang, M. Smith, E. Chao, J. Balk, E. Hatchwell, P. S. Eis, NUBPL mitochondrial disease: new patients and review of the genetic and clinical spectrum. J Med Genet. 58, 314–325 (2021).

64. J. B. Schuch, V. R. Paixão-Côrtes, D. Longo, T. Roman, R. D. S. Riesgo, J. Ranzan, M. M. Becker, M. Riegel, L. Schuler-Faccini, Analysis of a Protein Network Related to Copy Number Variations in Autism Spectrum Disorder. J Mol Neurosci. 69, 140–149 (2019).

65. Cancer Genome Atlas Research Network, The Molecular Taxonomy of Primary Prostate Cancer. Cell. 163, 1011–1025 (2015).

66. G.-Q. Qin, H.-C. He, Z.-D. Han, Y.-X. Liang, S.-B. Yang, Y.-Q. Huang, L. Zhou, H. Fu, J.-X. Li, F.-N. Jiang, W. Zhong, Combined overexpression of HIVEP3 and SOX9 predicts unfavorable biochemical recurrence-free survival in patients with prostate cancer. Onco Targets Ther. 7, 137–146 (2014).

67. J. Ham, D. Jeong, S. Park, H. W. Kim, H. Kim, S. J. Kim, Ginsenoside Rg3 and Korean Red Ginseng extract epigenetically regulate the tumor-related long noncoding RNAs RFX3-AS1 and STXBP5-AS1. J Ginseng Res. 43, 625–634 (2019).

68. D. A. Quigley, E. Fiorito, S. Nord, P. Van Loo, G. G. Alnæs, T. Fleischer, J. Tost, H. K. Moen Vollan, T. Tramm, J. Overgaard, I. R. Bukholm, A. Hurtado, A. Balmain, A.-L. Børresen-Dale, V. Kristensen, The 5p12 breast cancer susceptibility locus affects MRPS30 expression in estrogen-receptor positive tumors. Mol Oncol. 8, 273–284 (2014).

69. B. Wu, Y. Pan, G. Liu, T. Yang, Y. Jin, F. Zhou, Y. Wei, MRPS30-DT Knockdown Inhibits Breast Cancer Progression by Targeting Jab1/Cops5. Front Oncol. 9, 1170 (2019).

70. P. S. Felicio, L. T. Bidinotto, M. E. Melendez, R. S. Grasel, N. Campacci, H. C. R. Galvão, C. Scapulatempo-Neto, R. M. Dufloth, A. F. Evangelista, E. I. Palmero, Genetic alterations detected by comparative genomic hybridization in BRCAX breast and ovarian cancers of Brazilian population. Oncotarget. 9, 27525–27534 (2018).

71. P. Wang, Z. Zhang, Y. Ma, J. Lu, H. Zhao, S. Wang, J. Tan, B. Li, Prognostic values of GMPS, PR, CD40, and p21 in ovarian cancer. PeerJ. 7, e6301 (2019).

72. X.-X. Cui, C. Zhou, H. Lu, Y.-L. Han, F.-M. Wang, W.-R. Fan, Y. Ren, R. Zhang, High expression of ZNF93 promotes proliferation and migration of ovarian cancer cells and relates to poor prognosis. Int J Clin Exp Pathol. 13, 944–953 (2020).

73. Z. Chen, S. Xiong, J. Li, L. Ou, C. Li, J. Tao, Z. Jiang, J. Fan, J. He, W. Liang, DNA methylation markers that correlate with occult lymph node metastases of non-small cell lung cancer and a preliminary prediction model. Transl Lung Cancer Res. 9, 280–287 (2020).

74. D. Welcker, M. Jain, S. Khurshid, M. Jokić, M. Höhne, A. Schmitt, P. Frommolt, C. M. Niessen, J. Spiro, T. Persigehl, M. Wittersheim, R. Büttner, M. Fanciulli, B. Schermer, H. C. Reinhardt, T. Benzing, K. Höpker, AATF suppresses apoptosis, promotes proliferation and is critical for Kras-driven lung cancer. Oncogene. 37, 1503–1518 (2018).

75. Q. Ju, Y.-J. Zhao, S. Ma, X.-M. Li, H. Zhang, S.-Q. Zhang, Y.-M. Yang, S.-X. Yan, Genome-wide analysis of prognostic-related lncRNAs, miRNAs and mRNAs forming a competing endogenous RNA network in lung squamous cell carcinoma. J Cancer Res Clin Oncol. 146, 1711–1723 (2020).

76. J. H. Chi, A. Panner, K. Cachola, C. A. Crane, J. Murray, R. O. Pieper, C. D. James, A. T. Parsa, Increased expression of the glioma-associated antigen ARF4L after loss of the tumor suppressor PTEN. Laboratory investigation. J Neurosurg. 108, 299–303 (2008).

77. T. Jin, J. Zhang, G. Li, S. Li, B. Yang, C. Chen, L. Cai, TP53 and RPA3 gene variations were associated with risk of glioma in a Chinese Han population. Cancer Biother Radiopharm. 28, 248–253 (2013).

78. C. Tan, P. Scotting, Expression of Kit and Etv1 in restricted brain regions supports a brain-cell progenitor as an origin for cranial germinomas. Cancer Genet. 208, 55–61 (2015).

79. O. M. Wilkins, A. J. Titus, J. Gui, M. Eliot, R. A. Butler, E. M. Sturgis, G. Li, K. T. Kelsey, B. C. Christensen, Genome-scale identification of microRNA-related SNPs associated with risk of head and neck squamous cell carcinoma. Carcinogenesis. 38, 986–993 (2017).

80. Z. Dai, S. Wang, W. Zhang, Y. Yang, Elevated Expression of RPA3 Is Involved in Gastric Cancer Tumorigenesis and Associated with Poor Patient Survival. Dig Dis Sci. 62, 2369–2375 (2017).

81. P. G. Iyer, W. R. Taylor, M. L. Johnson, R. L. Lansing, K. A. Maixner, T. C. Yab, J. A. Simonson, M. E. Devens, S. W. Slettedahl, D. W. Mahoney, C. K. Berger, P. H. Foote, T. C. Smyrk, K. K. Wang, H. C. Wolfsen, D. A. Ahlquist, Highly Discriminant Methylated DNA Markers for the Non-endoscopic Detection of Barrett’s Esophagus. Am J Gastroenterol. 113, 1156–1166 (2018).

82. J. N. Crawley, W.-D. Heyer, J. M. LaSalle, Autism and cancer share risk genes, pathways and drug targets. Trends Genet. 32, 139–146 (2016).

83. A. P. Gabrielli, A. M. Manzardo, M. G. Butler, GeneAnalytics Pathways and Profiling of Shared Autism and Cancer Genes. Int J Mol Sci. 20, E1166 (2019).

84. A. Beggiato, H. Peyre, A. Maruani, I. Scheid, M. Rastam, F. Amsellem, C. I. Gillberg, M. Leboyer, T. Bourgeron, C. Gillberg, R. Delorme, Gender differences in autism spectrum disorders: Divergence among specific core symptoms. Autism Res. 10, 680–689 (2017).

85. S. Kuriyama, M. Kamiyama, M. Watanabe, S. Tamahashi, I. Muraguchi, T. Watanabe, A. Hozawa, T. Ohkubo, Y. Nishino, Y. Tsubono, I. Tsuji, S. Hisamichi, Pyridoxine treatment in a subgroup of children with pervasive developmental disorders. Dev Med Child Neurol. 44, 284–286 (2002).

86. T. Obara, M. Ishikuro, G. Tamiya, M. Ueki, C. Yamanaka, S. Mizuno, M. Kikuya, H. Metoki, H. Matsubara, M. Nagai, T. Kobayashi, M. Kamiyama, M. Watanabe, K. Kakuta, M. Ouchi, A. Kurihara, N. Fukuchi, A. Yasuhara, M. Inagaki, M. Kaga, S. Kure, S. Kuriyama, Potential identification of vitamin B6 responsiveness in autism spectrum disorder utilizing phenotype variables and machine learning methods. Sci Rep. 8, 14840 (2018).

87. World Medical Association, World Medical Association Declaration of Helsinki: ethical principles for medical research involving human subjects. JAMA. 310, 2191–2194 (2013).

88. A. M. Daniels, R. E. Rosenberg, C. Anderson, J. K. Law, A. R. Marvin, P. A. Law, Verification of parent-report of child autism spectrum disorder diagnosis to a web-based autism registry. J Autism Dev Disord. 42, 257–265 (2012).

89. N. Patterson, A. L. Price, D. Reich, Population Structure and Eigenanalysis. PLOS Genetics. 2, e190 (2006).

90. J. M. Nam, A simple approximation for calculating sample sizes for detecting linear trend in proportions. Biometrics. 43, 701–705 (1987).

91. D. R. Cutting, D. R. Karger, J. O. Pedersen, J. W. Tukey, “Scatter/Gather: a cluster-based approach to browsing large document collections” in Proceedings of the 15th annual international ACM SIGIR conference on Research and development in information retrieval (Association for Computing Machinery, New York, NY, USA, 1992; https://doi.org/10.1145/133160.133214), SIGIR ’92, pp. 318–329.

92. G. Guo, L. Chen, Y. Ye, Q. Jiang, Cluster Validation Method for Determining the Number of Clusters in Categorical Sequences. IEEE Transactions on Neural Networks and Learning Systems. 28, 2936–2948 (2017).

93. S. Purcell, B. Neale, K. Todd-Brown, L. Thomas, M. A. R. Ferreira, D. Bender, J. Maller, P. Sklar, P. I. W. de Bakker, M. J. Daly, P. C. Sham, PLINK: a tool set for whole-genome association and population-based linkage analyses. Am J Hum Genet. 81, 559–575 (2007).

94. K. Wang, M. Li, H. Hakonarson, ANNOVAR: functional annotation of genetic variants from high-throughput sequencing data. Nucleic Acids Res. 38, e164 (2010).

95. R Core Team, “R: A language and environment for statistical computing” (manual, Vienna, Austria, 2021), (available at https://www.R-project.org/).

