## Supplementary Figure1 for "Deep embedded clustering by relevant scales and genome-wide association study in autism"

### (a) Case=300,000 Control=300,000

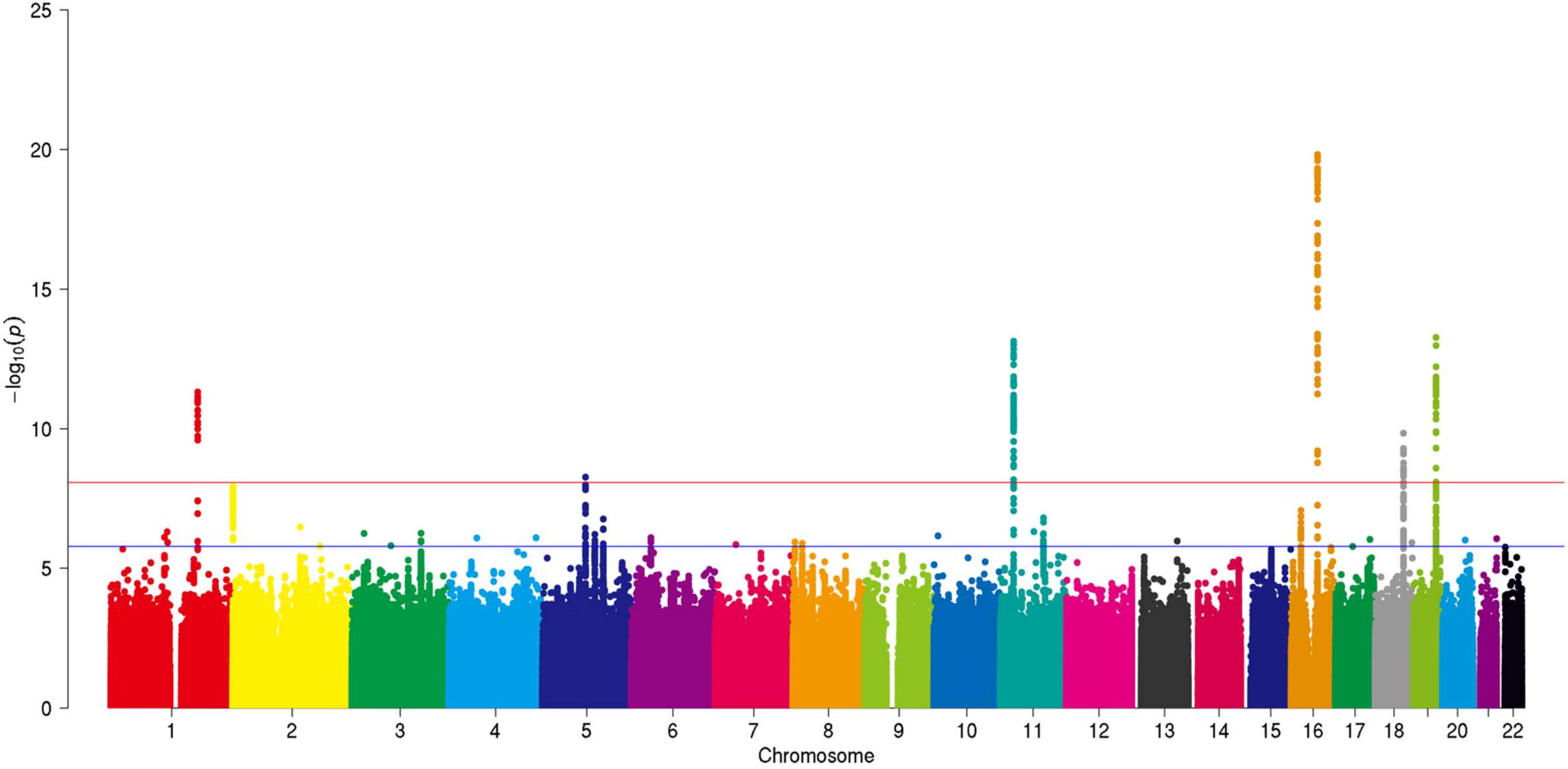


**Supplementary Figure 1.** Virtual Manhattan plots with different number of participants.

## (b) Case=30,000   Control=30,000

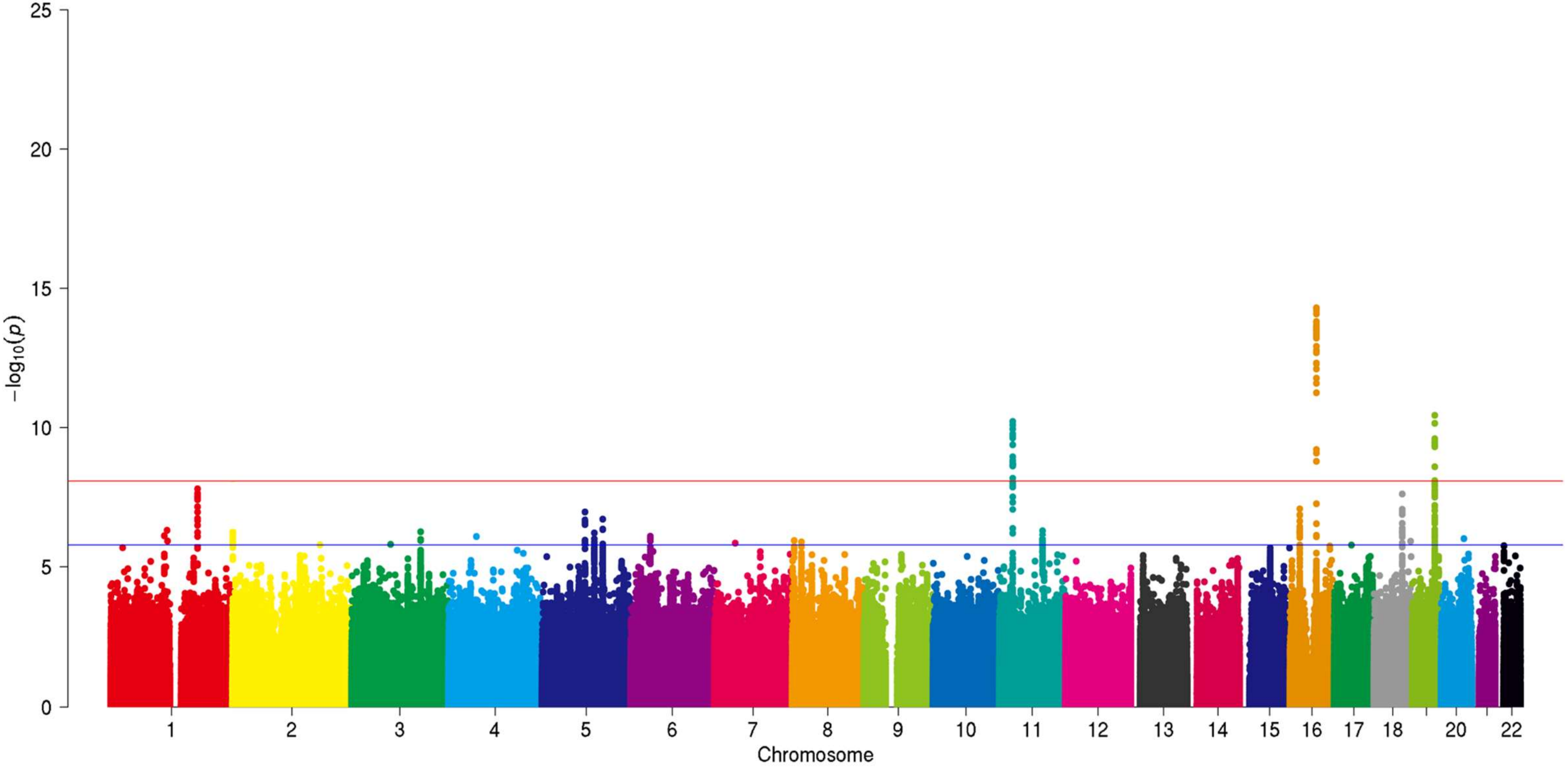

# (c) Case=3,000   Control=3,000

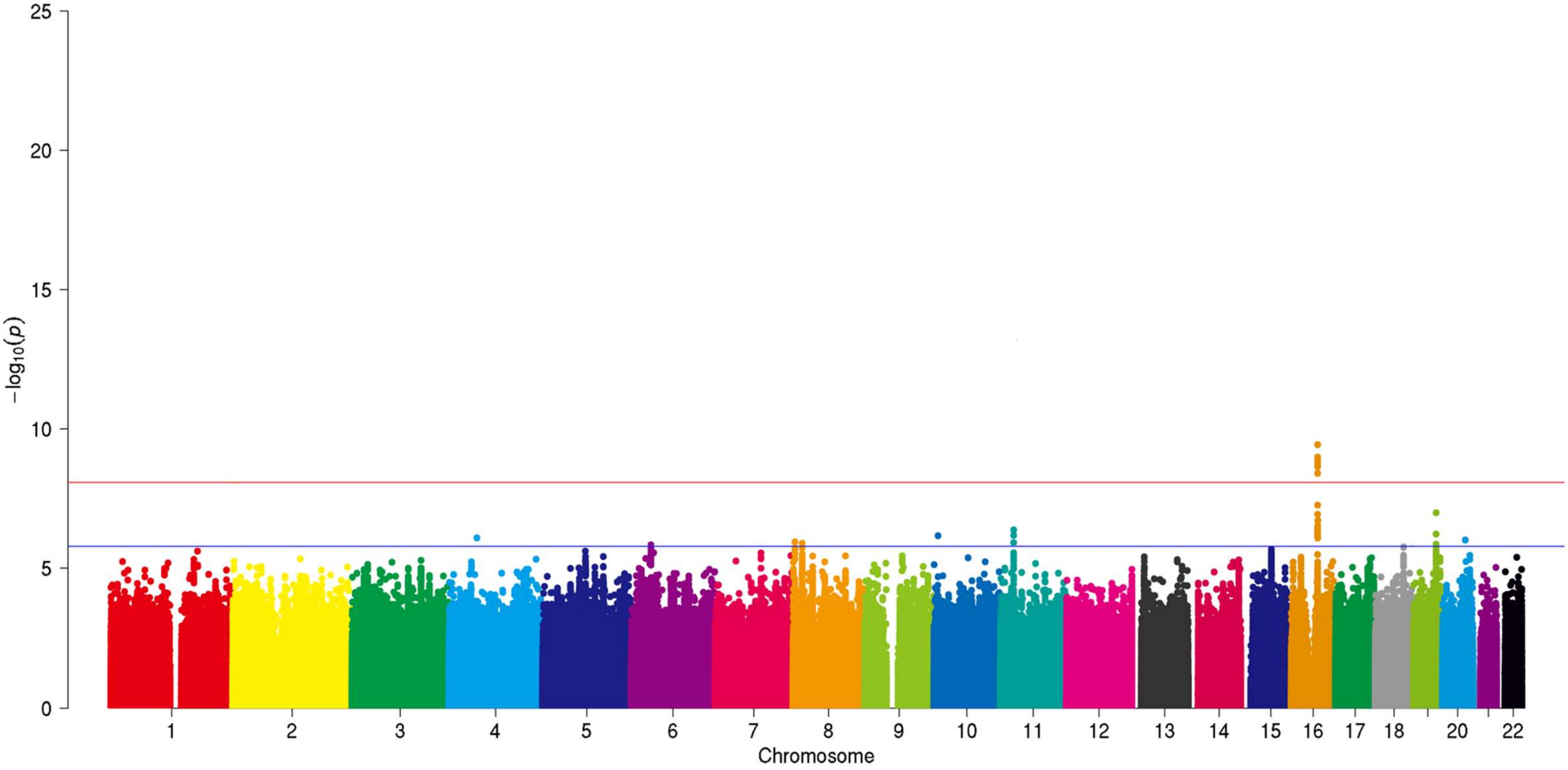

### (d) Case=300 Control=300

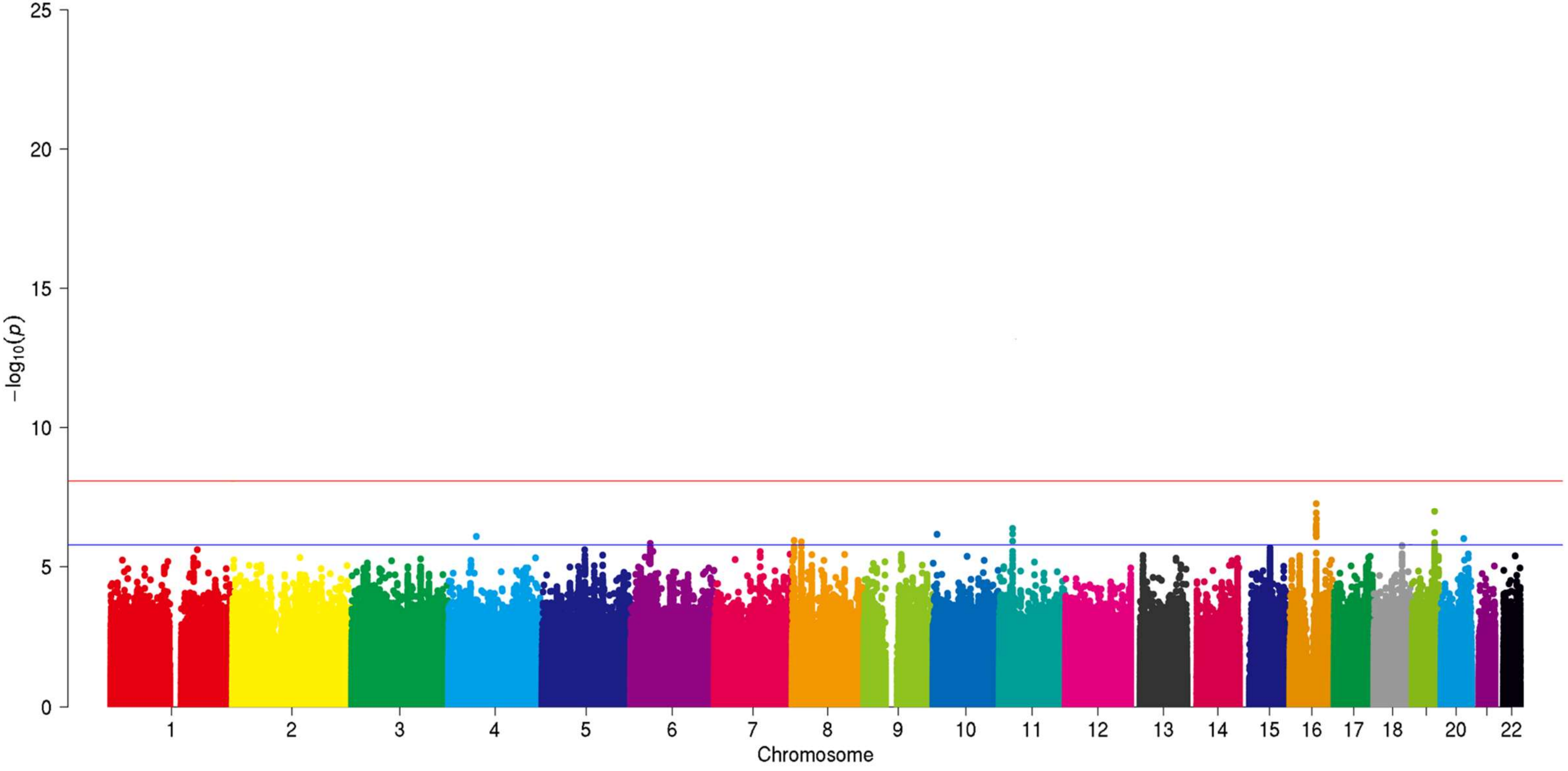
