## Supplementary figures and images for "Deep embedded clustering by relevant scales and genome-wide association study in autism"

### Supplementary Figure2

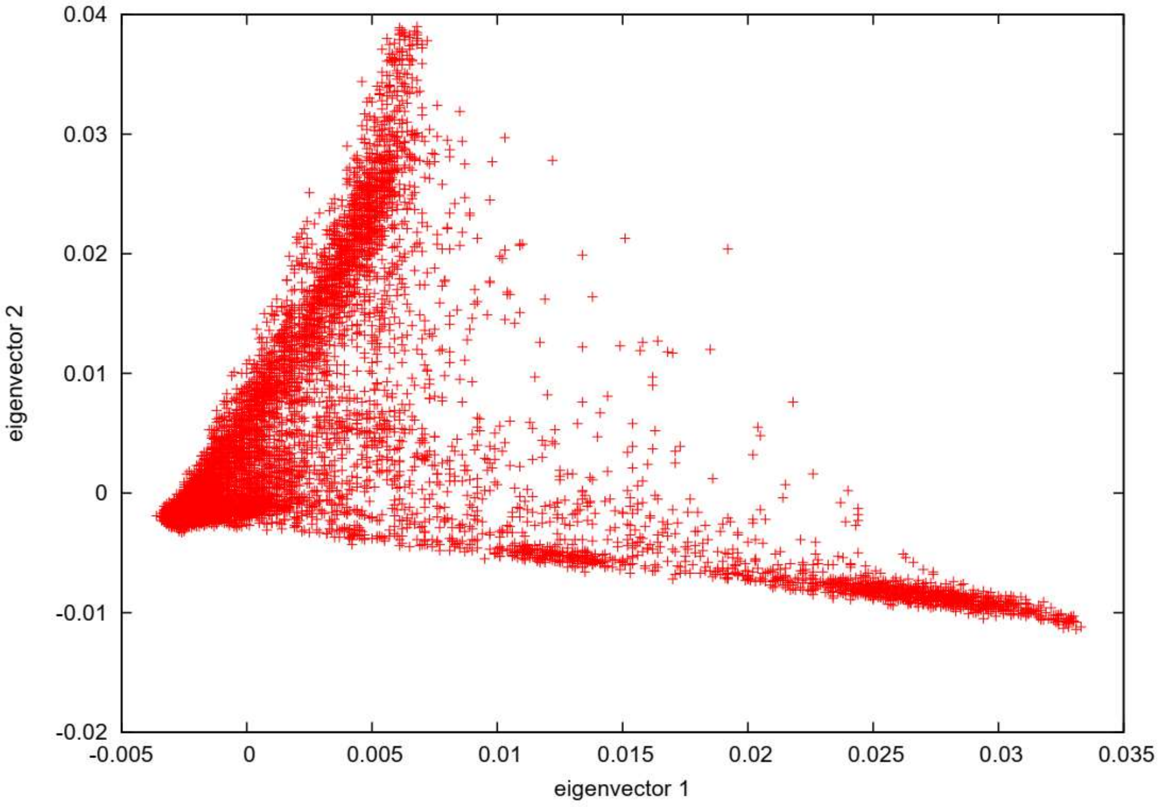


**Supplementary Figure 2.** Principal component analysis of the genotyping data.

### Supplementary Figure3

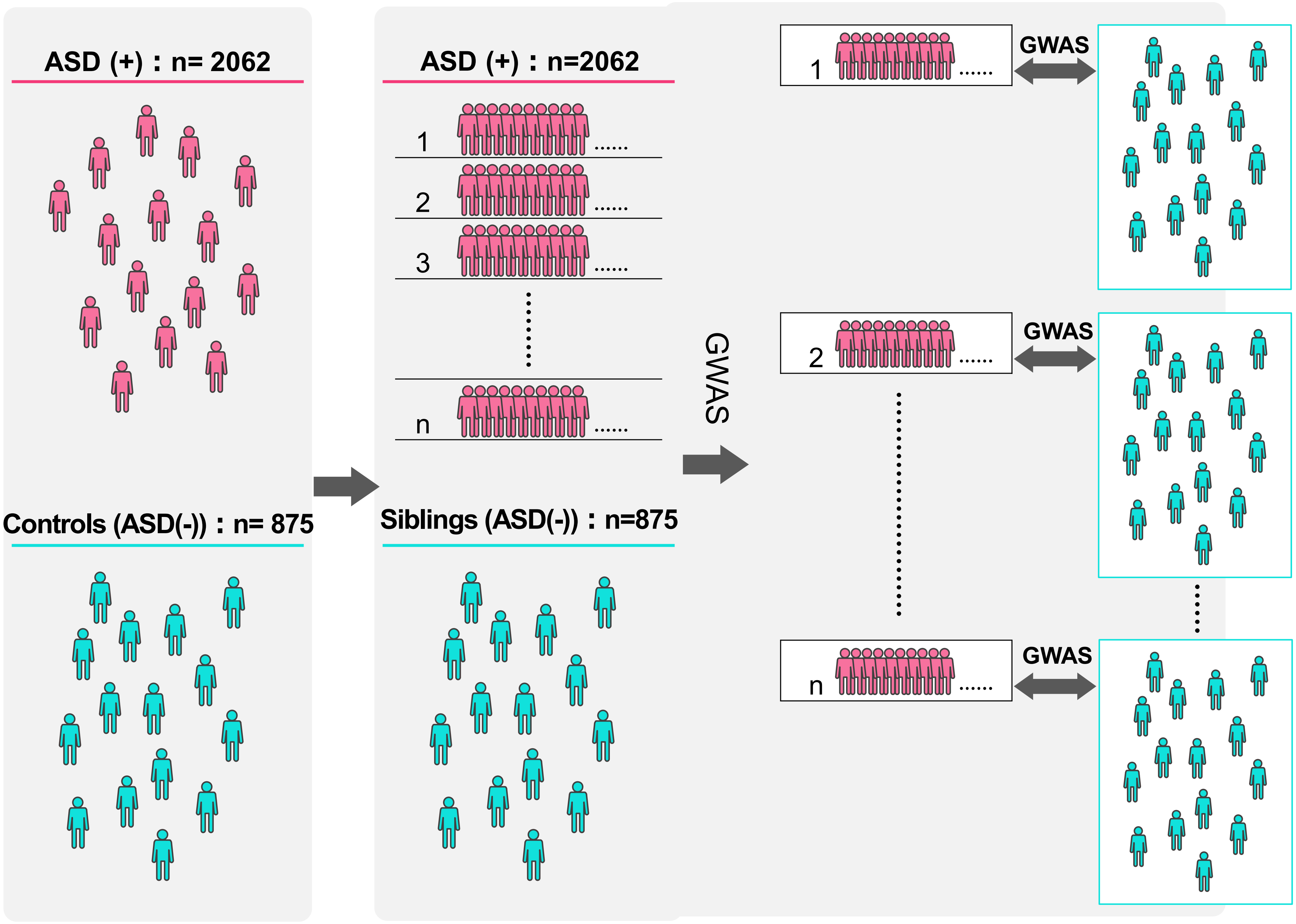


**Supplementary Figure 3.** Details of the cluster-based GWAS.
